# Altered NRG1/ErbB4 signaling and cholesterol metabolism dysregulation are key pathomechanisms in VRK1-related motor neuropathies and motor neuron diseases

**DOI:** 10.1101/2024.02.19.581053

**Authors:** Zeinab Hamzé, Camille Humbert, Khalil Rihan, Nathalie Roeckel-Trévisiol, Christel Castro, Nicolas Lenfant, Catherine Aubert, Karine Bertaux, Rosette Jabbour, Nathalie Bernard-Marissal, André Mégarbané, Valérie Delague

**Affiliations:** Aix Marseille Univ, INSERM, MMG, U 1251, Marseille, France; Centre de Ressources Biologiques, Assistance Publique Hôpitaux de Marseille, CRB APHM, Hôpital Timone Adulte, Biogénopôle, Marseille, France; Neurology Division, Department of Internal Medicine, St George Hospital University Medical Center, University of Balamand, Beirut, Lebanon; Department of Human Genetics, Gilbert and Rose-Marie Chagoury School of Medicine, Lebanese American University, Lebanon

**Keywords:** VRK1, Charcot-Marie-Tooth, motor neuropathy, Neuregulin1/ErbB4 pathway, cholesterol metabolism

## Abstract

Hereditary Motor and Sensory Neuropathy (HMSN), or Charcot-Marie-Tooth disease (CMT), are the most common group of Inherited peripheral neuropathies (IPN), characterized by a strong clinical and genetic heterogeneity. Among them, distal Hereditary Motor Neuropathy (dHMN), also known as neuronopathy, is a subgroup, where only motor nerves are affected. This subgroup is also genetically heterogeneous, with 25 genes described to date, of which VRK1, that we have recently described as responsible for dHMN, associated to upper motor neuron signs. There are now more than thirty mutations in VRK1, which cause a range of neurological diseases affecting motor neurons (mainly lower, but also upper) or their axons in the peripheral nervous system, that we design as VRK1-related motor neuron diseases.

In two previous studies, we have demonstrated that dHMN due to VRK1 mutations lead to reduced levels of VRK1 in the nucleus, and that this depletion alters the dynamics of coilin, a phosphorylation target of VRK1. hiPSC-derived Motor Neurons (hiPSC-MN) from these patients, display Cajal Bodies (CBs) disassembly and defects in neurite outgrowth and branching, altered Action Potential (AP) waveform and decreased Axonal Initial Segment (AIS) length.

In this study, we have further studied the link between the loss of VRK1 function and the defects observed in hiPSC-MNs, by realizing bulk mRNA-Seq sequencing in this in vitro model of the disease. Our results evidenced altered NRG1/ERBB4 signaling, leading to cholesterol metabolism dysregulation and deregulation of genes encoding the glutamate receptors AMPAR and NMDAR, which role in the Axonal Initial Segment and abnormal AP initiation in hiPSC-MNs remains to be investigated.

## Introduction

Hereditary Motor and Sensory Neuropathy (HMSN), or Charcot-Marie-Tooth disease (CMT), are the most common group of Inherited peripheral neuropathies (IPN), with an overall prevalence of 1/2500 (Skre, 1974). These diseases are characterized by length-dependent progressive degeneration of the Peripheral Nervous System (PNS) and extensive phenotypic and genetic heterogeneity, with alterations in over 100 genes reported to date (Laura, Pipis, Rossor, & Reilly, 2019; Pipis, Rossor, Laura, & Reilly, 2019). Although CMT is a generic term used to design Hereditary Motor and Sensory Neuropathies, there is a wide spectrum in the clinical presentations, ranging from pure motor neuropathies (HMNs) to pure sensory neuropathies (HSNs).

Patients presenting with exclusive motor neuropathy, are affected with distal Hereditary Motor Neuropathy (dHMN), also known as neuronopathy, distal Spinal Muscular Atrophy or spinal CMT (Rossor, Kalmar, Greensmith, & Reilly, 2012). Similarly to Hereditary Motor and sensory Neuropathies, the dHMN subgroup is genetically heterogeneous, with 25 genes described to date, of which VRK1, that we have recently described as responsible for dHMN, associated to upper motor neuron signs. Indeed, we have described two novel compound heterozygous missense mutations in the *VRK1* gene, in two siblings, from a non-consanguineous Lebanese family, affected with a distal hereditary motor neuropathy (dHMN), associated with upper motor neuron signs (El-Bazzal et al., 2019). Diseases due to mutations in *VRK1*, that we regrouped under the name of “ VRK1-related motor neuron diseases”, illustrate very well the clinical overlap which exists between CMT/dHMN and diseases affecting upper and/or lower motor neuron diseases, such as hereditary spastic paraplegias, Amyotrophic Lateral Sclerosis and Spinal Muscular Atrophy (Hanemann & Ludolph, 2002; Irobi et al., 2006; Martin, Hicks, Holbrook, & Cox, 2020; Rossor, Polke, Houlden, & Reilly, 2013; Timmerman, Clowes, & Reid, 2013).

*VRK1* encodes the Vaccinia Related kinase 1, a ubiquitously expressed, serine/threonine kinase, playing a crucial role in regulating cell cycle (Valbuena, Lopez-Sanchez, & Lazo, 2008; Valbuena, Sanz-Garcia, Lopez-Sanchez, Vega, & Lazo, 2011). VRK1 is mainly a nuclear protein, although the presence of a small fraction in the cytoplasm and membrane compartments has been described (Nichols & Traktman, 2004; Valbuena et al., 2007).

Several substrates are known to date for VRK1, including VRK1(Lopez-Borges & Lazo, 2000), the transcription factors p53(Belin et al., 2019), c-jun (Sevilla, Santos, Barcia, Vega, & Lazo, 2004), ATF2 (Sevilla, Santos, Vega, & Lazo, 2004) and CREB (Kang, Park, Kim, & Kim, 2008), proteins involved in DNA replication and repair, such as histone H2B (Lopez-Borges & Lazo, 2000), histone H3 (Kang et al., 2007) and NBS1 (Monsalve et al., 2016), coilin, the main component of Cajal Bodies (CBs) (Sanz-Garcia et al., 2011), myelin basic protein MBP (Lopez-Borges & Lazo, 2000), a component of central myelin and peripheral myelin, or BAF, a protein required for nuclear envelope assembly (Molitor & Traktman, 2014; Nichols, Wiebe, & Traktman, 2006).

There are now more than thirty mutations in VRK1, which cause a range of neurological diseases affecting motor neurons (mainly lower, but also upper) or their axons in the peripheral nervous system (Lazo & Morejon-Garcia, 2023), that we design as VRK1-related motor neuron diseases.

By studying patient’s cells, including induced Pluripotent Stem Cell (iPSC) derived Motor Neurons (hiPSC-MNs), we have shown that these mutations lead to reduced levels of VRK1 in the nucleus, and that this depletion alters the dynamics of coilin, a phosphorylation target of VRK1 (El-Bazzal et al., 2019). hiPSC-derived Motor Neurons (hiPSC-MN) from these patients, display Cajal Bodies (CBs) disassembly and defects in neurite outgrowth and branching, altered Action Potential (AP) waveform and decreased Axonal Initial Segment (AIS) length (Bos et al., 2022).

In this study, we have further studied the link between the loss of VRK1 function and the defects observed in hiPSC-MNs, by realizing bulk mRNA-Seq sequencing in this in vitro model of the disease. We show that the expression of the Epidermal Growth Factor Receptor ErbB4, is strongly upregulated in hiPSC-derived spinal motor neurons from our patient and that this upregulation leads to impaired Neuregulin1/ErbB4 signaling. Among the consequences of altered NRG1/ErbB4 signaling, our RNA-Seq experiments allowed us to identify strong deregulations of genes involved in cholesterol biosynthesis and metabolism, in particular genes under the control of SREBP, which are likely to induce the defects in neurite outgrowth observed in patients’ hiPSC-derived motor neurons. Finally, RNA-Seq data show upregulation of genes encoding glutamatergic receptors, probably resulting from depletion of cholesterol from membrane lipid rafts at synapses and likely to explain the abnormal initiation of the Action potential at the AIS in response to upper motor neuron stimulation. Further experiments are required to confirm these results, in hiPSC-MNs, in particular by using myotubes/motor neurons cocultures, which will help defining a more faithful biological context.

## Results

### hiPSC-MNs’ transcriptome from a patient with compound heterozygous missense mutations in *VRK1* is highly different from the controls’

In order to investigate the pathophysiological mechanisms underlying VRK1-related Motor Neuron diseases and neuropathies, we have studied the global expression of genes in hiPSC-MNs from one patient with compound heterozygous mutations in *VRK1* (Patient II.2), described in (El-Bazzal et al., 2019) and one control, in duplicate, by bulk mRNA-Seq. This investigation was initiated in particular to find the missing links between the presence of mutations in *VRK1* and the previously described defects in axonal growth and branching and in Action Potential initiation (Bos et al., 2022; El-Bazzal et al., 2019). The heatmap of unsupervised hierarchical clustering of the four samples showed that Patient II.2 clustered separately from the control (Figure 1A).

**Figure 1.**
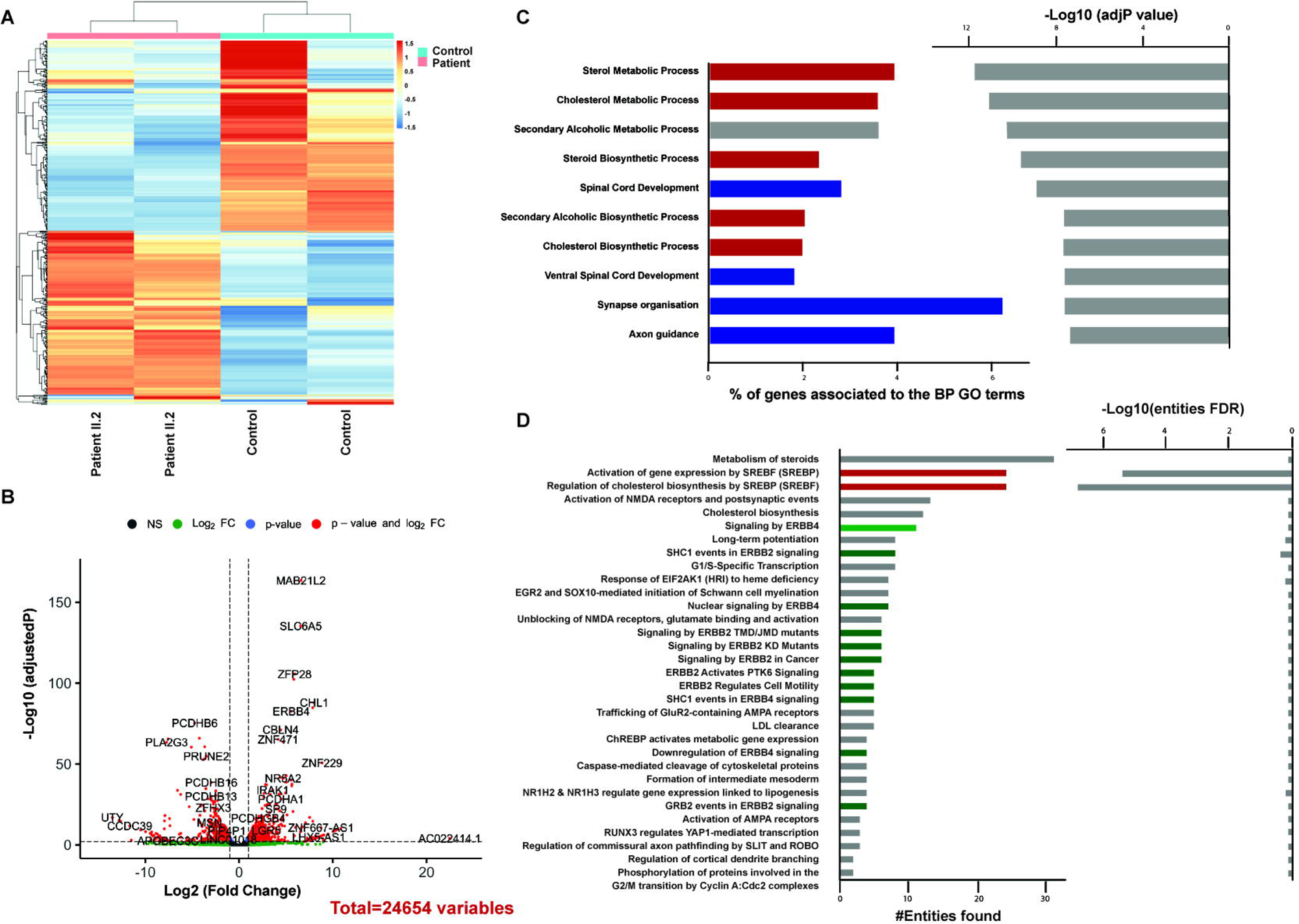
Transcriptional profiles of hiPSC-derived Motor Neurons (hiPSC-MN) from Patient II.2 and one control (hff19). (**A**) Full heat map of unsupervised hierarchical clustering of the 4 samples (n= 2 replicates for the patient and the control). The scale bar unit is obtained applying a variance stabilizing transformation (VST) to the count data (DESeq2: VarianceStabilizingTransformation) before normalization. (**B**) Volcano plot showing the distribution of gene expression fold changes and adjusted p-values between the two conditions. A total number of 24,654 genes were tested. Padj < 0.01 was used as the threshold to reject the null hypothesis and consider the difference in gene expression. Red dots represent significantly deregulated genes (adjusted p-value<0.01, with fold change (FC) >2 or <0.5). Genes with significant deregulation (adjusted p-value<0.01) but with small FC (0.5<FC<2) are indicated in blue. Green and gray dots represent genes with non-significant deregulations (adjusted p>0.01). (**C**) Top 10 Biological Process (BP) Gene Ontology (GO) terms enriched in the list of the significant DEGs (padj<0.01, FC>2 or <0.5), which are unique to the patient II.2. The bars on the left represent the percentage of DEGs determined for each represented GO term from the total number of DEGs. GO terms related to cholesterol metabolism and axonal function are highlighted in red and blue respectively. Light gray bars on the right represent the enrichment score (-Log10 of adjusted p-value) for each GO term. (**D**) Result of the Reactome Pathway analysis based on the list of genes significantly deregulated (869 genes, adjusted p-value <= 0.01 and 0.5 < FC < 2) and unique to Patient II.2 (i.e. that were not deregulated in the AR-CMT2A patient with a mutation on a different gene). We selected significantly enriched reactome pathways (p-value <=0.05) and ordered them by the number of entities found in descending order. On the left, the bars represent the number of DEGs found in each reactome pathway (entities), on the right, the -Log10(FDR). Pathways regarding cholesterol metabolism are highly significant (pvalue and FDR<0.01), and highlighted in red.

Gene Set Enrichment Analysis (GSEA) on the entire set of expression data highlighted a total of 1976 Differentially Expressed Genes (DEGs) between Patient II.2 and control’s hiPSC-MNs (Padj <0.05), of which 734 were up-regulated (FC>2) and 706 down-regulated (FC<0.5). When using a padjusted value of 0.01 as the significant threshold, we identify 1318 DEGs including 567 upregulated and 557 downregulated transcripts (FC>2 or FC<0.5) (Figure 1B). In the same batch, we had sequenced the mRNAs from hiPSC-MNs from another patient, affected with a completely different CMT subtype (AR-CMT2A, MIM), homozygous for the c. p.Arg298Cys (c.892C>T) missense mutation in the *LMNA* gene (De Sandre-Giovannoli et al., 2002). In order to further refine the list of deregulated genes, and to keep the deregulated genes, which arespecific to the VRK1 mutations and associated pathomechanisms, we “filtered” the list of significant DEGs (Padj <0.01 with FC>2 or <0.5) obtained for patient II.2, by removing all genes, which were found deregulated in both our patient II.2 and the other patient with the mutation in *LMNA*. This gave a list of 869 genes that was used for further analysis (Supplemental Table 1), such as GO annotation and Reactome.

### Enrichment analysis reveals a downregulation of genes involved in Cholesterol metabolism and the upregulation of transcripts involved in synaptic efficiency and plasticity in the patient’s motor neurons

Functional Gene Ontology (GO) annotation, showed enrichment in 122 and 45 terms, in the Biological Process (BP) and Cellular Component (CC) groups, respectively. Interestingly, out of the most 10 significantly enriched BP terms, more than half are related to cholesterol biosynthesis or metabolism, (Figure 1C). In particular, when looking at the enriched GO terms in the downregulated genes, the Top 10 are all related to cholesterol metabolism (Table 1 and Figure 2). Pathway analysis using reactome (Milacic et al., 2024) confirmed that cholesterol metabolism is a major defective pathway in the patients’MNs, specifically, the regulation of cholesterol biosynthesis by SREBP (Figure 1D, Figure 2C).

**Figure 2.**
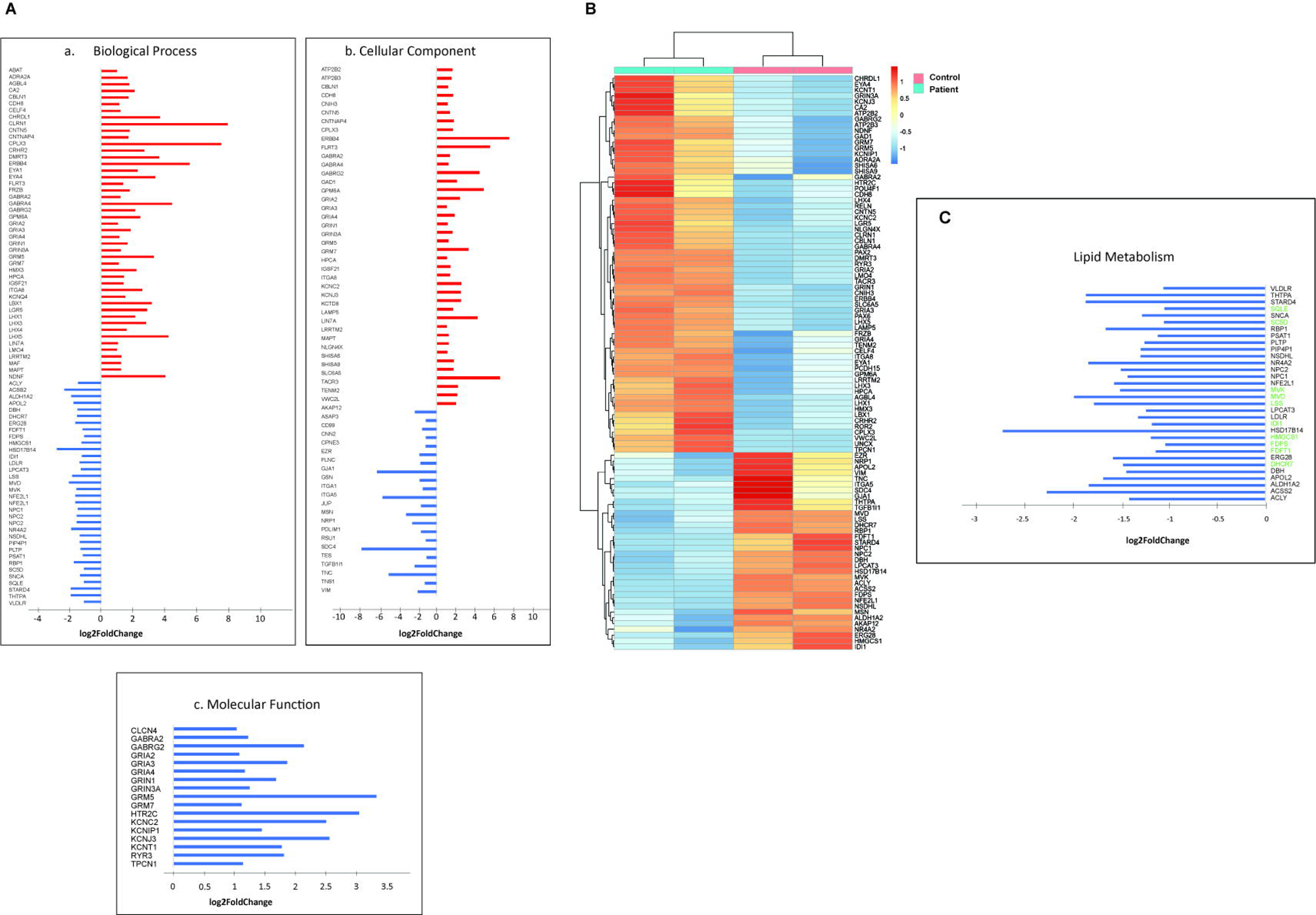
Transcriptional profiles of hiPSC-derived Motor Neurons (hiPSC-MN) from Patient II.2 and one control. (**A**) Bar graphs showing the Fold Change for the 122 genes found in the Top10 GO terms (a:BP, b:CC, c. MF). (**B**) Heatmap of supervised hierarchical clustering of the 4 samples for the genes shown in (**A**). (**C**) Bar graphs showing the Fold Change for the genes involved in lipid metabolism found in the Top10 GO terms. Transcripts related to regulation of cholesterol biosynthesis by SREBP are indicated in green on the right. *For (A) and (C), Fold Change is shown as Log2(FoldChange). Bars corresponding to downregulated transcripts are in blue. Bars corresponding to upregulated transcripts are in red*

**Table 1.**
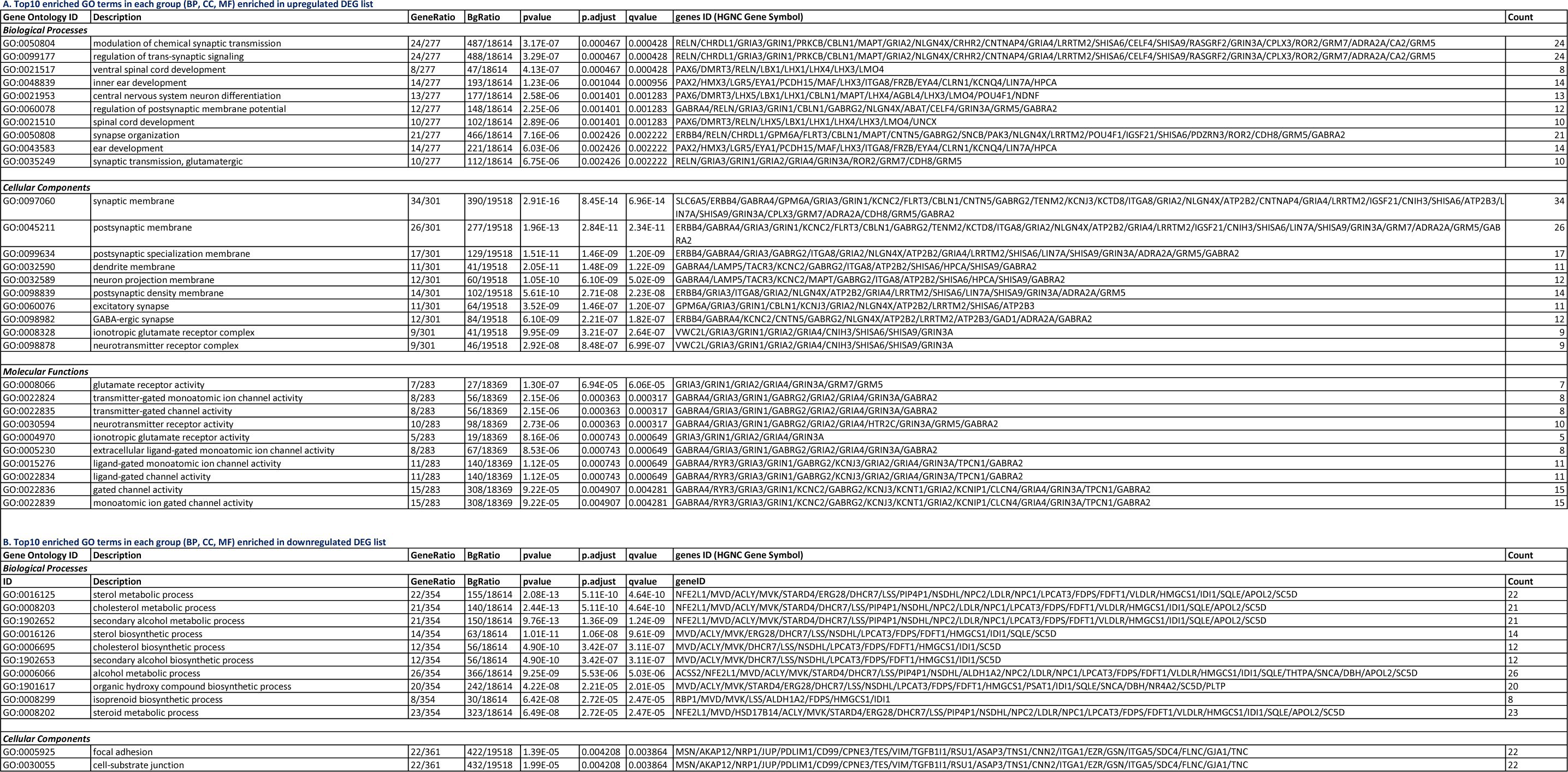
Results of Gene Ontology Enrichment Analysis for patient II.2 Differential Expressed Genes. (**A**) Top 10 enriched Gene Ontology (GO) terms in each group: BP (Biological Process), CC(Cellular Component), MF(Molecular Function). GO enrichment was calculated on the list of differentially expressed genes (DEGs) with **significant upregulation** (padj<0.01, and FC> 2). (**B**) Top 10 enriched Gene Ontology (GO) terms in each group: BP (Biological Process), CC(Cellular Component), MF(Molecular Function). GO enrichment was calculated on the list of differentially expressed genes (DEGs) with **significant downregulation** (padj<0.01, and FC< 0.5). *GO terms are sorted by padjusted value from most significant to least significant*.

Conversely, for upregulated genes, GO Enrichment analysis shows an enrichment of GO terms related to synaptic transmission and signaling and spinal cord development (Table 1 and Figure 1C). In particular, transcripts from genes encoding glutamate receptors subunits are significantly upregulated: NMDA (*GRIA2*, *GRIA3*, *GRIA4*), AMPA (*GRIN1*, *GRIN3A*) and metabotropic mGlu (*GRM5*, *GRM7*) (Table1 and Figure 2A). Surprisingly, transcripts encoding GABA receptors (*GABRA2*, *GABRA4*, *GABRG2*) are also significantly upregulated in the patient’s spinal motor neurons.

### ERBB4 signaling is significantly deregulated in Patient II.2’s hiPSC-MNs

Most interestingly, the reactome analysis points out dysregulations of ERBB2/ERBB4 signaling, with significant enrichment in pathways involving the epidermal growth factor member ERBB4 (p<0.05). Genes implicated in these pathways are highlighted in green in Figure 1D: three are upregulated (*ERBB4*, *GABRG2* and *WWOX*) and four are downregulated (*SHC1*, *NEDD4, NRG1* and *YAP1*) (Table1 and Figure 2A). ERBB4 itself is strongly upregulated with a Fold Change =47 and a highly significant padjusted value (3.45E^-83^).

This strong upregulation was confirmed by Western-Blot performed on proteins extracted from D30 hiPSC-MNs from patient II.2 and one control (Figure 3, n=3 experiments).

**Figure 3.**
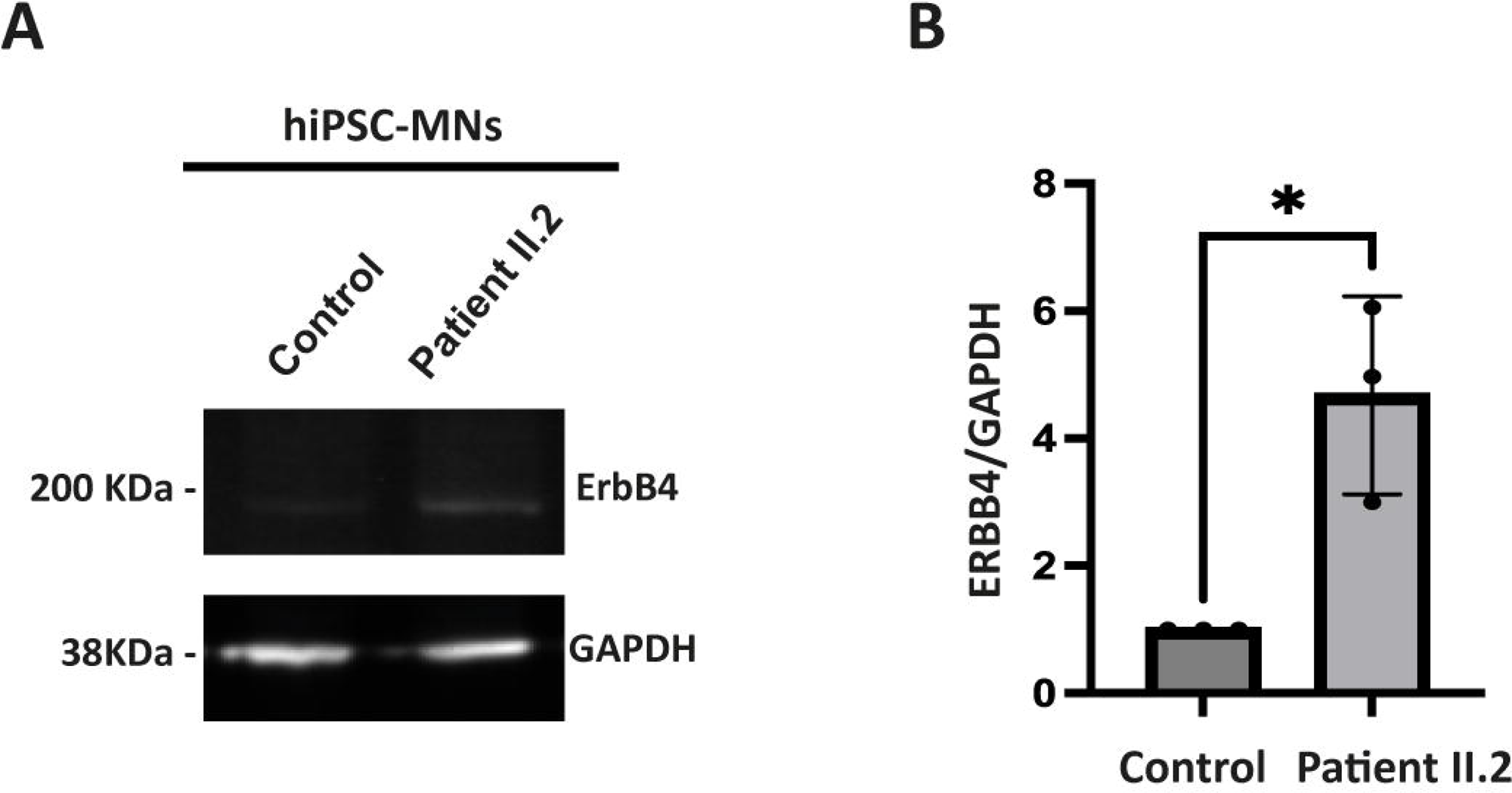
Aletered expression of ERBB4 in Patient II.2’s hiPSC-MNs. (**A**) Western Blot showing expression levels of ErbB4 in hiPSC-MNs from Patient II.2 and one control (AG161B) at D30 of differentiation. (**B**) Quantification of ErbB4 expression levels in Western Blot observed in (A). Data are expressed as mean ± sem (n=3 differentiations) and normalized to the control values. Statistical analysis: unpaired Student’s t-test.

### Proteins involved in splicing deregulated in SMA are not deregulated in the patient’s transcriptome

From the 152 genes, encoding proteins involved in splicing and identified deregulated in (Varderidou-Minasian et al., 2021), only one was found significantly (padj<0.01) deregulated in our patient (POLR*2A*, FC=0.62).

## Discussion

Mutations in the *VRK1* gene cause a range of diseases, described as SMA, distal SMA, ALS, juvenile ALS, HMSN, pure motor neuropathy (distal HMN), that we name VRK1-related motor neuron diseases. They affect the axons of lower, and also often upper, motor neurons. We have described VRK1, as the gene responsible for dHMN in two patients from a family, with a clinical diagnosis of distal Hereditary Motor Neuropathy associated with upper MN signs (El-Bazzal et al., 2019). There are currently more than thirty mutations described in this gene (Lazo & Morejon-Garcia, 2023), but the pathomechanisms underlying the disease remain unclear. By studying cells from patient II.2 described in (El-Bazzal et al., 2019), including induced Pluripotent Stem Cell (iPSC) derived Motor Neurons (hiPSC-MNs), we have shown that the mutations identified in *VRK1* lead to reduced levels of VRK1 in the nucleus. We have shown that this depletion alters the dynamics of coilin, a phosphorylation target of VRK1 (El-Bazzal et al. 2019). hiPSC-MNs from the patient, display Cajal Bodies (CBs) disassembly and defects in neurite outgrowth and branching, these defects are consistent with a length dependent axonopathy (El-Bazzal et al. 2019). In the same hiPSC-MNs, we have measured the electrical activity and demonstrated a modification in the shape of the Action Potential (AP) and a depolarized AP threshold with no alteration of the proportion of firing patterns. The smaller and broader AP recorded in patients with VRK1 mutations was associated with a decrease of the peak Na + currents accompanied by a decrease in the length of the ankyrinG (AnkG)-positive Axonal Initial Segment (AIS) (Bos et al., 2022). Our results reflect a hypoexcitable state of our patient’s MNs, opposite to the common idea that hyperexcitability is an intrinsic feature of motor neurons defects, like in Amyotrophic Lateral Sclerosis (ALS) (Wainger et al., 2014) or Spinal Muscular Atrophy (SMA) (Liu et al. 2015). Hypoexcitability has, however, recently been proposed to reflect MN firing defects in mouse models of ALS (Filipchuk et al. 2021; Martinez-Silva et al. 2018), in accordance with results obtained in hiPSC-MNs of ALS patients with C9ORF72 triplet expansions (Sareen et al., 2013).

In order to further explore the mechanistic links between hyperexcitability and hypoexcitability and to investigate the missing cues between VRK1 loss of function and the defects of patients’ motor neurons (disassembly of CB, altered neurite growth, alterations of the AIS and abnormal AP waveform), we have performed Bulk RNA-Seq in hiPSC-derived Motor Neurons from one patient (patient II.2) published in (El-Bazzal et al., 2019) and (Bos et al., 2022) as compared to a control.

First of all, our results do not bring further support to the hypothesis we made of defective RNA metabolism in dHMN-VRK1, due to Cajal Bodies disassembly. Indeed, we observed no deregulations in the genes encoding proteins involved in splicing, that were found deregulated in hiPSC-MNs from SMA patients (Varderidou-Minasian, Verheijen et al. 2021).

However, one of the most deregulated gene in patient II.2 hiPSC-MN is *ERBB4*, which is strongly upregulated (FC=47, padj= 3.45E-83) in Patient II.2 motor neurons. Most interestingly, upon examining the sequencing reads, we demonstrate that ERBB4 *ERBB4* is not just upregulated, but that it is ectopically expressed in hiPSC-MNs from the patient, while it is not endogenously expressed in controls’ hiPSC-MNs (data not shown). At the protein level, we confirm this strong upregulation of ErbB4 in Patient’s II.2 hiPSC-MNs, but the levels of ErbB4, though very low, are not completely at zero, in the control’s hiPSC-MNs. This discrepancy between transcript and protein quantification may be due to the fact that around 30% of cells are not motor neurons in the culture dish at the end of differentiation, as can be seen from HB9 immunostaining (see supplementary figure 1).

ErbB4 is a tyrosine kinase transmembrane receptor, which is activated by binding its ligand, Neuregulin 1 (Mei & Nave, 2014; Modol-Caballero, Santos, Navarro, & Herrando-Grabulosa, 2017). Our results confirm those published by Casanovas et al. (Casanovas et al., 2017), in primary cultures of dissociated mouse spinal cord, showing that Neuregulin 1 (NRG1) is concentrated in postsynaptic sites, on the lower motor neurons, while ErbB4 receptors are present in the presynaptic compartment, suggesting that they are not present in the membranes of spinal lower motor neurons. ErbB4 mutations have been described in a rare subtype of Amyotrophic Lateral Sclerosis (Amyotrophic Lateral Sclerosis Type 19, MIM 615515) and shown to disrupt the Neuregulin 1/ErbB4 Pathway (Takahashi et al., 2013). Also, it has been shown, in an *in-vitro* spinal cord organotypic culture model subjected to excitotoxicity, that NRG1 reduces motor neuron cell death and promotes neurite growth (Modol-Caballero et al., 2017). Here, additionally to ErbB4 upregulation, there is a significant downregulation (FC= 0.367, padj value= 3.4E-06) of NRG1 in the hiPSC-derived spinal motor neurons from the patient II.2, with VRK1 mutations. This decrease in the levels of NRG1 transcripts, might be a compensatory mechanism trying to restore balance in NRG1/ErbB4 signaling. Here we don’t know if there is paracrine aberrant signaling between hiPSC-MNs, (which do not exist *in vivo*, because the only synapse for lower motor neurons is the neuromuscular junction), or if there is an autocrine NRG1 mediated signaling in those MNs, as ErbB4 is expressed in the patient’s MNs.

While we do not know yet the exact mechanisms leading to the upregulation of ErbB4 and downregulation of NRG1 in our patient’s motor neurons, our RNA-Seq data clearly suggest decreased NRG1/ErbB4 signaling in motor neurons from patients with bi-allelic mutations in *VRK1*.Thisfurther underlines the relevance of this pathway in motor neuron diseases and axonal peripheral neuropathies. Indeed, functional analysis of the mRNA-Seq data show a significant downregulation of genes involved in cholesterol biosynthesis and metabolism, in particular, those regulating cholesterol biosynthesis by SREBP. These downregulations are most probably a consequence of decreased NRG1/ErbB4 signaling, which has been shown to be involved in regulating the SREBP-2 cholesterol biosynthetic pathway. Indeed, in human breast cancer cells, NRG1 binding to ErbB4 has been shown to activate SREBP-2 by inducing its cleavage and to increase the expression of cholesterol metabolic enzymes, LDL uptake and cholesterol biosynthesis (Haskins et al., 2015). Here, our results show the opposite, with decreased NRG1 levels, and increased ERBB4 levels. This imbalance likely leads to altered NRG1/ErbB4 signaling, and decreased cholesterol metabolism, through downregulation of targets of SREBP. Considering the fact that the ErbB4 isoform expressed is our patient’s hiPSC-MN is the JM-b (Elenius et al., 1997) (data not shown), which cannot be cleaved to release the soluble ICD domain (Haskins et al., 2015), our results confirm that formation of ErbB4 ICD is not required to activate SREBP. Altered lipid metabolism has been described, by multi-omic analysis, in 17 ALS patient hiPSC-derived spinal MNs (Lee et al., 2021), strengthening the importance of lipids in the structure and function of axons. Indeed, axonal growth and regeneration rely on many signaling pathways, which use the fixation of a ligand on its receptor. These receptors are located in specialized raft membrane microdomains with a precise lipid composition, cholesterol being one of its major components. Therefore, any disruption in cholesterol metabolism, might have consequences on axonal growth and survival, by perturbating the precise membrane lipid balance necessary to define neuronal architecture (Jose, Sivanand, & Channakeshava, 2021; Modol-Caballero et al., 2017; Rosello-Busquets et al., 2019).

In the case of patients with *VRK1* mutations, we have demonstrated previously, that hiPSC-MNs have defects in neurite outgrowth and branching (El-Bazzal et al., 2019). These defects are likely to be linked to the altered cholesterol and lipid metabolism, found here by the RNA-Seq analysis.

Another consequence of altered cholesterol metabolism might be the upregulation, in our patient’s hiPSC-MNs, of genes encoding NMDA and AMPA glutamate receptors subunits. Indeed, cholesterol is a structural component of cellular membranes particularly enriched in synapses, and its depletion in rat hippocampal cultures resulted in a significant reduction of both NMDA receptor (NMDAR) and AMPA/kainate receptor-mediated evoked excitatory postsynaptic currents (Korinek et al., 2020). At the opposite, Brachet et al. (Brachet et al., 2015) reported that cholesterol levels participate in synaptic plasticity by engaging small GTPase signaling and driving AMPAR trafficking to the synaptic membrane. Finally, in rat organotypic hippocampal slices, Li et al (Li, Woo, Mei, & Malinow, 2007) have shown that normal activity-driven glutamatergic synapse development is impaired by deficits in NRG1/ErbB4 signaling leading to glutamatergic hypofunction and that preventing NRG1/erbB4 signaling destabilizes synaptic AMPA receptors and leads to loss of synaptic NMDA currents and spines. For dHMN-VRK1, we are here in the context of the Peripheral Nervous system, and lower motor neurons (LMN), which axons are affected in the pathology (dHMN-VRK1), are cholinergic neurons, not glutamatergic. However, in the patients, those LMNs do receive input signals from Upper Motor Neurons (UMNs) and this communication might be affected as a result of impaired NRG1/ErbB4 signaling and the resulting in altered cholesterol metabolism. In our *in vitro* model, the hiPSC-MNs, unlike in the PNS, are able to communicate and most likely to establish synapses between them and the increase of glutamate receptors transcripts in our patients’ hiPSC-MNs, might reflect a compensatory mechanism to reverse decreased glutamatergic signaling or fast removal of receptors from the synapse. This increase of AMPA and NMDA receptors’ transcript might be a sign of excitotoxicity (Brockington et al., 2013), but this would be contradictory to what we showed in hiPSC-MNs from our patients (Bos et al., 2022). We still need to confirm that these receptors are indeed enriched in the patient’s MNs. Although translation of this observation into what is happening in the patients’ nerves is complicated, the deregulations observed in our model point on defective synaptic plasticity.

Taken together, our results help understanding the link between VRK1 loss of function and the neurite outgrowth observed in motor neurons from patients with mutations in VRK1. Nrg1/ErbB4 signaling and the resulting downregulated cholesterol metabolism pathway appear as a key defective processes in the physiopathology of VRK1 related motor neuron diseases. These results open new avenues for defining treatments in VRK1 related motor neuron diseases by targeting cholesterol metabolism.

## Material and Methods

### Patients’ cells and cell lines

The control hiPSC cell lines (hff19 and AG161B) were provided by MaSC, our Cell Reprogramming and Differentiation Facility at U1251/Marseille Medical Genetics (Marseille, France). This iPS cell line was obtained by reprogramming of a commercial fibroblast cell line (FibroGRO™ Xeno-Free Human Foreskin Fibroblasts, #SCC058, Merck Millipore, Germany). Patients’ hiPSC cell lines were obtained by reprogramming of skin primary fibroblasts from: i)the female patient II.2, affected with autosomal recessive distal Hereditary Motor Neuropathy (dHMN) due to bi-allelic in *VRK1* described in (El-Bazzal et al., 2019) and ii) from a patient homozygous for the c. p.Arg298Cys (c.892C>T) missense mutation in the *LMNA* gene (De Sandre-Giovannoli et al., 2002). All hiPSCs included in the study are declared in a collection authorized by the competent authorities in France (declaration number DC-2018-3207).

### Generation, culture and maintenance of hiPSCs

Reprogramming of skin fibroblasts from patient and control into induced pluripotent stem cells (hiPSC) was performed by our Cell Reprogramming and Differentiation Facility (MaSC) at U 1251/Marseille Medical Genetics (Marseille, France), using Sendai virus-mediated transduction (Thermofisher Scientific, #A16517) or nucleofection (Amaxa, Lonza, # VAPD-1001 NHDF) of episomal vectors containing OCT3/4, SOX2, KLF4, C-MYC and shRNA against p53. hiPSCs colonies were picked about two weeks after transduction/nucleofection based on ES cell-like morphology. Colonies were grown and expanded in mTeSR1 medium (Stemcells technologies, #85850) on BD Matrigel™ (Corning, #354277) coated dishes. hiPSCs clones were fully characterized using classical protocols as described in (El-Bazzal et al., 2019).

After thawing, hiPSCs were plated on Matrigel (Corning Life Sciences, USA, #354248) coated plastic dishes and maintained in mTeSR1 medium (Stemcell Technologies, Canada, #05851), which was exchanged daily.

### Differentiation of hiPSCs into spinal Motor Neurons (hiPSC-MNs)

For differentiation of hiPSCs into spinal MNs, we used an established 30 days differentiation protocol based on early activation of the Wnt signaling pathway, coupled to activation of the Hedgehog pathway and inhibition of Notch signaling (Maury et al., 2015).

Using this protocol, we obtained mature MNs, which express acetylcholine transferase and can fire Action Potentials(Bos et al., 2022; El-Bazzal et al., 2019). For each experiment, differentiation efficiency was assessed by HB9 and ISLET1 immunostaining as described (El-Bazzal et al., 2019). Neurites were labelled using antibody to the neurofilament Medium chain (NF-M), a neuronal cytoskeletal protein (**Supp Figure 1**). Both hiPSCs and differentiated hiPSC-MNs were tested for mycoplasma contamination every other week.

### Immunostaining and microscopy

At Day30 of differentiation (DIV30), hiPSC-MNs, grown on matrigel coated in 4-wells culture chambers from Sarstedt (#94.6140.402) were washed with Dulbecco’s Phosphate Buffer Saline buffer (PBS) and fixed in 4% paraformaldehyde in PBS for 15 minutes. Cells were then washed for 10 min in PBS and were permeabilized using 0.1% Triton X-100 in PBS for 10 min at room temperature. After blocking for 30 min at room temperature in PBS containing 0.1%Triton X-100 and 1% Bovine Serum Albumin (BSA), hiPSC-MNs were incubated overnight at 4°c with the primary antibodies diluted in the blocking solution. Primary antibodies used were: rabbit polyclonal to Olig-2 (Merck Millipore, #AB9610) (dilution 1:500), Mouse monoclonal to Islet-1 (Biorbyt, #Orb95122) (1:200), mouse monoclonal to HB9 (DSHB, #81.5C10) (1:40), Chicken polyclonal to NF-M (Biolegend, #PCK-593P) (1:1000). After 3 washes in PBS, cells were incubated for 2 hours with the appropriate secondary antibody, in the blocking solution at room temperature in the dark. The following secondary antibodies were used at 1:1000 dilution: donkey Anti-Mouse IgG (DyLight® 550) (abcam, #ab96876), donkey Anti-Rabbit IgG (DyLight® 550) (abcam, #ab96892), goat anti-Chicken IgY H&L (Alexa Fluor® 488) (abcam, # ab150169)

The samples were then mounted in Vectashield mounting medium (Vector Laboratories, #H-2000) with 100ng/ml DAPI (4,6-diamidino-2-phenylindole), coverslipped, and sealed. Digitized microphotographs were recorded using an ApoTome fluorescent microscope (ZEISS, Germany), equipped with an AxioCam MRm camera. These images were merged with the ZEN software, and were treated using ImageJ software (National Institutes of Health, MD, USA).

### Protein extraction and Immunoblotting

Protein extraction and immunoblotting were performed from hiPSC-MNs at day 30 of differentiation. Cell pellets were lysed on ice with 50 µl of commercialized RIPA lysis buffer (Thermo Scientific #89900) containing triton 1X Triton X-100, 0.1% SDS, 0.15M NaCl, sodium desoxycholate, 1mM EDTA, 20µM Tris–HCl, pH7.5, and a cocktail of proteases and phosphatases inhibitor (Thermo scientific #78442). The lysate was passed through an 18-21gauge needle and submitted to 4 cycles of sonication (15 sec followed by 15 second pause on ice between each sonication). After centrifugation for 10 min at 10000 g at 4°C, the supernatant was removed, protein concentration was measured by use of a BCA Protein Assay (SIGMA Aldrich) and samples stored at -80°C.

We used the following primary antibodies: rabbit polyclonal antibody to ErbB4 (Cell Signaling, #sc-(111B2) 4795) and mouse polyclonal antibody to GAPDH (Abcam, #ab125247), diluted at 1:1000 for all the antibodies. GAPDH served as a loading control. Secondary antibodies were donkey anti-rabbit IRDye 680 and donkey anti-mouse IRDye 800, (Li-Cor Biosciences), diluted at 1:10000.

### RNA extraction

At day 30 of the differentiation protocol, cells were collected using a scraper in PBS1X. After centrifugation for 5 min at 1000 g, the supernatant was removed, and the cell pellets was stored at -80°C. Total RNA was extracted from the pellet using Purelink silica-membrane, anion exchange resin, spin-column kits, following the manufacturer’s instructions (Purelink RNA Minikit, #<u>12183018A</u>, ThermoFisher Scientific, USA).

### RNA sequencing

RNA-Sequencing and bioinformatics analysis was performed by the Genomics and Bioinformatics facility (GBiM) from the U 1251/Marseille Medical Genetics lab. mRNA sequencing (mRNA-Seq) was performed, in duplicate, on total RNA samples extracted from hiPSC-derived motor neurons (iPSC-MN) at day30 of differentiation, from one patient and one control (see above). Before sequencing, the quality of total RNA samples was assessed by using the Agilent Bioanalyzer (Agilent Technologies, USA): only RNAs with RNA Integrity Numbers (RIN) above 8 were deemed suitable for sequencing and used for library preparation. For each sample, a library of stranded mRNA was prepared from 500 ng of total RNA after capture of RNA species with poly-A tail poly(A), using the Kapa mRNA HyperPrep kit (Roche, Switzerland), using 500 ng of total RNA and following the manufacturer’s instructions. The quality and profile of individual libraries have been quantified and visualized using Qubit™ and the Agilent Bioanalyzer dsDNA High Sensibility Kit (Agilent Technologies, USA) respectively. Indexed libraries were pooled and sequenced (2*75 bp paired-end sequencing) on an Illumina NextSeq 500 platform (Illumina, USA).

### Data processing and differential gene expression analysis (DGE)

The quality of sequencing reads was assessed using fastQC v0.11.5 (https://www.bioinformatics.babraham.ac.uk/projects/fastqc/). Raw sequencing reads were mapped to the human reference genome GRCh38 (hg38) and sorted using STAR v2.7.2b (Dobin & Gingeras, 2015), and bam files were indexed using Sambamba v0.6.6 (Tarasov, Vilella, Cuppen, Nijman, & Prins, 2015). After mapping, the number of reads per feature (GENCODE v34 annotations) was determined using Stringtie v1.3.1c (Pertea et al., 2015).

The presence of the expected mutations in each sample was verified by visualization of the reads from the bam files on the Integrated Genome Viewer IGV v 2.16.1, using GRCh38 as the reference genome. Differential gene expression analysis was performed using a Wald test from the DESeq2 package (v1.18.1) (Love, Huber, & Anders, 2014). *P*-values were adjusted for multiple testing using the method described by Benjamini and Hochberg (Benjamini & Hochberg, 1995). **Only transcripts with an adjusted *P*-value (FDR, False Discovery Rate) below 0·01 were considered as significantly differentially expressed**. Relative expressions of the most variable features between samples have been plotted as heatmaps using the Pheatmap R package. Differentially Expressed Genes (DEGs) were visualized in the form of volcano plots, representing the Log2FC (log2 of the expression fold change) and the adjusted P-value, generated by the EnhancedVolcano R package. Enrichment (Gene set enrichment Analysis, GSEA) and overrepresentation (Singular Enrichment Analysis, SEA) of GO term annotation in the DEGs were performed using a Mann-Whitney test and hypergeometric test respectively, using enrichGO from the R-package clusterProfiler (v3.10.15). For functional Gene Ontology (GO) annotation, we selected the differentially expressed genes, with a statistical significance of *Padj* value < 0.01 (SEA), and with a fold change over 2 or below 0.5, for upregulated and downregulated genes respectively. The list of significant DEGs was “filtered” with data obtained from other hiPSC-MNs, in the same sequencing batch, from a patient affected with a totally different CMT subtype (AR-CMT2A,MIM 605588), and homozygous for the c. p.Arg298Cys (c.892C>T) missense mutation in the *LMNA* gene (De Sandre-Giovannoli et al., 2002). We removed the genes deregulated both in Patient II.2 and the AR-CMT2A patient, which are hallmarks of axonal degeneration common to any axonal CMT, to keep the genes, that were uniquely deregulated in our patient II.2 and more likely to be related to VRK1. Pathway analysis was conducted, using PathwayBrowser from Reactome.org, which uses a hypergeometric test with an adjustment of the p-value (BH) to detect pathway, enriched in the dataset of genes (DEG).

## Statistical analysis

For experiments, other than RNA-Seq, the applied statistical tests, as well as the number of replicates, are indicated in the figure legends. Results were obtained from at least three independent experiments.

The significance of the results was evaluated with unpaired student’s t-test according to experimental design. Statistical analyses were performed using GraphPad Prism. All data are presented as mean ± standard error of the mean (SEM).

## Acknowledgements

We would like to thank the patients and families for their kind cooperation and their participation in this study. We thank Claire El Yazidi, from the Cell Reprogramming and Differentiation Facility (MaSC) at U 1251/Marseille Medical Genetics (Marseille, France) for reprogrammation and characterization of patient’s II.2 fibroblasts into iPSC cell line and for providing control iPSC lines. We are grateful to Mazen Bou Malhab for his help in the design of figures.

## Figure legends

**Supplementary Table1. Results of Differential Gene Expression analysis for Patient II.2 after removing DEGs common to the AR-CMT2A patient.**

Genes are sorted by padjusted value in ascending order (from most significant to least significant). FC=Fold Change; padj=padjusted value. Basemean represents the average of the normalized count values, dividing by size factors, taken over all samples. Pvalue=P-value of the test for the gene or transcript. padj=Adjusted P-value for multiple testing for the gene or transcript.NA=Not Assessed

