## Supplementary figures and images for "Altered NRG1/ErbB4 signaling and cholesterol metabolism dysregulation are key pathomechanisms in VRK1-related motor neuropathies and motor neuron diseases"

### Supplementary Figure 1

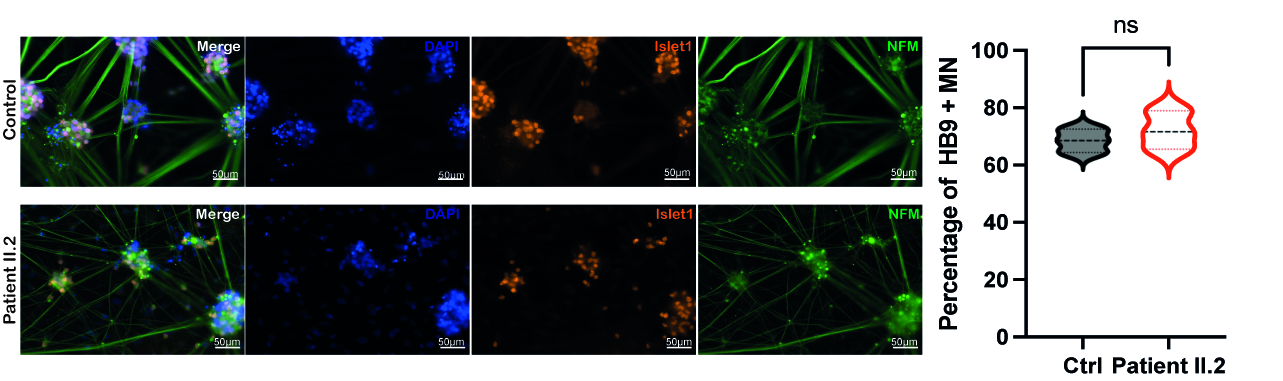
