## Supplementary Table 1 for "Altered NRG1/ErbB4 signaling and cholesterol metabolism dysregulation are key pathomechanisms in VRK1-related motor neuropathies and motor neuron diseases"

| padj | pvalue | geneID | gene_name | baseMean | log2FoldCha | lfcSE | stat |
|---|---|---|---|---|---|---|---|
| 1.52E-164 | 6.15E-169 | ENSG00000181541.6 | MAB21L2 | 7165.8531 | 6.6008356 | 0.2382578 | 27.704589 |
| 1.30E-135 | 1.06E-139 | ENSG00000165970.12 | SLC6A5 | 6364.0408 | 6.57741 | 0.2614079 | 25.161477 |
| **3.45E-83** | **9.80E-87** | **ENSG00000178568.15** | **ERBB4** | **3544.4607** | **5.5621372** | **0.2817717** | **19.739874** |
| 3.21E-66 | 1.56E-69 | ENSG00000067646.12 | ZFY | 1004.4602 | -7.576552 | 0.4298561 | -17.62579 |
| 1.57E-64 | 8.26E-68 | ENSG00000075891.22 | PAX2 | 1140.5139 | 4.6997561 | 0.2701023 | 17.39991 |
| 3.56E-64 | 2.02E-67 | ENSG00000100078.4 | PLA2G3 | 857.13473 | -7.742113 | 0.4462675 | -17.34859 |
| 1.12E-61 | 6.83E-65 | ENSG00000165246.14 | NLGN4Y | 3509.6386 | -3.658707 | 0.2150813 | -17.01081 |
| 3.58E-61 | 2.32E-64 | ENSG00000241859.7 | ANOS2P | 1060.8704 | -5.081401 | 0.2999826 | -16.93899 |
| 1.18E-55 | 8.16E-59 | ENSG00000106772.18 | PRUNE2 | 22396.532 | -3.490112 | 0.2158339 | -16.17036 |
| 2.07E-42 | 1.85E-45 | ENSG00000124785.9 | NRN1 | 3482.2952 | 4.4130144 | 0.3118562 | 14.150797 |
| 1.25E-41 | 1.16E-44 | ENSG00000116833.14 | NR5A2 | 3273.5926 | 4.6967493 | 0.3349828 | 14.020869 |
| 1.42E-39 | 1.44E-42 | ENSG00000197705.10 | KLHL14 | 402.01832 | 5.024064 | 0.3673991 | 13.674676 |
| 3.85E-38 | 4.38E-41 | ENSG00000007372.23 | PAX6 | 670.99372 | 5.6423267 | 0.420318 | 13.423949 |
| 6.03E-38 | 7.10E-41 | ENSG00000185352.8 | HS6ST3 | 9554.702 | 2.933393 | 0.2191042 | 13.388122 |
| 3.08E-37 | 3.75E-40 | ENSG00000284862.3 | CCDC39 | 327.89111 | 5.6257412 | 0.4241377 | 13.263949 |
| 1.25E-35 | 1.57E-38 | ENSG00000099725.14 | PRKY | 662.24856 | -3.544474 | 0.2730517 | -12.98096 |
| 1.23E-34 | 1.60E-37 | ENSG00000184216.14 | IRAK1 | 1630.3714 | 3.582222 | 0.2798229 | 12.801747 |
| 2.22E-34 | 2.97E-37 | ENSG00000253159.3 | PCDHGA12 | 415.87641 | -6.567279 | 0.5149262 | -12.75383 |
| 9.15E-33 | 1.30E-35 | ENSG00000147596.4 | PRDM14 | 595.11331 | 4.9422169 | 0.3967767 | 12.455914 |
| 1.26E-32 | 1.85E-35 | ENSG00000145730.20 | PAM | 9650.8865 | 2.6494385 | 0.2131863 | 12.427808 |
| 4.04E-32 | 6.07E-35 | ENSG00000109158.11 | GABRA4 | 897.74221 | 4.4497932 | 0.3608236 | 12.332325 |
| 4.57E-32 | 7.23E-35 | ENSG00000125869.10 | LAMP5 | 2843.055 | 4.2632793 | 0.3460954 | 12.318218 |
| 5.54E-32 | 8.98E-35 | ENSG00000286619.1 | AC119673.3 | 270.88666 | -6.227335 | 0.5062594 | -12.30068 |
| 1.92E-31 | 3.19E-34 | ENSG00000120324.9 | PCDHB10 | 3625.5475 | -2.484679 | 0.2036974 | -12.19789 |
| 1.89E-30 | 3.30E-33 | ENSG00000287243.2 | AC005908.3 | 2858.0084 | 2.6419742 | 0.2200518 | 12.006144 |
| 1.32E-29 | 2.36E-32 | ENSG00000102384.13 | CENPI | 2024.3046 | -2.966337 | 0.2504882 | -11.84222 |
| 4.06E-29 | 7.50E-32 | ENSG00000204970.10 | PCDHA1 | 280.97526 | 4.4859162 | 0.3819461 | 11.744893 |
| 4.06E-29 | 7.57E-32 | ENSG00000182168.15 | UNC5C | 4848.3218 | -3.319239 | 0.2826301 | -11.74411 |
| 3.04E-28 | 5.80E-31 | ENSG00000131069.20 | ACSS2 | 10711.746 | -2.286696 | 0.197628 | -11.57071 |
| 5.76E-28 | 1.12E-30 | ENSG00000109084.14 | TMEM97 | 38611.377 | -2.885189 | 0.2505802 | -11.51403 |
| 7.95E-28 | 1.58E-30 | ENSG00000250120.8 | PCDHA10 | 1024.505 | -2.730742 | 0.2377781 | -11.48441 |
| 4.72E-26 | 9.76E-29 | ENSG00000121769.8 | FABP3 | 7353.8234 | -2.737165 | 0.2460947 | -11.1224 |
| 5.36E-26 | 1.13E-28 | ENSG00000196092.14 | PAX5 | 948.17672 | 3.9227583 | 0.3531043 | 11.109349 |
| 5.54E-26 | 1.19E-28 | ENSG00000151623.15 | NR3C2 | 408.87448 | 4.0101679 | 0.3611235 | 11.104699 |
| 2.81E-25 | 6.38E-28 | ENSG00000132256.19 | TRIM5 | 530.90332 | -3.834347 | 0.3500515 | -10.95367 |
| 1.17E-24 | 2.74E-27 | ENSG00000255622.3 | PCDHB17P | 240.69723 | -4.129873 | 0.3816574 | -10.82089 |
| 2.48E-24 | 5.95E-27 | ENSG00000269001.2 | AC092070.2 | 223.10091 | -5.309431 | 0.4939148 | -10.74969 |
| 1.29E-23 | 3.15E-26 | ENSG00000140836.17 | ZFHX3 | 6866.7453 | -2.755043 | 0.2600337 | -10.59494 |
| 3.69E-23 | 9.13E-26 | ENSG00000111218.12 | PRMT8 | 3877.5589 | 2.4531273 | 0.2337469 | 10.494803 |
| 5.07E-23 | 1.29E-25 | ENSG00000162772.17 | ATF3 | 2092.8656 | -2.288332 | 0.2187333 | -10.46174 |
| 6.83E-23 | 1.77E-25 | ENSG00000101670.12 | LIPG | 4831.9887 | -2.593572 | 0.2486192 | -10.43191 |
| 8.22E-23 | 2.17E-25 | ENSG00000217236.2 | SP9 | 1106.7171 | 3.9351562 | 0.3779156 | 10.412792 |
| 1.86E-22 | 4.98E-25 | ENSG00000185304.15 | RGPD2 | 2258.5376 | -2.582266 | 0.2498986 | -10.33326 |
| 1.33E-21 | 3.67E-24 | ENSG00000089250.19 | NOS1 | 831.68013 | 4.3556729 | 0.4295482 | 10.140126 |
| 2.11E-21 | 5.91E-24 | ENSG00000128617.2 | OPN1SW | 185.14728 | 5.753483 | 0.5700251 | 10.093385 |
| 7.94E-21 | 2.29E-23 | ENSG00000182348.7 | ZNF804B | 363.71831 | 3.2556441 | 0.3268813 | 9.9597136 |
| 4.43E-20 | 1.30E-22 | ENSG00000092068.20 | SLC7A8 | 3114.3692 | -2.422108 | 0.2475116 | -9.785838 |
| 5.50E-19 | 1.67E-21 | ENSG00000249624.9 | AP000295.1 | 231.65079 | -4.16741 | 0.4375923 | -9.5235 |
| 7.63E-19 | 2.35E-21 | ENSG00000171570.11 | RAB4B-EGLN | 1340.8637 | -7.908063 | 0.8334653 | -9.488173 |
| 8.81E-19 | 2.75E-21 | ENSG00000276805.2 | AL133216.2 | 349.67366 | 3.0663995 | 0.3237401 | 9.4717945 |
| 8.85E-19 | 2.80E-21 | ENSG00000131015.5 | ULBP2 | 336.97221 | -3.025223 | 0.3194541 | -9.469974 |
| 1.10E-18 | 3.51E-21 | ENSG00000187627.16 | RGPD1 | 2012.6142 | -2.430267 | 0.2572715 | -9.446312 |
| 1.87E-18 | 6.15E-21 | ENSG00000277954.1 | AC092376.2 | 6796.2802 | 2.1203689 | 0.2258747 | 9.3873665 |
| 2.26E-18 | 7.51E-21 | ENSG00000183036.11 | PCP4 | 1063.9276 | 2.2839921 | 0.243852 | 9.366303 |
| 2.26E-18 | 7.60E-21 | ENSG00000080200.10 | CRYBG3 | 669.48409 | 3.0714714 | 0.3279687 | 9.3651343 |
| 3.48E-18 | 1.20E-20 | ENSG00000159248.5 | GJD2 | 885.01714 | 2.5542413 | 0.2741542 | 9.3168068 |
| 6.51E-18 | 2.27E-20 | ENSG00000138821.13 | SLC39A8 | 3234.7414 | 3.8476216 | 0.4160139 | 9.2487813 |
| 9.59E-18 | 3.38E-20 | ENSG00000016082.15 | ISL1 | 11796.566 | -2.724157 | 0.2959104 | -9.206018 |

| | | | | | | | |
|---|---|---|---|---|---|---|---|
| 1.16E-17 | 4.18E-20 | ENSG00000183878.15 | UTY | 1016.3553 | -13.51062 | 1.4712248 | -9.183244 |
| 2.09E-17 | 7.64E-20 | ENSG00000049130.16 | KITLG | 3062.7735 | 1.8359585 | 0.2013516 | 9.11817 |
| 3.41E-17 | 1.27E-19 | ENSG00000260196.1 | AC124798.1 | 1498.9484 | -2.086255 | 0.2302004 | -9.062776 |
| 4.03E-17 | 1.52E-19 | ENSG00000064218.5 | DMRT3 | 199.2147 | 3.6420378 | 0.4027386 | 9.0431808 |
| 4.70E-17 | 1.79E-19 | ENSG00000180828.3 | BHLHE22 | 159.30078 | 4.609904 | 0.5107746 | 9.0253194 |
| 1.32E-16 | 5.25E-19 | ENSG00000189056.14 | RELN | 8682.5718 | 2.62096 | 0.294261 | 8.9069234 |
| 1.53E-16 | 6.16E-19 | ENSG00000198838.13 | RYR3 | 4530.1005 | 1.8114573 | 0.2037856 | 8.8890358 |
| 2.74E-16 | 1.14E-18 | ENSG00000136367.14 | ZFHX2 | 1717.4746 | -2.071839 | 0.2348787 | -8.820886 |
| 6.15E-16 | 2.57E-18 | ENSG00000283930.1 | AL117339.4 | 183.64423 | 4.5301413 | 0.5189752 | 8.7290122 |
| 7.31E-16 | 3.08E-18 | ENSG00000198692.10 | EIF1AY | 620.68756 | -12.79894 | 1.4697377 | -8.708318 |
| 8.50E-16 | 3.62E-18 | ENSG00000130675.15 | MNX1 | 1174.7852 | -2.170117 | 0.2497239 | -8.690066 |
| 1.11E-15 | 4.79E-18 | ENSG00000186862.20 | PDZD7 | 2207.1714 | -2.001921 | 0.2312174 | -8.658176 |
| 1.19E-15 | 5.15E-18 | ENSG00000178187.7 | ZNF454 | 385.37448 | 2.5189645 | 0.2912134 | 8.6498925 |
| 1.40E-15 | 6.15E-18 | ENSG00000215580.11 | BCORP1 | 178.25881 | -7.954637 | 0.9217809 | -8.629639 |
| 1.70E-15 | 7.70E-18 | ENSG00000271474.1 | AC106881.1 | 1117.9615 | -3.221724 | 0.3744464 | -8.603966 |
| 1.70E-15 | 7.74E-18 | ENSG00000145423.5 | SFRP2 | 768.55371 | 4.1180448 | 0.4786508 | 8.6034422 |
| 3.13E-15 | 1.46E-17 | ENSG00000274049.4 | INO80B-WB | 553.70237 | -12.63407 | 1.4811182 | -8.53009 |
| 3.89E-15 | 1.83E-17 | ENSG00000114805.17 | PLCH1 | 1387.5354 | 2.4143133 | 0.2839024 | 8.5040257 |
| 2.00E-14 | 9.81E-17 | ENSG00000147065.17 | MSN | 8674.6437 | -3.143972 | 0.3784671 | -8.30712 |
| 2.25E-14 | 1.13E-16 | ENSG00000188211.9 | NCR3LG1 | 9486.4225 | -2.239926 | 0.2701863 | -8.290303 |
| 2.29E-14 | 1.16E-16 | ENSG00000152894.14 | PTPRK | 700.7418 | 2.2220314 | 0.268126 | 8.287266 |
| 2.40E-14 | 1.23E-16 | ENSG00000204110.6 | LINC02520 | 639.33966 | -4.135775 | 0.499456 | -8.28056 |
| 2.96E-14 | 1.52E-16 | ENSG00000163536.12 | SERPINI1 | 3142.5812 | 1.7800557 | 0.2156441 | 8.2545976 |
| 3.30E-14 | 1.72E-16 | ENSG00000236501.6 | AC074286.1 | 1749.7769 | 1.7985581 | 0.2182725 | 8.2399683 |
| 3.30E-14 | 1.74E-16 | ENSG00000131016.17 | AKAP12 | 81898.779 | -2.21689 | 0.269073 | -8.238991 |
| 4.15E-14 | 2.20E-16 | ENSG00000148600.15 | CDHR1 | 10850.42 | -2.247087 | 0.273688 | -8.210394 |
| 5.02E-14 | 2.69E-16 | ENSG00000186897.5 | C1QL4 | 1960.3943 | 2.1807538 | 0.2663819 | 8.1865691 |
| 5.39E-14 | 2.91E-16 | ENSG00000173930.9 | SLCO4C1 | 837.38013 | -2.755434 | 0.3369718 | -8.177047 |
| 7.08E-14 | 3.85E-16 | ENSG00000101938.15 | CHRDL1 | 1146.9296 | 3.7023113 | 0.45465 | 8.1432128 |
| 8.30E-14 | 4.54E-16 | ENSG00000120328.6 | PCDHB12 | 716.03923 | -2.416972 | 0.2975415 | -8.123142 |
| 9.33E-14 | 5.18E-16 | ENSG00000114861.22 | FOXP1 | 13600.093 | -2.637048 | 0.3252744 | -8.107148 |
| 1.53E-13 | 8.57E-16 | ENSG00000230847.4 | OCLNP1 | 132.54822 | 4.4842517 | 0.5573397 | 8.0458147 |
| 1.80E-13 | 1.03E-15 | ENSG00000227128.5 | LBX1-AS1 | 1246.9152 | 3.2240213 | 0.4018365 | 8.0232167 |
| 2.33E-13 | 1.34E-15 | ENSG00000174721.10 | FGFBP3 | 15618.637 | -1.862442 | 0.2330807 | -7.990546 |
| 2.36E-13 | 1.37E-15 | ENSG00000185813.11 | PCYT2 | 7470.0924 | -1.823875 | 0.2283227 | -7.988144 |
| 3.32E-13 | 1.94E-15 | ENSG00000135828.11 | RNASEL | 179.07589 | -3.880128 | 0.4883659 | -7.945126 |
| 4.36E-13 | 2.56E-15 | ENSG00000061455.11 | PRDM6 | 296.33583 | 2.8224092 | 0.3567893 | 7.9105781 |
| 5.23E-13 | 3.12E-15 | ENSG00000157219.5 | HTR5A | 407.09784 | -2.417351 | 0.3065312 | -7.88615 |
| 5.62E-13 | 3.38E-15 | ENSG00000082641.16 | NFE2L1 | 39832.056 | -1.583157 | 0.2010069 | -7.87613 |
| 5.77E-13 | 3.49E-15 | ENSG00000213719.8 | CLIC1 | 1553.137 | -2.699648 | 0.3429364 | -7.872155 |
| 6.72E-13 | 4.09E-15 | ENSG00000114115.10 | RBP1 | 32749.331 | -1.670972 | 0.212802 | -7.852239 |
| 7.16E-13 | 4.39E-15 | ENSG00000174453.10 | VWC2L | 854.8638 | 2.0316822 | 0.2590338 | 7.8433097 |
| 8.96E-13 | 5.53E-15 | ENSG00000089116.4 | LHX5 | 145.51401 | 4.228332 | 0.541099 | 7.8143413 |
| 9.84E-13 | 6.15E-15 | ENSG00000143340.6 | FAM163A | 3185.8811 | 1.7587939 | 0.2254603 | 7.800903 |
| 1.22E-12 | 7.69E-15 | ENSG00000257434.1 | AC073525.1 | 105.51804 | 4.6059711 | 0.59259 | 7.7726103 |
| 1.26E-12 | 7.95E-15 | ENSG00000150625.16 | GPM6A | 10340.434 | 2.4617537 | 0.3168956 | 7.7683433 |
| 1.32E-12 | 8.43E-15 | ENSG00000140450.9 | ARRDC4 | 3193.4012 | 1.5633238 | 0.2014328 | 7.7610183 |
| 1.32E-12 | 8.48E-15 | ENSG00000273906.1 | AC011297.1 | 118.99247 | -7.360159 | 0.9484568 | -7.760142 |
| 1.32E-12 | 8.51E-15 | ENSG00000169891.18 | REPS2 | 867.85182 | 2.0500418 | 0.2641869 | 7.7598166 |
| 1.40E-12 | 9.08E-15 | ENSG00000168490.14 | PHYHIP | 786.71041 | -2.046795 | 0.2640524 | -7.751474 |
| 1.40E-12 | 9.16E-15 | ENSG00000167508.12 | MVD | 24048.136 | -2.003571 | 0.2585122 | -7.750392 |
| 1.42E-12 | 9.35E-15 | ENSG00000125675.18 | GRIA3 | 2447.5541 | 1.8654337 | 0.24077 | 7.7477813 |
| 1.68E-12 | 1.12E-14 | ENSG00000109062.12 | SLC9A3R1 | 693.31093 | -2.060152 | 0.2666973 | -7.724683 |
| 1.80E-12 | 1.21E-14 | ENSG00000145075.13 | CCDC39 | 285.45951 | -11.67883 | 1.5138219 | -7.714801 |
| 2.31E-12 | 1.57E-14 | ENSG00000180787.6 | ZFP3 | 268.30313 | 2.5686438 | 0.3343726 | 7.6819808 |
| 2.53E-12 | 1.72E-14 | ENSG00000187140.6 | FOXD3 | 202.28176 | 4.8618345 | 0.6338841 | 7.6699113 |
| 2.93E-12 | 2.01E-14 | ENSG00000132109.10 | TRIM21 | 392.67365 | -2.26651 | 0.2962743 | -7.650041 |
| 2.95E-12 | 2.04E-14 | ENSG00000188620.11 | HMX3 | 1016.0711 | 2.2413673 | 0.2930559 | 7.6482595 |
| 2.98E-12 | 2.07E-14 | ENSG00000176884.15 | GRIN1 | 3801.2629 | 1.6835915 | 0.2201845 | 7.6462763 |

| | | | | | | | |
|---|---|---|---|---|---|---|---|
| 3.38E-12 | 2.36E-14 | ENSG00000169836.5 | TACR3 | 445.96298 | 2.2259815 | 0.2917668 | 7.6293182 |
| 3.39E-12 | 2.38E-14 | ENSG00000113805.8 | CNTN3 | 628.50246 | 2.8107861 | 0.3684826 | 7.6280016 |
| 3.39E-12 | 2.39E-14 | ENSG00000087076.9 | HSD17B14 | 296.77018 | -2.748311 | 0.3603091 | -7.62765 |
| 3.55E-12 | 2.54E-14 | ENSG00000235608.1 | NKX1-1 | 153.67144 | 3.5864989 | 0.4706693 | 7.6199971 |
| 3.60E-12 | 2.58E-14 | ENSG00000251664.5 | PCDHA12 | 4614.0669 | -2.218791 | 0.2912713 | -7.617609 |
| 4.18E-12 | 3.02E-14 | ENSG00000078725.13 | BRINP1 | 8427.8455 | 2.1686393 | 0.285436 | 7.5976383 |
| 4.56E-12 | 3.31E-14 | ENSG00000135406.14 | PRPH | 4542.5998 | -2.145377 | 0.2828258 | -7.585508 |
| 5.20E-12 | 3.80E-14 | ENSG00000131473.17 | ACLY | 82946.701 | -1.425572 | 0.1883769 | -7.567659 |
| 5.22E-12 | 3.83E-14 | ENSG00000171004.18 | HS6ST2 | 10223.587 | 1.5078793 | 0.1992801 | 7.5666337 |
| 5.47E-12 | 4.06E-14 | ENSG00000046651.15 | OFD1 | 1866.8951 | -1.852296 | 0.2450465 | -7.558957 |
| 6.65E-12 | 4.98E-14 | ENSG00000197324.9 | LRP10 | 955.35417 | -2.431883 | 0.3228537 | -7.532461 |
| 6.65E-12 | 4.99E-14 | ENSG00000112246.10 | SIM1 | 729.86602 | 6.7605328 | 0.897553 | 7.5321818 |
| 9.07E-12 | 6.84E-14 | ENSG00000110921.14 | MVK | 6168.9644 | -1.520435 | 0.2029724 | -7.490846 |
| 1.74E-11 | 1.35E-13 | ENSG00000181652.19 | ATG9B | 1088.8812 | -2.091164 | 0.2825436 | -7.401208 |
| 1.87E-11 | 1.46E-13 | ENSG00000168546.11 | GFRA2 | 3769.9438 | 2.1759465 | 0.2944148 | 7.3907513 |
| 1.95E-11 | 1.53E-13 | ENSG00000237975.7 | FLG-AS1 | 107.82873 | 6.6682613 | 0.9029716 | 7.384796 |
| 1.97E-11 | 1.55E-13 | ENSG00000100445.17 | SDR39U1 | 2201.5762 | 1.5463512 | 0.2094541 | 7.3827701 |
| 2.51E-11 | 2.01E-13 | ENSG00000285918.1 | AC092376.3 | 1748.5304 | 1.9677447 | 0.2677985 | 7.3478554 |
| 2.80E-11 | 2.26E-13 | ENSG00000215612.8 | HMX1 | 139.75612 | -3.431821 | 0.4680207 | -7.332627 |
| 2.87E-11 | 2.34E-13 | ENSG00000134824.14 | FADS2 | 64146.958 | -1.491039 | 0.2034819 | -7.327623 |
| 2.87E-11 | 2.35E-13 | ENSG00000169744.13 | LDB2 | 795.20402 | 1.9900049 | 0.2715988 | 7.3270031 |
| 2.95E-11 | 2.43E-13 | ENSG00000154678.18 | PDE1C | 9012.7805 | 2.7232336 | 0.3718819 | 7.3228448 |
| 3.73E-11 | 3.09E-13 | ENSG00000206053.13 | JPT2 | 11531.451 | -1.404908 | 0.1927015 | -7.290594 |
| 4.80E-11 | 4.01E-13 | ENSG00000138136.6 | LBX1 | 1587.3174 | 3.2039657 | 0.4416031 | 7.2553063 |
| 4.84E-11 | 4.06E-13 | ENSG00000099194.6 | SCD | 223194.68 | -1.567082 | 0.2160449 | -7.253504 |
| 4.84E-11 | 4.09E-13 | ENSG00000189337.17 | KAZN | 10117.905 | 1.4158 | 0.1952109 | 7.2526676 |
| 5.20E-11 | 4.41E-13 | ENSG00000183287.14 | CCBE1 | 393.46316 | 3.0717287 | 0.4241269 | 7.2424754 |
| 5.90E-11 | 5.03E-13 | ENSG00000273706.5 | LHX1 | 880.08948 | 2.1411415 | 0.2963714 | 7.2245215 |
| 7.01E-11 | 6.00E-13 | ENSG00000152931.9 | PART1 | 171.86165 | 3.8410174 | 0.5334427 | 7.2004309 |
| 7.06E-11 | 6.07E-13 | ENSG00000198597.9 | ZNF536 | 2641.2781 | 1.7243844 | 0.2395365 | 7.1988362 |
| 8.34E-11 | 7.21E-13 | ENSG00000229544.9 | NKX1-2 | 268.90087 | 2.4074956 | 0.3355185 | 7.175449 |
| 1.01E-10 | 8.80E-13 | ENSG00000106038.13 | EVX1 | 176.99201 | 10.852838 | 1.5182771 | 7.1481274 |
| 1.02E-10 | 8.95E-13 | ENSG00000169330.9 | MINAR1 | 5365.8633 | 1.4745539 | 0.206353 | 7.1457839 |
| 1.02E-10 | 8.97E-13 | ENSG00000184347.15 | SLIT3 | 16358.52 | -1.859817 | 0.2602795 | -7.145461 |
| 1.44E-10 | 1.28E-12 | ENSG00000011021.23 | CLCN6 | 18857.381 | -1.335438 | 0.1881716 | -7.096916 |
| 1.52E-10 | 1.36E-12 | ENSG00000111981.5 | ULBP1 | 362.99549 | -2.598752 | 0.3666162 | -7.088482 |
| 1.66E-10 | 1.48E-12 | ENSG00000147202.18 | DIAPH2 | 394.28047 | 2.0267764 | 0.2864322 | 7.0759366 |
| 1.82E-10 | 1.64E-12 | ENSG00000276550.4 | HERC2P2 | 5223.5746 | -1.888267 | 0.2673818 | -7.062063 |
| 1.88E-10 | 1.70E-12 | ENSG00000183662.11 | TAFA1 | 109.84848 | 4.3413201 | 0.6151862 | 7.0569204 |
| 3.69E-10 | 3.38E-12 | ENSG00000140022.13 | STON2 | 2520.5521 | -1.47492 | 0.2118881 | -6.960844 |
| 3.77E-10 | 3.47E-12 | ENSG00000164211.13 | STARD4 | 17327.737 | -1.879895 | 0.2702114 | -6.957126 |
| 3.93E-10 | 3.64E-12 | ENSG00000223865.11 | HLA-DPB1 | 242.83171 | -2.73298 | 0.3932034 | -6.950552 |
| 6.61E-10 | 6.22E-12 | ENSG00000120437.9 | ACAT2 | 38773.473 | -1.477964 | 0.2149927 | -6.874482 |
| 8.90E-10 | 8.47E-12 | ENSG00000100344.11 | PNPLA3 | 4436.1705 | -1.762582 | 0.2580485 | -6.830428 |
| 9.38E-10 | 8.98E-12 | ENSG00000163961.4 | RNF168 | 6283.9424 | -1.352677 | 0.1982841 | -6.821914 |
| 9.84E-10 | 9.46E-12 | ENSG00000102385.12 | DRP2 | 651.47962 | -1.859222 | 0.2728352 | -6.814448 |
| 1.01E-09 | 9.76E-12 | ENSG00000106948.16 | AKNA | 540.9785 | -2.097879 | 0.3080593 | -6.809982 |
| 1.02E-09 | 9.87E-12 | ENSG00000166006.14 | KCNC2 | 978.17588 | 2.5070441 | 0.36823 | 6.8083637 |
| 1.05E-09 | 1.02E-11 | ENSG00000166501.14 | PRKCB | 9341.4685 | 1.4253137 | 0.2095109 | 6.8030514 |
| 1.25E-09 | 1.22E-11 | ENSG00000162630.6 | B3GALT2 | 3991.036 | 1.4950158 | 0.220587 | 6.7774435 |
| 1.27E-09 | 1.24E-11 | ENSG00000133935.7 | ERG28 | 10249.259 | -1.595362 | 0.2354756 | -6.775064 |
| 1.28E-09 | 1.27E-11 | ENSG00000242779.7 | ZNF702P | 740.24882 | -1.980205 | 0.2923855 | -6.772581 |
| 1.29E-09 | 1.28E-11 | ENSG00000145416.13 | MARCHF1 | 9694.2606 | 1.3412928 | 0.1980944 | 6.7709784 |
| 1.43E-09 | 1.42E-11 | ENSG00000139292.13 | LGR5 | 397.03039 | 2.9013576 | 0.4294585 | 6.7558512 |
| 1.46E-09 | 1.46E-11 | ENSG00000163346.17 | PBXIP1 | 236.59756 | -2.577735 | 0.3817818 | -6.751854 |
| 1.46E-09 | 1.47E-11 | ENSG00000149782.11 | PLCB3 | 238.36497 | -2.65509 | 0.3933039 | -6.750733 |
| 1.46E-09 | 1.47E-11 | ENSG00000286381.1 | AL078622.1 | 113.88485 | 10.214955 | 1.5131682 | 6.750707 |
| 1.48E-09 | 1.50E-11 | ENSG00000095397.16 | WHRN | 394.70375 | -2.750311 | 0.4075727 | -6.748027 |
| 1.53E-09 | 1.56E-11 | ENSG00000172893.16 | DHCR7 | 17954.127 | -1.489624 | 0.2209464 | -6.742017 |

| | | | | | | | |
|---|---|---|---|---|---|---|---|
| 1.54E-09 | 1.58E-11 | ENSG00000157600.12 | TMEM164 | 4956.3157 | 1.3296798 | 0.1972617 | 6.7406894 |
| 1.88E-09 | 1.94E-11 | ENSG00000160285.15 | LSS | 28441.416 | -1.793934 | 0.2673306 | -6.710544 |
| 2.14E-09 | 2.21E-11 | ENSG00000165782.10 | PIP4P1 | 6131.7105 | -1.305103 | 0.195043 | -6.691358 |
| 2.19E-09 | 2.28E-11 | ENSG00000158270.12 | COLEC12 | 870.78592 | 1.8250555 | 0.2729245 | 6.6870346 |
| 2.25E-09 | 2.34E-11 | ENSG00000188158.15 | NHS | 4461.6273 | 1.5394975 | 0.2303685 | 6.6827616 |
| 2.28E-09 | 2.38E-11 | ENSG00000158195.11 | WASF2 | 3358.239 | -1.388631 | 0.2078705 | -6.680272 |
| 2.37E-09 | 2.49E-11 | ENSG00000125848.10 | FLRT3 | 4370.4089 | 1.3979371 | 0.2094624 | 6.6739292 |
| 2.48E-09 | 2.61E-11 | ENSG00000214575.9 | CPEB1 | 1170.8025 | -1.761623 | 0.2642332 | -6.666925 |
| 2.75E-09 | 2.93E-11 | ENSG00000118263.15 | KLF7 | 26790.634 | -1.240067 | 0.186468 | -6.650293 |
| 2.77E-09 | 2.96E-11 | ENSG00000147383.11 | NSDHL | 7367.1595 | -1.307042 | 0.1965858 | -6.648709 |
| 2.88E-09 | 3.10E-11 | ENSG00000185774.16 | KCNIP4 | 4843.2364 | -2.567858 | 0.3866199 | -6.641816 |
| 2.88E-09 | 3.11E-11 | ENSG00000183837.9 | PNMA3 | 3696.9127 | 1.3995391 | 0.2107325 | 6.6413078 |
| 2.88E-09 | 3.11E-11 | ENSG00000102924.12 | CBLN1 | 1188.1065 | 1.7361492 | 0.2614232 | 6.6411454 |
| 3.29E-09 | 3.58E-11 | ENSG00000140403.12 | DNAJA4 | 1288.292 | 1.8647466 | 0.2816604 | 6.6205492 |
| 3.79E-09 | 4.14E-11 | ENSG00000215252.11 | GOLGA8B | 15738.859 | 1.3017692 | 0.1972681 | 6.5989842 |
| 3.82E-09 | 4.18E-11 | ENSG00000046889.19 | PREX2 | 990.60264 | 2.4709206 | 0.3745252 | 6.5974756 |
| 4.04E-09 | 4.44E-11 | ENSG00000172037.14 | LAMB2 | 192.33046 | -3.862676 | 0.5862542 | -6.58874 |
| 4.69E-09 | 5.18E-11 | ENSG00000235934.2 | AC007405.3 | 126.86006 | 3.9575725 | 0.6027604 | 6.5657473 |
| 5.13E-09 | 5.72E-11 | ENSG00000261227.2 | AC140912.1 | 441.98998 | -1.80638 | 0.2757446 | -6.550919 |
| 5.43E-09 | 6.09E-11 | ENSG00000072571.20 | HMMR | 336.82383 | -2.478972 | 0.3789618 | -6.541482 |
| 5.43E-09 | 6.10E-11 | ENSG00000099250.18 | NRP1 | 12297.821 | -2.490679 | 0.3807635 | -6.541274 |
| 5.56E-09 | 6.29E-11 | ENSG00000186868.15 | MAPT | 36843.107 | 1.2684753 | 0.1940534 | 6.5367336 |
| 5.56E-09 | 6.29E-11 | ENSG00000010310.9 | GIPR | 195.66286 | -2.889596 | 0.4420567 | -6.53671 |
| 6.34E-09 | 7.20E-11 | ENSG00000288049.1 | AC010889.2 | 87.631379 | -9.974548 | 1.5306713 | -6.516453 |
| 6.53E-09 | 7.46E-11 | ENSG00000135549.15 | PKIB | 16576.967 | -1.683538 | 0.2585673 | -6.511025 |
| 6.91E-09 | 7.93E-11 | ENSG00000103018.17 | CYB5B | 22178.223 | -1.399079 | 0.2151813 | -6.50186 |
| 6.95E-09 | 8.00E-11 | ENSG00000172789.3 | HOXC5 | 3618.7423 | -2.114142 | 0.3252238 | -6.500577 |
| 7.20E-09 | 8.33E-11 | ENSG00000149972.11 | CNTN5 | 8013.8811 | 1.8113848 | 0.2789073 | 6.4945754 |
| 7.60E-09 | 8.81E-11 | ENSG00000115194.11 | SLC30A3 | 182.93963 | 2.6199157 | 0.4039303 | 6.4860584 |
| 8.22E-09 | 9.60E-11 | ENSG00000049246.14 | PER3 | 1793.7879 | -1.727783 | 0.2669173 | -6.473101 |
| 1.02E-08 | 1.20E-10 | ENSG00000256463.8 | SALL3 | 4121.838 | 1.6341753 | 0.253756 | 6.4399464 |
| 1.11E-08 | 1.32E-10 | ENSG00000205279.8 | CTXN3 | 1122.8829 | 1.9251811 | 0.2996293 | 6.4252089 |
| 1.20E-08 | 1.43E-10 | ENSG00000113327.16 | GABRG2 | 4964.5179 | 2.1409489 | 0.3338853 | 6.4122292 |
| 1.26E-08 | 1.51E-10 | ENSG00000168875.3 | SOX14 | 2131.7087 | 2.3559252 | 0.3678895 | 6.4038938 |
| 1.33E-08 | 1.60E-10 | ENSG00000067182.8 | TNFRSF1A | 594.5899 | -3.206254 | 0.5013422 | -6.39534 |
| 1.48E-08 | 1.80E-10 | ENSG00000184221.13 | OLIG1 | 114.0591 | -3.695228 | 0.5793883 | -6.377809 |
| 1.50E-08 | 1.83E-10 | ENSG00000121454.6 | LHX4 | 4747.8355 | 1.6227851 | 0.2545482 | 6.3751579 |
| 1.59E-08 | 1.94E-10 | ENSG00000140057.9 | AK7 | 227.71087 | -2.249948 | 0.3534274 | -6.366083 |
| 1.69E-08 | 2.07E-10 | ENSG00000105355.9 | PLIN3 | 392.25696 | -2.476743 | 0.3896597 | -6.35617 |
| 2.06E-08 | 2.54E-10 | ENSG00000145428.15 | RNF175 | 1383.3577 | 1.4408134 | 0.2278202 | 6.3243431 |
| 2.13E-08 | 2.64E-10 | ENSG00000122786.20 | CALD1 | 2416.402 | -1.452104 | 0.2298183 | -6.318488 |
| 2.36E-08 | 2.93E-10 | ENSG00000119655.11 | NPC2 | 1642.58 | -1.513756 | 0.2401785 | -6.302631 |
| 2.37E-08 | 2.95E-10 | ENSG00000117602.12 | RCAN3 | 5490.3944 | -1.750086 | 0.2777381 | -6.301209 |
| 2.55E-08 | 3.20E-10 | ENSG00000113430.10 | IRX4 | 169.80362 | 4.1769883 | 0.6641794 | 6.2889462 |
| 3.10E-08 | 3.91E-10 | ENSG00000133101.10 | CCNA1 | 1270.6043 | -1.851526 | 0.2958815 | -6.257662 |
| 3.13E-08 | 3.98E-10 | ENSG00000128645.15 | HOXD1 | 1413.1944 | -2.059373 | 0.3292368 | -6.254991 |
| 3.24E-08 | 4.12E-10 | ENSG00000145934.16 | TENM2 | 7925.8251 | 2.145681 | 0.3433477 | 6.249294 |
| 3.43E-08 | 4.38E-10 | ENSG00000263146.2 | LINC01896 | 1905.6794 | 1.7545196 | 0.2811785 | 6.2398772 |
| 3.71E-08 | 4.75E-10 | ENSG00000130164.14 | LDLR | 19225.386 | -1.333942 | 0.2142189 | -6.227006 |
| 3.71E-08 | 4.77E-10 | ENSG00000074317.11 | SNCB | 1798.7201 | 1.3895441 | 0.2231701 | 6.2263895 |
| 3.99E-08 | 5.15E-10 | ENSG00000006756.16 | ARSD | 1884.4955 | 1.2945399 | 0.2083137 | 6.2143779 |
| 4.14E-08 | 5.36E-10 | ENSG00000265972.6 | TXNIP | 6119.2287 | 1.8766232 | 0.3022859 | 6.2081071 |
| 4.15E-08 | 5.39E-10 | ENSG00000240875.6 | LINC00886 | 86.313475 | 3.5442673 | 0.5709845 | 6.207292 |
| 4.52E-08 | 5.89E-10 | ENSG00000173801.17 | JUP | 9444.3874 | -1.569782 | 0.25346 | -6.193413 |
| 5.03E-08 | 6.61E-10 | ENSG00000175265.17 | GOLGA8A | 15134.704 | 1.1705354 | 0.1895529 | 6.175245 |
| 5.52E-08 | 7.27E-10 | ENSG00000133107.15 | TRPC4 | 384.22894 | 2.0520265 | 0.3331177 | 6.1600639 |
| 5.53E-08 | 7.32E-10 | ENSG00000182742.6 | HOXB4 | 6700.4411 | -1.289413 | 0.2093525 | -6.159051 |
| 5.94E-08 | 7.88E-10 | ENSG00000134324.12 | LPIN1 | 12041.289 | -1.271963 | 0.2069156 | -6.147254 |
| 6.03E-08 | 8.02E-10 | ENSG00000162989.5 | KCNJ3 | 1151.9545 | 2.5611744 | 0.416827 | 6.1444546 |

| | | | | | | | |
|---|---|---|---|---|---|---|---|
| 6.12E-08 | 8.16E-10 | ENSG00000128578.10 | STRIP2 | 1966.0813 | 1.3198024 | 0.2148905 | 6.1417447 |
| 6.19E-08 | 8.28E-10 | ENSG00000120594.17 | PLXDC2 | 2566.6134 | 1.6679754 | 0.2716814 | 6.1394533 |
| 6.70E-08 | 9.01E-10 | ENSG00000174669.12 | SLC29A2 | 1597.2483 | -1.482751 | 0.2420446 | -6.125941 |
| 6.99E-08 | 9.44E-10 | ENSG00000178538.10 | CA8 | 283.79524 | 2.0021854 | 0.3272286 | 6.1186144 |
| 7.09E-08 | 9.61E-10 | ENSG00000186094.17 | AGBL4 | 486.7933 | 1.7955971 | 0.2936014 | 6.1157644 |
| 8.12E-08 | 1.11E-09 | ENSG00000176728.9 | TTTY14 | 110.80324 | -8.870256 | 1.4558894 | -6.092671 |
| 8.72E-08 | 1.20E-09 | ENSG00000206159.11 | GYG2P1 | 62.712587 | -9.491721 | 1.5609402 | -6.080772 |
| 8.76E-08 | 1.21E-09 | ENSG00000168779.19 | SHOX2 | 318.56489 | 2.8922315 | 0.4757308 | 6.0795545 |
| 9.76E-08 | 1.35E-09 | ENSG00000156298.12 | TSPAN7 | 24924.431 | 1.1426659 | 0.1885157 | 6.0613844 |
| 1.03E-07 | 1.42E-09 | ENSG00000267034.1 | AC010980.2 | 820.23877 | 1.4701697 | 0.242885 | 6.0529467 |
| 1.09E-07 | 1.51E-09 | ENSG00000188242.4 | AC010442.1 | 69.216347 | 4.0632578 | 0.6723911 | 6.042998 |
| 1.18E-07 | 1.66E-09 | ENSG00000189114.8 | BLOC1S3 | 1169.7689 | -1.594875 | 0.2645888 | -6.027751 |
| 1.26E-07 | 1.78E-09 | ENSG00000139793.18 | MBNL2 | 1301.0176 | -2.029786 | 0.3373499 | -6.016856 |
| 1.32E-07 | 1.86E-09 | ENSG00000065534.18 | MYLK | 798.34404 | -2.15699 | 0.3589477 | -6.009205 |
| 1.38E-07 | 1.96E-09 | ENSG00000198468.9 | FLVCR1-DT | 582.38481 | -1.524747 | 0.2540885 | -6.000849 |
| 1.52E-07 | 2.17E-09 | ENSG00000166897.15 | ELFN2 | 2410.153 | 1.5319778 | 0.2559878 | 5.9845744 |
| 1.66E-07 | 2.37E-09 | ENSG00000240240.9 | BX664727.3 | 216.56806 | 2.2683906 | 0.3799545 | 5.970164 |
| 1.79E-07 | 2.57E-09 | ENSG00000077264.15 | PAK3 | 50373.59 | 1.4733467 | 0.2473378 | 5.9568203 |
| 1.90E-07 | 2.74E-09 | ENSG00000139289.13 | PHLDA1 | 11708.006 | 1.690691 | 0.284314 | 5.9465636 |
| 1.91E-07 | 2.77E-09 | ENSG00000104313.20 | EYA1 | 808.29541 | 2.2969749 | 0.3863973 | 5.9445939 |
| 1.96E-07 | 2.85E-09 | ENSG00000141458.13 | NPC1 | 5837.1348 | -1.443059 | 0.2429359 | -5.940079 |
| 2.21E-07 | 3.23E-09 | ENSG00000114374.13 | USP9Y | 11176.411 | -8.706955 | 1.4708414 | -5.919711 |
| 2.26E-07 | 3.31E-09 | ENSG00000002586.20 | CD99 | 655.29739 | -1.481445 | 0.2504368 | -5.915444 |
| 2.30E-07 | 3.38E-09 | ENSG00000143816.8 | WNT9A | 646.95227 | -1.511314 | 0.2556408 | -5.911865 |
| 2.48E-07 | 3.66E-09 | ENSG00000002586.20_PA | CD99 | 655.69726 | -1.478062 | 0.2505686 | -5.898834 |
| 2.56E-07 | 3.79E-09 | ENSG00000132821.12 | VSTM2L | 8612.6373 | 1.6387747 | 0.2780762 | 5.8932572 |
| 2.56E-07 | 3.81E-09 | ENSG00000105939.13 | ZC3HAV1 | 1801.8206 | -1.748887 | 0.2968027 | -5.892423 |
| 2.66E-07 | 3.96E-09 | ENSG00000116016.14 | EPAS1 | 70.292953 | 9.5156338 | 1.6166773 | 5.8859203 |
| 2.74E-07 | 4.08E-09 | ENSG00000176771.17 | NCKAP5 | 1848.382 | -1.924602 | 0.3272685 | -5.880806 |
| 3.09E-07 | 4.64E-09 | ENSG00000150275.18 | PCDH15 | 3331.7969 | 1.9917565 | 0.3399079 | 5.8596939 |
| 3.33E-07 | 5.03E-09 | ENSG00000261146.1 | AC007159.1 | 93.948843 | 3.8213283 | 0.6536443 | 5.8461892 |
| 3.47E-07 | 5.27E-09 | ENSG00000185869.14 | ZNF829 | 291.32984 | 1.9606803 | 0.335816 | 5.8385558 |
| 3.67E-07 | 5.60E-09 | ENSG00000128272.14 | ATF4 | 20125.355 | -1.282178 | 0.2199952 | -5.828211 |
| 4.18E-07 | 6.39E-09 | ENSG00000254858.10 | MPV17L2 | 1467.3794 | -1.394231 | 0.2401221 | -5.80634 |
| 4.22E-07 | 6.47E-09 | ENSG00000260197.1 | AC010889.1 | 56.390687 | -9.33952 | 1.6091388 | -5.804049 |
| 4.66E-07 | 7.16E-09 | ENSG00000147234.10 | FRMPD3 | 4196.0162 | 1.3004703 | 0.2247208 | 5.787049 |
| 4.90E-07 | 7.55E-09 | ENSG00000182263.14 | FIGN | 2340.4656 | 1.3795356 | 0.2387485 | 5.7781966 |
| 5.29E-07 | 8.18E-09 | ENSG00000178573.7 | MAF | 4941.0103 | 1.2786168 | 0.221803 | 5.7646517 |
| 5.38E-07 | 8.34E-09 | ENSG00000099251.14 | HSD17B7P2 | 441.7063 | 1.746346 | 0.3031045 | 5.7615304 |
| 5.55E-07 | 8.65E-09 | ENSG00000160145.15 | KALRN | 20035.303 | -1.164278 | 0.2022959 | -5.75532 |
| 6.03E-07 | 9.42E-09 | ENSG00000155368.16 | DBI | 5160.1619 | -1.366823 | 0.2380853 | -5.740897 |
| 6.62E-07 | 1.04E-08 | ENSG00000058866.15 | DGKG | 97.173168 | 2.8722314 | 0.5017391 | 5.7245517 |
| 6.69E-07 | 1.05E-08 | ENSG00000143545.10 | RAB13 | 107.50653 | -3.378829 | 0.5904553 | -5.722413 |
| 6.77E-07 | 1.07E-08 | ENSG00000133710.16 | SPINK5 | 361.88321 | 2.8571393 | 0.4995062 | 5.7199275 |
| 6.77E-07 | 1.07E-08 | ENSG00000151929.10 | BAG3 | 792.20685 | -1.654514 | 0.2892791 | -5.719438 |
| 6.79E-07 | 1.07E-08 | ENSG00000234969.1 | AL627389.1 | 50.578645 | 9.0443695 | 1.5815552 | 5.7186556 |
| 6.84E-07 | 1.09E-08 | ENSG00000273301.1 | AC016717.2 | 612.29392 | -1.792212 | 0.3134977 | -5.716826 |
| 7.01E-07 | 1.11E-08 | ENSG00000137968.16 | SLC44A5 | 10436.594 | -1.426844 | 0.2497861 | -5.712264 |
| 7.21E-07 | 1.15E-08 | ENSG00000075035.10 | WSCD2 | 2461.7661 | 1.2309583 | 0.2156858 | 5.7071838 |
| 8.25E-07 | 1.32E-08 | ENSG00000139219.19 | COL2A1 | 257.46947 | -3.29673 | 0.5801173 | -5.682868 |
| 8.65E-07 | 1.40E-08 | ENSG00000272163.1 | AF106564.1 | 3054.0213 | 1.3825634 | 0.2436732 | 5.673842 |
| 8.65E-07 | 1.40E-08 | ENSG00000057704.13 | TMCC3 | 597.77431 | 2.3289132 | 0.4104685 | 5.6737933 |
| 8.85E-07 | 1.44E-08 | ENSG00000253690.1 | AC021678.2 | 48.896772 | 8.995971 | 1.5868805 | 5.6689657 |
| 9.06E-07 | 1.47E-08 | ENSG00000279375.1 | AC244517.7 | 135.35996 | -2.349639 | 0.4147925 | -5.664614 |
| 9.20E-07 | 1.50E-08 | ENSG00000186235.11 | LINC02610 | 172.16157 | 2.4248245 | 0.4282959 | 5.6615635 |
| 1.09E-06 | 1.78E-08 | ENSG00000198914.5 | POU3F3 | 1483.3828 | 1.3308984 | 0.2363004 | 5.6322313 |
| 1.14E-06 | 1.88E-08 | ENSG00000177453.7 | NIM1K | 392.95713 | 1.7664774 | 0.3141537 | 5.6229715 |
| 1.16E-06 | 1.91E-08 | ENSG00000182134.16 | TDRKH | 5864.3654 | -1.482102 | 0.2637304 | -5.619761 |
| 1.18E-06 | 1.95E-08 | ENSG00000107187.17 | LHX3 | 356.30369 | 2.8293445 | 0.5037699 | 5.6163426 |

| | | | | | | | |
|---|---|---|---|---|---|---|---|
| 1.31E-06 | 2.17E-08 | ENSG00000085719.13 | CPNE3 | 5810.0716 | -1.128345 | 0.2015542 | -5.598221 |
| 1.34E-06 | 2.22E-08 | ENSG00000100441.10 | KHNYN | 2791.5584 | -1.363701 | 0.2437845 | -5.593879 |
| 1.37E-06 | 2.28E-08 | ENSG00000170419.10 | VSTM2A | 5804.6838 | 1.5122288 | 0.2705531 | 5.5893969 |
| 1.43E-06 | 2.39E-08 | ENSG00000250579.2 | AC022424.1 | 700.62393 | 1.9421383 | 0.3479709 | 5.5813235 |
| 1.45E-06 | 2.44E-08 | ENSG00000111684.11 | LPCAT3 | 2558.062 | -1.249653 | 0.2240442 | -5.577706 |
| 1.48E-06 | 2.50E-08 | ENSG00000153208.17 | MERTK | 216.5246 | 1.9768129 | 0.3546902 | 5.57335 |
| 1.49E-06 | 2.53E-08 | ENSG00000169432.18 | SCN9A | 14032.996 | -1.969862 | 0.3535719 | -5.571321 |
| 1.49E-06 | 2.53E-08 | ENSG00000112531.17 | QKI | 11913.587 | -1.267079 | 0.2274305 | -5.57128 |
| 1.51E-06 | 2.56E-08 | ENSG00000155366.17 | RHOC | 2151.8366 | -1.835438 | 0.3295838 | -5.568956 |
| 1.57E-06 | 2.67E-08 | ENSG00000165810.17 | BTNL9 | 1637.8988 | 1.7658538 | 0.3175107 | 5.5615569 |
| 1.62E-06 | 2.77E-08 | ENSG00000215790.7 | SLC35E2A | 22355.975 | 1.0479837 | 0.1886345 | 5.5556315 |
| 1.72E-06 | 2.96E-08 | ENSG00000165507.9 | DEPP1 | 374.19868 | -1.790755 | 0.3230046 | -5.544055 |
| 1.75E-06 | 3.02E-08 | ENSG00000160752.14 | FDPS | 16598.502 | -1.047918 | 0.1891448 | -5.540297 |
| 1.77E-06 | 3.06E-08 | ENSG00000122694.16 | GLIPR2 | 612.76929 | -1.748027 | 0.3156401 | -5.538038 |
| 1.80E-06 | 3.12E-08 | ENSG00000182199.11 | SHMT2 | 3800.3882 | -1.119172 | 0.2022141 | -5.53459 |
| 1.83E-06 | 3.17E-08 | ENSG00000133789.15 | SWAP70 | 1150.5134 | -1.239785 | 0.2241236 | -5.531703 |
| 1.90E-06 | 3.30E-08 | ENSG00000285953.1 | AC000120.4 | 47.439795 | -9.088403 | 1.6450735 | -5.524618 |
| 1.98E-06 | 3.45E-08 | ENSG00000183783.7 | KCTD8 | 835.29501 | 1.7539523 | 0.3179345 | 5.5167097 |
| 1.99E-06 | 3.48E-08 | ENSG00000077943.8 | ITGA8 | 416.19569 | 2.596775 | 0.4708295 | 5.5153191 |
| 2.40E-06 | 4.23E-08 | ENSG00000286431.1 | BX842570.1 | 555.47819 | 1.5967265 | 0.2913125 | 5.4811464 |
| 2.44E-06 | 4.30E-08 | ENSG00000155792.10 | DEPTOR | 487.64005 | 1.6803975 | 0.3067576 | 5.4779331 |
| 2.44E-06 | 4.31E-08 | ENSG00000008128.23 | CDK11A | 1755.5456 | 1.1623459 | 0.212194 | 5.4777511 |
| 2.44E-06 | 4.32E-08 | ENSG00000233608.4 | TWIST2 | 47.874202 | 5.8976258 | 1.0767762 | 5.4771138 |
| 2.50E-06 | 4.44E-08 | ENSG00000134207.16 | SYT6 | 1523.6156 | 1.5990636 | 0.2922017 | 5.4724654 |
| 2.50E-06 | 4.46E-08 | ENSG00000250360.1 | AC008808.2 | 149.55317 | -2.21844 | 0.4054455 | -5.47161 |
| 2.70E-06 | 4.83E-08 | ENSG00000174705.13 | SH3PXD2B | 11258.62 | -1.244384 | 0.2280175 | -5.457407 |
| 2.79E-06 | 4.99E-08 | ENSG00000166402.9 | TUB | 7763.0712 | 1.1301685 | 0.2073124 | 5.4515235 |
| 2.85E-06 | 5.13E-08 | ENSG00000286945.1 | AL513478.4 | 131.54579 | 2.3765814 | 0.4363218 | 5.4468541 |
| 2.88E-06 | 5.19E-08 | ENSG00000122483.17 | CCDC18 | 542.05321 | 1.4111793 | 0.2591895 | 5.4445853 |
| 2.95E-06 | 5.33E-08 | ENSG00000180537.13 | RNF182 | 735.44807 | -1.561353 | 0.2870082 | -5.4401 |
| 2.95E-06 | 5.34E-08 | ENSG00000188981.11 | MSANTD1 | 126.25896 | -3.114769 | 0.5725995 | -5.439699 |
| 3.03E-06 | 5.50E-08 | ENSG00000107147.13 | KCNT1 | 391.69399 | 1.7776852 | 0.3271252 | 5.4342656 |
| 3.03E-06 | 5.50E-08 | ENSG00000154620.6 | TMSB4Y | 372.61029 | -8.45294 | 1.5554891 | -5.434265 |
| 3.05E-06 | 5.55E-08 | ENSG00000178401.16 | DNAJC22 | 1683.5003 | 1.6334213 | 0.3006607 | 5.4327739 |
| 3.24E-06 | 5.92E-08 | ENSG00000107819.13 | SFXN3 | 8850.5818 | -1.172605 | 0.2162999 | -5.4212 |
| 3.34E-06 | 6.13E-08 | ENSG00000243978.8 | RTL9 | 827.10682 | 1.3980876 | 0.2581842 | 5.4150788 |
| 3.37E-06 | 6.19E-08 | ENSG00000226752.9 | CUTALP | 3673.018 | 1.1071138 | 0.2045169 | 5.4133122 |
| 3.38E-06 | 6.22E-08 | ENSG00000120251.20 | GRIA2 | 12247.757 | 1.0814702 | 0.1998177 | 5.4122844 |
| 3.40E-06 | 6.28E-08 | ENSG00000157168.20 | NRG1 | 18316.483 | -1.442994 | 0.2667014 | -5.410525 |
| 3.64E-06 | 6.73E-08 | ENSG00000172828.13 | CES3 | 161.7539 | -2.503374 | 0.4637325 | -5.398314 |
| 3.69E-06 | 6.84E-08 | ENSG00000170312.16 | CDK1 | 51.07948 | -9.194994 | 1.7042575 | -5.395308 |
| 3.90E-06 | 7.26E-08 | ENSG00000112309.11 | B3GAT2 | 6978.279 | 1.0529493 | 0.1955502 | 5.3845468 |
| 3.93E-06 | 7.33E-08 | ENSG00000047230.15 | CTPS2 | 5175.5859 | 1.1338403 | 0.2106414 | 5.3828001 |
| 3.93E-06 | 7.34E-08 | ENSG00000173599.15 | PC | 1748.7918 | -1.273392 | 0.2365774 | -5.382561 |
| 4.19E-06 | 7.85E-08 | ENSG00000120306.11 | CYSTM1 | 1064.7053 | -1.477542 | 0.2751148 | -5.370638 |
| 4.21E-06 | 7.91E-08 | ENSG00000077616.11 | NAALAD2 | 1584.5827 | 1.8429329 | 0.3432436 | 5.3691693 |
| 4.26E-06 | 8.02E-08 | ENSG00000169118.18 | CSNK1G1 | 4454.5176 | -1.048717 | 0.1954132 | -5.366667 |
| 4.69E-06 | 8.84E-08 | ENSG00000249690.1 | AC110813.1 | 38.97838 | 8.6669224 | 1.6202868 | 5.3490052 |
| 4.89E-06 | 9.24E-08 | ENSG00000079459.13 | FDFT1 | 26192.155 | -1.149398 | 0.2152009 | -5.341044 |
| 5.72E-06 | 1.09E-07 | ENSG00000011105.14 | TSPAN9 | 3659.3335 | 1.3063298 | 0.2459346 | 5.311696 |
| 6.04E-06 | 1.15E-07 | ENSG00000286353.1 | AC115282.1 | 141.486 | 3.1798332 | 0.5998219 | 5.3012959 |
| 6.14E-06 | 1.17E-07 | ENSG00000135269.18 | TES | 6175.5048 | -1.023208 | 0.1931415 | -5.29771 |
| 6.22E-06 | 1.19E-07 | ENSG00000013293.6 | SLC7A14 | 6101.2523 | 1.3610235 | 0.2570375 | 5.2950391 |
| 6.75E-06 | 1.30E-07 | ENSG00000124191.18 | TOX2 | 1106.5803 | -1.457216 | 0.2760116 | -5.279547 |
| 6.88E-06 | 1.32E-07 | ENSG00000196482.18 | ESRRG | 2510.5094 | 1.3631696 | 0.2583927 | 5.2755726 |
| 6.90E-06 | 1.33E-07 | ENSG00000260691.6 | ANKRD20A1 | 219.75001 | 2.0456724 | 0.3878187 | 5.274816 |
| 6.91E-06 | 1.33E-07 | ENSG00000264589.4 | MAPT-AS1 | 131.95852 | 2.5096518 | 0.4758456 | 5.2740883 |
| 6.94E-06 | 1.34E-07 | ENSG00000136010.14 | ALDH1L2 | 3550.8116 | -1.438758 | 0.2728539 | -5.272995 |
| 6.96E-06 | 1.35E-07 | ENSG00000171094.18 | ALK | 2427.687 | 1.7191172 | 0.3260844 | 5.2720001 |

| | | | | | | | |
|---|---|---|---|---|---|---|---|
| 7.05E-06 | 1.37E-07 | ENSG00000006047.13 | YBX2 | 336.71005 | -1.70374 | 0.3233323 | -5.269316 |
| 7.19E-06 | 1.40E-07 | ENSG00000026025.16 | VIM | 12052.148 | -1.923256 | 0.3652756 | -5.26522 |
| 7.68E-06 | 1.50E-07 | ENSG00000177000.13 | MTHFR | 8447.7916 | -1.069514 | 0.203622 | -5.252448 |
| 7.68E-06 | 1.51E-07 | ENSG00000087245.13 | MMP2 | 2944.6393 | -1.188327 | 0.2262649 | -5.251929 |
| 8.08E-06 | 1.59E-07 | ENSG00000253405.1 | EVX1-AS | 37.847059 | 8.6284923 | 1.6459198 | 5.2423529 |
| 8.21E-06 | 1.62E-07 | ENSG00000172014.12 | ANKRD20A4 | 152.21014 | 2.7036253 | 0.5160745 | 5.2388276 |
| 8.62E-06 | 1.70E-07 | ENSG00000119614.3 | VSX2 | 4099.8065 | 1.561938 | 0.2986777 | 5.22951 |
| 8.78E-06 | 1.73E-07 | ENSG00000155090.15 | KLF10 | 4294.3565 | -1.188604 | 0.2274515 | -5.225746 |
| 8.81E-06 | 1.74E-07 | ENSG00000198496.12 | NBR2 | 366.93844 | -1.681654 | 0.3218585 | -5.224823 |
| 8.84E-06 | 1.75E-07 | ENSG00000182132.13 | KCNIP1 | 4890.4802 | 1.4492091 | 0.2774297 | 5.2236987 |
| 9.38E-06 | 1.87E-07 | ENSG00000241489.8 | AC244197.3 | 1367.4741 | 2.2555764 | 0.4327944 | 5.2116584 |
| 9.38E-06 | 1.88E-07 | ENSG00000230552.6 | AC092162.2 | 102.88764 | 2.642781 | 0.507128 | 5.2112703 |
| 9.48E-06 | 1.90E-07 | ENSG00000124343.13 | XG | 45.1325 | 4.1997281 | 0.8063142 | 5.20855 |
| 9.56E-06 | 1.92E-07 | ENSG00000109906.14 | ZBTB16 | 1515.6689 | 1.791648 | 0.3441078 | 5.2066474 |
| 9.59E-06 | 1.93E-07 | ENSG00000241839.10 | PLEKHO2 | 678.67616 | -1.729271 | 0.3321876 | -5.205706 |
| 9.79E-06 | 1.98E-07 | ENSG00000197576.14 | HOXA4 | 4879.5079 | -1.20144 | 0.2309788 | -5.201514 |
| 9.85E-06 | 1.99E-07 | ENSG00000092377.15 | TBL1Y | 33.872926 | -8.602763 | 1.6544132 | -5.199888 |
| 9.86E-06 | 2.00E-07 | ENSG00000223768.2 | LINC00205 | 9077.3069 | -1.060929 | 0.2040486 | -5.199392 |
| 1.03E-05 | 2.09E-07 | ENSG00000166839.17 | ANKDD1A | 919.79619 | -1.227365 | 0.2364233 | -5.191387 |
| 1.06E-05 | 2.16E-07 | ENSG00000100101.15 | Z83844.1 | 893.51799 | -1.29503 | 0.2497807 | -5.184668 |
| 1.10E-05 | 2.25E-07 | ENSG00000089472.16 | HEPH | 1362.0611 | -1.570401 | 0.3033014 | -5.177691 |
| 1.14E-05 | 2.33E-07 | ENSG00000187079.19 | TEAD1 | 2106.8461 | -1.06535 | 0.2060417 | -5.170556 |
| 1.21E-05 | 2.49E-07 | ENSG00000129566.13 | TEP1 | 802.38394 | -1.466069 | 0.284189 | -5.158781 |
| 1.22E-05 | 2.52E-07 | ENSG00000285188.1 | AC008397.2 | 715.28929 | -1.593377 | 0.3090397 | -5.155897 |
| 1.30E-05 | 2.69E-07 | ENSG00000146938.16 | NLGN4X | 12452.299 | 1.153327 | 0.2242107 | 5.1439417 |
| 1.35E-05 | 2.79E-07 | ENSG00000168672.4 | LRATD2 | 1324.8206 | 1.8841413 | 0.3667906 | 5.1368306 |
| 1.39E-05 | 2.88E-07 | ENSG00000140682.19 | TGFB1I1 | 218.3468 | -2.270758 | 0.4425558 | -5.131009 |
| 1.41E-05 | 2.94E-07 | ENSG00000151229.13 | SLC2A13 | 1148.653 | 1.3153043 | 0.2565377 | 5.1271378 |
| 1.43E-05 | 3.00E-07 | ENSG00000147852.16 | VLDLR | 3914.4236 | -1.067866 | 0.2084239 | -5.12353 |
| 1.43E-05 | 3.00E-07 | ENSG00000070748.19 | CHAT | 3185.0596 | -1.277522 | 0.2493586 | -5.123234 |
| 1.46E-05 | 3.06E-07 | ENSG00000143013.13 | LMO4 | 5660.7417 | 1.0200917 | 0.1992572 | 5.1194715 |
| 1.50E-05 | 3.14E-07 | ENSG00000186019.11 | AC021092.1 | 3011.0772 | 1.4308758 | 0.2797609 | 5.1146374 |
| 1.50E-05 | 3.16E-07 | ENSG00000115423.19 | DNAH6 | 560.56482 | -1.668996 | 0.3263945 | -5.113432 |
| 1.50E-05 | 3.17E-07 | ENSG00000101843.19 | PSMD10 | 5398.0675 | 1.0511721 | 0.2055854 | 5.1130675 |
| 1.55E-05 | 3.27E-07 | ENSG00000106006.6 | HOXA6 | 849.16354 | 1.9787325 | 0.3874312 | 5.1073128 |
| 1.57E-05 | 3.32E-07 | ENSG00000230453.9 | ANKRD18B | 110.68185 | 2.5085814 | 0.4914415 | 5.1045371 |
| 1.59E-05 | 3.39E-07 | ENSG00000257935.2 | LHX5-AS1 | 34.885199 | 8.5113363 | 1.6687293 | 5.1004895 |
| 1.63E-05 | 3.48E-07 | ENSG00000013573.17 | DDX11 | 1073.1444 | 1.3410025 | 0.263177 | 5.09544 |
| 1.71E-05 | 3.66E-07 | ENSG00000253661.2 | ZFHX4-AS1 | 275.1476 | 1.8409969 | 0.361976 | 5.0859645 |
| 1.71E-05 | 3.67E-07 | ENSG00000106113.19 | CRHR2 | 131.30413 | 2.7239842 | 0.5356767 | 5.085127 |
| 1.73E-05 | 3.72E-07 | ENSG00000157087.20 | ATP2B2 | 2225.3841 | 1.5701079 | 0.3089124 | 5.082696 |
| 1.75E-05 | 3.78E-07 | ENSG00000139178.11 | C1RL | 161.03735 | -2.652645 | 0.5221953 | -5.079796 |
| 1.77E-05 | 3.83E-07 | ENSG00000186193.9 | SAPCD2 | 350.28821 | -1.641865 | 0.3233705 | -5.07735 |
| 1.77E-05 | 3.84E-07 | ENSG00000198105.15 | ZNF248 | 7674.6237 | 1.2743488 | 0.2510061 | 5.0769632 |
| 1.93E-05 | 4.20E-07 | ENSG00000152910.19 | CNTNAP4 | 556.41562 | 1.7145289 | 0.3388705 | 5.0595398 |
| 1.94E-05 | 4.22E-07 | ENSG00000128253.15 | RFPL2 | 580.95059 | 1.3304402 | 0.262987 | 5.0589577 |
| 1.96E-05 | 4.28E-07 | ENSG00000067082.15 | KLF6 | 8290.0019 | -1.139537 | 0.225381 | -5.056047 |
| 1.98E-05 | 4.34E-07 | ENSG00000164294.14 | GPX8 | 57.0332 | -6.216175 | 1.2300744 | -5.053495 |
| 1.98E-05 | 4.35E-07 | ENSG00000110693.18 | SOX6 | 938.25933 | -2.182847 | 0.4320023 | -5.05286 |
| 2.02E-05 | 4.45E-07 | ENSG00000112972.15 | HMGCS1 | 164398.56 | -1.199652 | 0.2376251 | -5.048509 |
| 2.07E-05 | 4.57E-07 | ENSG00000183044.12 | ABAT | 18077.356 | 1.0071245 | 0.1996798 | 5.0436982 |
| 2.14E-05 | 4.75E-07 | ENSG00000249896.2 | LINC02495 | 527.41564 | 1.4087705 | 0.2797219 | 5.0363256 |
| 2.18E-05 | 4.84E-07 | ENSG00000259363.6 | AC090825.1 | 578.24367 | -1.611799 | 0.3202873 | -5.032353 |
| 2.19E-05 | 4.89E-07 | ENSG00000147421.18 | HMBOX1 | 2654.4756 | -1.130921 | 0.2248029 | -5.030721 |
| 2.27E-05 | 5.10E-07 | ENSG00000001460.18 | STPG1 | 528.05209 | -1.472868 | 0.2932491 | -5.022585 |
| 2.27E-05 | 5.10E-07 | ENSG00000165084.16 | C8orf34 | 1081.7166 | 1.3655047 | 0.271878 | 5.0224906 |
| 2.27E-05 | 5.10E-07 | ENSG00000135069.14 | PSAT1 | 3369.6485 | -1.127382 | 0.2244678 | -5.022467 |
| 2.31E-05 | 5.19E-07 | ENSG00000171444.18 | MCC | 1488.0824 | -1.146482 | 0.2284289 | -5.018987 |
| 2.36E-05 | 5.34E-07 | ENSG00000147255.19 | IGSF1 | 1998.5306 | 1.1847335 | 0.2362945 | 5.0138008 |

| | | | | | | | |
|---|---|---|---|---|---|---|---|
| 2.37E-05 | 5.37E-07 | ENSG00000104419.16 | NDRG1 | 1239.5072 | -1.500569 | 0.2993522 | -5.012721 |
| 2.44E-05 | 5.54E-07 | ENSG00000155849.15 | ELMO1 | 5466.5972 | 1.0430352 | 0.2083283 | 5.006689 |
| 2.49E-05 | 5.66E-07 | ENSG00000283005.1 | CDC27P3 | 59.609398 | 3.2454364 | 0.6487623 | 5.0025044 |
| 2.60E-05 | 5.95E-07 | ENSG00000111783.12 | RFX4 | 49.966662 | -5.485973 | 1.0987485 | -4.992929 |
| 2.66E-05 | 6.12E-07 | ENSG00000169710.9 | FASN | 71820.973 | -1.339968 | 0.2686732 | -4.987353 |
| 2.80E-05 | 6.46E-07 | ENSG00000261722.1 | AC092376.1 | 293.75559 | 1.675736 | 0.3366949 | 4.9770166 |
| 2.95E-05 | 6.82E-07 | ENSG00000198680.4 | TUSC1 | 936.15611 | 1.3796014 | 0.2777885 | 4.966373 |
| 3.24E-05 | 7.54E-07 | ENSG00000117266.15 | CDK18 | 202.02873 | 1.9157303 | 0.3872642 | 4.9468301 |
| 3.30E-05 | 7.71E-07 | ENSG00000123685.9 | BATF3 | 151.43568 | -3.041971 | 0.6154763 | -4.942466 |
| 3.36E-05 | 7.87E-07 | ENSG00000078114.18 | NEBL | 8417.6834 | 1.1673899 | 0.2363879 | 4.9384509 |
| 3.41E-05 | 8.00E-07 | ENSG00000067064.11 | IDI1 | 28235.32 | -1.188851 | 0.2408813 | -4.935423 |
| 3.69E-05 | 8.71E-07 | ENSG00000166886.13 | NAB2 | 2190.0355 | -1.122727 | 0.2282546 | -4.918746 |
| 3.73E-05 | 8.83E-07 | ENSG00000073464.12 | CLCN4 | 13437.907 | 1.0361682 | 0.210774 | 4.9160152 |
| 4.02E-05 | 9.55E-07 | ENSG00000280061.1 | AC011504.1 | 172.87726 | 2.0313802 | 0.4145163 | 4.9006039 |
| 4.03E-05 | 9.59E-07 | ENSG00000234710.2 | AC060834.2 | 27.887924 | -8.322143 | 1.6984457 | -4.899859 |
| 4.04E-05 | 9.64E-07 | ENSG00000160233.8 | LRRC3 | 2179.1167 | -1.047711 | 0.2138711 | -4.898798 |
| 4.08E-05 | 9.74E-07 | ENSG00000198342.10 | ZNF442 | 586.20832 | -1.365849 | 0.2789247 | -4.896835 |
| 4.11E-05 | 9.83E-07 | ENSG00000253308.2 | AC004080.1 | 29.445756 | 8.2663735 | 1.6887156 | 4.8950655 |
| 4.35E-05 | 1.04E-06 | ENSG00000234444.10 | ZNF736 | 1583.0847 | -1.255403 | 0.2570864 | -4.883192 |
| 4.37E-05 | 1.05E-06 | ENSG00000138180.16 | CEP55 | 37.582284 | -8.75218 | 1.7929112 | -4.881547 |
| 4.38E-05 | 1.06E-06 | ENSG00000102230.14 | PCYT1B | 4702.5771 | -1.154502 | 0.2365392 | -4.880806 |
| 4.47E-05 | 1.08E-06 | ENSG00000174276.7 | ZNHIT2 | 939.29825 | -1.474726 | 0.3024528 | -4.875889 |
| 4.52E-05 | 1.10E-06 | ENSG00000176463.14 | SLCO3A1 | 6953.3148 | 1.2818896 | 0.2630287 | 4.8735728 |
| 4.53E-05 | 1.10E-06 | ENSG00000251209.9 | LINC00923 | 451.38245 | 1.936102 | 0.3973243 | 4.8728508 |
| 4.55E-05 | 1.11E-06 | ENSG00000104549.12 | SQLE | 52229.096 | -1.057703 | 0.2171285 | -4.871325 |
| 4.60E-05 | 1.12E-06 | ENSG00000152578.13 | GRIA4 | 15790.867 | 1.174805 | 0.2413034 | 4.8685795 |
| 4.74E-05 | 1.16E-06 | ENSG00000165194.15 | PCDH19 | 1721.2315 | 1.7716211 | 0.3643791 | 4.8620275 |
| 4.76E-05 | 1.17E-06 | ENSG00000137831.15 | UACA | 138.36561 | -2.633576 | 0.5417791 | -4.860978 |
| 4.88E-05 | 1.20E-06 | ENSG00000135617.4 | PRADC1 | 605.28483 | -1.38589 | 0.2854185 | -4.855642 |
| 5.12E-05 | 1.26E-06 | ENSG00000188681.11 | TEKT4P2 | 28.777662 | 8.2272572 | 1.6979238 | 4.8454807 |
| 5.17E-05 | 1.28E-06 | ENSG00000112539.15 | C6orf118 | 153.14237 | -2.071195 | 0.4276477 | -4.843228 |
| 5.22E-05 | 1.29E-06 | ENSG00000119638.13 | NEK9 | 3471.9408 | 1.1083678 | 0.2289598 | 4.8408827 |
| 5.27E-05 | 1.31E-06 | ENSG00000214336.5 | FOXI3 | 143.33858 | 1.938597 | 0.4006455 | 4.8386844 |
| 5.27E-05 | 1.31E-06 | ENSG00000143507.18 | DUSP10 | 903.06273 | 1.2845914 | 0.2655033 | 4.8383261 |
| 5.29E-05 | 1.32E-06 | ENSG00000198879.12 | SFMBT2 | 4693.4158 | -1.298086 | 0.2683431 | -4.83741 |
| 5.33E-05 | 1.33E-06 | ENSG00000130032.17 | PRRG3 | 1138.66 | 1.1055826 | 0.2286269 | 4.8357495 |
| 5.37E-05 | 1.34E-06 | ENSG00000218336.9 | TENM3 | 13123.197 | 1.0888815 | 0.2252735 | 4.8335986 |
| 5.40E-05 | 1.35E-06 | ENSG00000279672.1 | AP006621.5 | 235.1387 | -2.191076 | 0.4534351 | -4.83217 |
| 5.46E-05 | 1.37E-06 | ENSG00000228775.8 | WEE2-AS1 | 69.676131 | -2.699168 | 0.5589141 | -4.829307 |
| 5.46E-05 | 1.37E-06 | ENSG00000235026.6 | DPP10-AS1 | 427.80164 | 1.7514975 | 0.3626974 | 4.8290879 |
| 5.94E-05 | 1.50E-06 | ENSG00000131089.17 | ARHGEF9 | 21945.442 | -1.040556 | 0.2162658 | -4.811468 |
| 6.02E-05 | 1.52E-06 | ENSG00000146006.8 | LRRTM2 | 4380.0693 | 1.29265 | 0.2688219 | 4.808574 |
| 6.15E-05 | 1.56E-06 | ENSG00000251573.2 | LINC02106 | 126.50179 | -2.040646 | 0.4248236 | -4.803514 |
| 6.32E-05 | 1.61E-06 | ENSG00000112238.12 | PRDM13 | 27.046535 | 8.1431975 | 1.6974246 | 4.797384 |
| 6.32E-05 | 1.61E-06 | ENSG00000164236.12 | ANKRD33B | 46.904475 | -4.002701 | 0.8343683 | -4.797284 |
| 6.41E-05 | 1.63E-06 | ENSG00000280234.1 | AC124303.2 | 458.68178 | 1.7638867 | 0.3679056 | 4.7944005 |
| 6.76E-05 | 1.73E-06 | ENSG00000134594.5 | RAB33A | 3487.9571 | -1.059654 | 0.2215384 | -4.783161 |
| 6.76E-05 | 1.73E-06 | ENSG00000160326.14 | SLC2A6 | 2102.1523 | -1.494421 | 0.3124446 | -4.782994 |
| 6.81E-05 | 1.74E-06 | ENSG00000165118.15 | C9orf64 | 209.31064 | 7.277715 | 1.5221683 | 4.7811499 |
| 6.92E-05 | 1.77E-06 | ENSG00000250303.4 | LINC02762 | 1528.1862 | -1.10749 | 0.2318045 | -4.77769 |
| 7.35E-05 | 1.89E-06 | ENSG00000070669.17 | ASNS | 17003.713 | -1.199513 | 0.2517641 | -4.764433 |
| 7.36E-05 | 1.90E-06 | ENSG00000261594.4 | TPBGL | 712.06034 | 1.5433219 | 0.3239661 | 4.7638383 |
| 7.47E-05 | 1.93E-06 | ENSG00000276043.5 | UHRF1 | 119.03763 | -3.494524 | 0.7340608 | -4.760538 |
| 7.47E-05 | 1.93E-06 | ENSG00000168938.6 | PPIC | 176.60839 | 1.8904185 | 0.3971272 | 4.7602346 |
| 7.49E-05 | 1.94E-06 | ENSG00000188917.15 | TRMT2B | 556.52227 | -1.60187 | 0.3365654 | -4.759462 |
| 7.52E-05 | 1.95E-06 | ENSG00000152192.8 | POU4F1 | 780.28333 | 2.0578669 | 0.4324793 | 4.7583012 |
| 7.60E-05 | 1.98E-06 | ENSG00000261292.2 | AC110491.1 | 173.84167 | 2.4195636 | 0.5087619 | 4.7557881 |
| 7.91E-05 | 2.06E-06 | ENSG00000121064.13 | SCPEP1 | 178.22477 | -2.029191 | 0.4274322 | -4.7474 |
| 8.14E-05 | 2.12E-06 | ENSG00000162998.5 | FRZB | 2321.0266 | 1.7979853 | 0.379215 | 4.7413343 |

| | | | | | | | |
|---|---|---|---|---|---|---|---|
| 8.16E-05 | 2.13E-06 | ENSG00000145536.15 | ADAMTS16 | 300.84796 | 1.5481867 | 0.3265778 | 4.7406373 |
| 8.30E-05 | 2.17E-06 | ENSG00000112319.19 | EYA4 | 54.82522 | 3.411846 | 0.7202861 | 4.7367926 |
| 8.39E-05 | 2.20E-06 | ENSG00000248801.7 | C8orf34-AS1 | 363.13942 | 2.101694 | 0.443941 | 4.7341745 |
| 8.41E-05 | 2.21E-06 | ENSG00000114450.10 | GNB4 | 2204.1303 | -1.323944 | 0.2797018 | -4.733412 |
| 8.83E-05 | 2.32E-06 | ENSG00000175985.10 | PLEKHD1 | 457.70234 | -1.511308 | 0.3199683 | -4.723305 |
| 9.07E-05 | 2.39E-06 | ENSG00000229407.5 | AL359853.1 | 450.3477 | 1.6897615 | 0.3581901 | 4.7174994 |
| 9.07E-05 | 2.39E-06 | ENSG00000117154.12 | IGSF21 | 2314.4586 | 1.4209745 | 0.3012389 | 4.7171018 |
| 9.22E-05 | 2.43E-06 | ENSG00000182752.10 | PAPPA | 1177.207 | -1.956235 | 0.415022 | -4.713571 |
| 9.35E-05 | 2.47E-06 | ENSG00000255408.4 | PCDHA3 | 1048.2456 | 1.0752577 | 0.2282739 | 4.7103849 |
| 9.44E-05 | 2.50E-06 | ENSG00000143786.8 | CNIH3 | 348.9623 | 1.3975222 | 0.2968407 | 4.7079879 |
| 9.58E-05 | 2.54E-06 | ENSG00000173626.10 | TRAPPC3L | 58.75849 | 2.9459451 | 0.6261673 | 4.704725 |
| 9.61E-05 | 2.55E-06 | ENSG00000185022.12 | MAFF | 1705.5481 | -1.298505 | 0.2760576 | -4.703748 |
| 9.68E-05 | 2.57E-06 | ENSG00000165175.15 | MID1IP1 | 2569.0143 | -1.009325 | 0.2146533 | -4.702119 |
| 9.69E-05 | 2.58E-06 | ENSG00000154079.6 | SDHAF4 | 1178.1382 | 1.1212984 | 0.2385007 | 4.7014476 |
| 9.85E-05 | 2.63E-06 | ENSG00000229236.3 | TTTY10 | 32.42893 | -4.489757 | 0.9557073 | -4.697837 |
| 9.90E-05 | 2.65E-06 | ENSG00000272916.5 | AC022400.7 | 1196.4869 | 1.0885611 | 0.2317759 | 4.6966107 |
| 9.99E-05 | 2.68E-06 | ENSG00000284762.1 | AC022414.1 | 81.573515 | 22.462492 | 4.7853762 | 4.6939867 |
| 0.0001006 | 2.70E-06 | ENSG00000204386.11 | NEU1 | 900.32688 | -1.123463 | 0.2394214 | -4.692409 |
| 0.0001016 | 2.74E-06 | ENSG00000148484.18 | RSU1 | 3082.9608 | -1.097138 | 0.2339453 | -4.689721 |
| 0.0001025 | 2.76E-06 | ENSG00000286058.1 | AC018865.2 | 390.38322 | -1.328048 | 0.2833131 | -4.687563 |
| 0.0001038 | 2.81E-06 | ENSG00000196544.8 | BORCS6 | 1253.9283 | -1.428582 | 0.3049652 | -4.684411 |
| 0.0001053 | 2.85E-06 | ENSG00000205213.14 | LGR4 | 4856.741 | 1.0528835 | 0.2249173 | 4.681204 |
| 0.0001055 | 2.86E-06 | ENSG00000101255.11 | TRIB3 | 1575.2534 | -1.423124 | 0.3040586 | -4.680428 |
| 0.0001057 | 2.87E-06 | ENSG00000250007.7 | AC087457.1 | 64.03733 | 3.3029372 | 0.7057935 | 4.6797504 |
| 0.0001057 | 2.88E-06 | ENSG00000198756.12 | COLGALT2 | 2875.1712 | 1.2835646 | 0.2742963 | 4.679482 |
| 0.0001096 | 2.99E-06 | ENSG00000080503.24 | SMARCA2 | 3257.2919 | -1.1652 | 0.2494357 | -4.671344 |
| 0.0001124 | 3.07E-06 | ENSG00000233016.7 | SNHG7 | 1313.953 | -1.180411 | 0.2529926 | -4.665794 |
| 0.0001136 | 3.11E-06 | ENSG00000204520.14 | MICA | 25.451621 | -8.189892 | 1.7562154 | -4.663376 |
| 0.0001218 | 3.36E-06 | ENSG00000058668.14 | ATP2B4 | 7118.1791 | -1.444303 | 0.3107514 | -4.647778 |
| 0.0001246 | 3.44E-06 | ENSG00000086205.18 | FOLH1 | 55.081884 | 3.4050057 | 0.7333795 | 4.6428973 |
| 0.0001248 | 3.45E-06 | ENSG00000164853.9 | UNCX | 24.614177 | 8.0022038 | 1.7237941 | 4.6422038 |
| 0.000127 | 3.52E-06 | ENSG00000163820.15 | FYCO1 | 176.18012 | -1.85032 | 0.3989736 | -4.6377 |
| 0.000127 | 3.52E-06 | ENSG00000088280.19 | ASAP3 | 2535.2633 | -1.100038 | 0.2371992 | -4.637612 |
| 0.0001285 | 3.57E-06 | ENSG00000261606.5 | AC091230.1 | 23.688834 | 7.949994 | 1.7152469 | 4.6348978 |
| 0.0001309 | 3.65E-06 | ENSG00000225988.1 | LAMP5-AS1 | 24.006313 | 7.9708873 | 1.7213812 | 4.6305183 |
| 0.0001372 | 3.85E-06 | ENSG00000140545.15 | MFGE8 | 203.9682 | -2.875764 | 0.6225208 | -4.619547 |
| 0.00014 | 3.94E-06 | ENSG00000116830.12 | TTF2 | 399.82723 | -1.412218 | 0.3060496 | -4.614344 |
| 0.00014 | 3.94E-06 | ENSG00000188803.15 | SHISA6 | 5243.618 | 1.8118638 | 0.3926641 | 4.6142843 |
| 0.00014 | 3.95E-06 | ENSG00000278847.1 | AC006157.1 | 22.546429 | -8.01561 | 1.7371605 | -4.614202 |
| 0.0001426 | 4.03E-06 | ENSG00000164318.18 | EGFLAM | 72.322457 | 2.9902967 | 0.6486455 | 4.610063 |
| 0.0001437 | 4.06E-06 | ENSG00000123360.12 | PDE1B | 765.32824 | 1.6332354 | 0.3544231 | 4.6081517 |
| 0.0001474 | 4.17E-06 | ENSG00000062282.15 | DGAT2 | 344.68628 | 1.3457447 | 0.292392 | 4.6025358 |
| 0.0001475 | 4.18E-06 | ENSG00000160685.13 | ZBTB7B | 206.46526 | -1.975524 | 0.4292685 | -4.602071 |
| 0.0001477 | 4.19E-06 | ENSG00000167065.13 | DUSP18 | 1278.4863 | -1.10164 | 0.2394091 | -4.601494 |
| 0.0001518 | 4.32E-06 | ENSG00000125844.16 | RRBP1 | 3051.817 | -1.316311 | 0.286434 | -4.595514 |
| 0.0001544 | 4.40E-06 | ENSG00000277957.1 | SENP3-EIF4/ | 652.62191 | 1.6791084 | 0.3657093 | 4.5913744 |
| 0.0001582 | 4.52E-06 | ENSG00000186153.17 | WWOX | 1544.8413 | 1.0785669 | 0.2352002 | 4.5857401 |
| 0.0001614 | 4.62E-06 | ENSG00000223518.5 | CSNK1A1P1 | 414.59074 | 1.6698481 | 0.3644955 | 4.5812588 |
| 0.0001675 | 4.82E-06 | ENSG00000163646.11 | CLRN1 | 23.48487 | 7.9399643 | 1.7364168 | 4.5726143 |
| 0.0001695 | 4.88E-06 | ENSG00000057593.14 | F7 | 137.80916 | 2.0460093 | 0.4477215 | 4.5698263 |
| 0.000184 | 5.31E-06 | ENSG00000067842.17 | ATP2B3 | 1671.0108 | 1.1946068 | 0.2624171 | 4.5523211 |
| 0.0001901 | 5.49E-06 | ENSG00000248918.2 | AC008808.1 | 44.003832 | -3.37897 | 0.7434232 | -4.545149 |
| 0.0001971 | 5.71E-06 | ENSG00000129451.12 | KLK10 | 466.76817 | -1.347853 | 0.2970843 | -4.536937 |
| 0.0002009 | 5.83E-06 | ENSG00000262877.5 | AC110285.2 | 459.50295 | 1.5687733 | 0.3461133 | 4.5325421 |
| 0.0002015 | 5.85E-06 | ENSG00000185742.7 | C11orf87 | 2925.4529 | 1.5402024 | 0.3398756 | 4.5316656 |
| 0.0002153 | 6.28E-06 | ENSG00000141753.7 | IGFBP4 | 576.49334 | 1.4324926 | 0.3171473 | 4.5168055 |
| 0.0002167 | 6.35E-06 | ENSG00000117013.17 | KCNQ4 | 338.08316 | 1.5319136 | 0.3393316 | 4.5145037 |
| 0.0002223 | 6.52E-06 | ENSG00000274220.1 | AC009163.7 | 278.7157 | 1.4128819 | 0.3133609 | 4.5088006 |
| 0.0002259 | 6.63E-06 | ENSG00000231925.12 | TAPBP | 405.59372 | -1.275577 | 0.283138 | -4.505142 |

| | | | | | | | |
|---|---|---|---|---|---|---|---|
| 0.0002354 | 6.92E-06 | ENSG00000259431.6 | THTPA | 281.19056 | -1.878356 | 0.4177802 | -4.496039 |
| 0.0002391 | 7.04E-06 | ENSG00000115271.11 | GCA | 942.62506 | -1.152965 | 0.2566452 | -4.492446 |
| 0.0002411 | 7.12E-06 | ENSG00000108018.15 | SORCS1 | 19183.406 | 1.6096705 | 0.3584943 | 4.4900865 |
| 0.0002453 | 7.25E-06 | ENSG00000152580.8 | IGSF10 | 229.44 | -2.359877 | 0.5260455 | -4.48607 |
| 0.0002456 | 7.27E-06 | ENSG00000090889.12 | KIF4A | 91.070603 | -2.800448 | 0.6243299 | -4.485526 |
| 0.0002488 | 7.38E-06 | ENSG00000166435.15 | XRRA1 | 703.83149 | -1.120661 | 0.2500082 | -4.482499 |
| 0.0002502 | 7.44E-06 | ENSG00000169903.7 | TM4SF4 | 961.44687 | 4.1799072 | 0.9328662 | 4.4807147 |
| 0.0002588 | 7.72E-06 | ENSG00000164867.11 | NOS3 | 147.95288 | -3.144369 | 0.7030129 | -4.472705 |
| 0.0002588 | 7.74E-06 | ENSG00000099282.10 | TSPAN15 | 252.47269 | -1.459272 | 0.3262905 | -4.472311 |
| 0.0002669 | 7.99E-06 | ENSG00000111052.7 | LIN7A | 14285.126 | 1.0692647 | 0.2394512 | 4.4654806 |
| 0.0002778 | 8.34E-06 | ENSG00000186998.16 | EMID1 | 569.15957 | 1.5769158 | 0.353863 | 4.4562888 |
| 0.0002791 | 8.40E-06 | ENSG00000260244.1 | AC104083.1 | 635.47187 | 1.1402693 | 0.2559681 | 4.4547314 |
| 0.0002842 | 8.58E-06 | ENSG00000178031.17 | ADAMTSL1 | 4829.3157 | 1.6908205 | 0.3799422 | 4.4502043 |
| 0.0002842 | 8.59E-06 | ENSG00000140105.18 | WARS1 | 17116.761 | -1.16875 | 0.2626437 | -4.449947 |
| 0.0002876 | 8.71E-06 | ENSG00000159387.8 | IRX6 | 181.27536 | -2.834738 | 0.6374621 | -4.446912 |
| 0.0002876 | 8.71E-06 | ENSG00000214944.9 | ARHGEF28 | 217.39818 | 2.8265012 | 0.6356172 | 4.4468611 |
| 0.0002909 | 8.83E-06 | ENSG00000234996.4 | AC098934.2 | 299.35473 | -1.447207 | 0.3256578 | -4.443949 |
| 0.0002909 | 8.84E-06 | ENSG00000112812.16 | PRSS16 | 138.35222 | -1.844409 | 0.4150492 | -4.443831 |
| 0.0002986 | 9.08E-06 | ENSG00000061337.15 | LZTS1 | 6275.5668 | -1.384947 | 0.3120719 | -4.437909 |
| 0.000299 | 9.12E-06 | ENSG00000159314.11 | ARHGAP27 | 198.22179 | 1.9509361 | 0.4396937 | 4.437035 |
| 0.0003047 | 9.32E-06 | ENSG00000135116.9 | HRK | 2673.9598 | 1.0809269 | 0.2438679 | 4.4324274 |
| 0.0003051 | 9.34E-06 | ENSG00000279565.1 | AL121835.2 | 578.39212 | 1.235273 | 0.2787263 | 4.43185 |
| 0.0003084 | 9.46E-06 | ENSG00000116991.10 | SIPA1L2 | 9481.0837 | 1.1764463 | 0.2656083 | 4.4292535 |
| 0.0003122 | 9.59E-06 | ENSG00000165458.14 | INPPL1 | 2152.9173 | -1.278918 | 0.2889367 | -4.42629 |
| 0.0003159 | 9.71E-06 | ENSG00000151615.3 | POU4F2 | 21.759463 | -7.963711 | 1.8003187 | -4.423501 |
| 0.000327 | 1.01E-05 | ENSG00000134215.16 | VAV3 | 4092.5385 | 1.4422455 | 0.3266173 | 4.4157052 |
| 0.0003278 | 1.01E-05 | ENSG00000277586.3 | NEFL | 417923.09 | -1.159766 | 0.2627095 | -4.414631 |
| 0.0003287 | 1.02E-05 | ENSG00000214548.18 | MEG3 | 39.220781 | -4.876727 | 1.104895 | -4.413747 |
| 0.0003299 | 1.02E-05 | ENSG00000102699.6 | PARP4 | 1414.2384 | -1.129006 | 0.2558831 | -4.412195 |
| 0.0003299 | 1.02E-05 | ENSG00000188223.9 | AD000671.1 | 157.22444 | -1.869435 | 0.4237033 | -4.412132 |
| 0.0003395 | 1.06E-05 | ENSG00000121440.15 | PDZRN3 | 4885.732 | 1.7498713 | 0.397218 | 4.4053174 |
| 0.0003418 | 1.06E-05 | ENSG00000183963.18 | SMTN | 773.03687 | -1.382836 | 0.3140241 | -4.403598 |
| 0.0003425 | 1.07E-05 | ENSG00000275160.1 | AL354718.1 | 73.794268 | 2.3687459 | 0.5379998 | 4.4028747 |
| 0.0003462 | 1.08E-05 | ENSG00000138771.16 | SHROOM3 | 482.28685 | -2.036238 | 0.4627847 | -4.399969 |
| 0.0003527 | 1.10E-05 | ENSG00000258334.1 | AC125611.4 | 332.25708 | -2.151758 | 0.4895183 | -4.395664 |
| 0.0003602 | 1.13E-05 | ENSG00000285253.1 | AC090517.4 | 309.48124 | 1.3972322 | 0.3182405 | 4.3904918 |
| 0.0003672 | 1.15E-05 | ENSG00000181019.13 | NQO1 | 210.86419 | -1.848071 | 0.4213527 | -4.386044 |
| 0.0003756 | 1.18E-05 | ENSG00000177045.10 | SIX5 | 45.274272 | -3.037451 | 0.6933459 | -4.38086 |
| 0.0003832 | 1.21E-05 | ENSG00000213064.10 | SFT2D2 | 1768.5963 | -1.105036 | 0.2525112 | -4.376184 |
| 0.0003907 | 1.23E-05 | ENSG00000138794.10 | CASP6 | 258.54427 | -1.490719 | 0.3410172 | -4.371391 |
| 0.0003913 | 1.24E-05 | ENSG00000012817.15 | KDM5D | 1688.0883 | -9.404139 | 2.1516131 | -4.370739 |
| 0.0003913 | 1.24E-05 | ENSG00000196724.12 | ZNF418 | 985.53081 | -1.040754 | 0.2381313 | -4.370504 |
| 0.0004069 | 1.29E-05 | ENSG00000125531.7 | FNDC11 | 397.96058 | -1.662101 | 0.3810931 | -4.361404 |
| 0.0004177 | 1.33E-05 | ENSG00000250284.3 | AC109439.1 | 1496.0963 | -1.169322 | 0.2684765 | -4.355398 |
| 0.0004312 | 1.37E-05 | ENSG00000287174.1 | AC092596.2 | 127.99889 | 2.0799285 | 0.4783796 | 4.347862 |
| 0.0004324 | 1.38E-05 | ENSG00000079308.19 | TNS1 | 2590.4945 | -1.204282 | 0.2770703 | -4.346486 |
| 0.0004324 | 1.38E-05 | ENSG00000286009.2 | AC244213.1 | 18.869851 | -7.75982 | 1.7853357 | -4.34642 |
| 0.0004377 | 1.40E-05 | ENSG00000099958.15 | DERL3 | 63.117593 | -2.575978 | 0.59311 | -4.343171 |
| 0.000459 | 1.48E-05 | ENSG00000278376.1 | AP004609.5 | 207.38703 | 1.4926677 | 0.3445985 | 4.3316135 |
| 0.00047 | 1.52E-05 | ENSG00000273113.1 | AC133528.1 | 136.68439 | -1.895342 | 0.438169 | -4.325595 |
| 0.0004755 | 1.54E-05 | ENSG00000187135.7 | VSTM2B | 433.75748 | 1.5940429 | 0.3687591 | 4.322722 |
| 0.0004811 | 1.56E-05 | ENSG00000145808.10 | ADAMTS19 | 142.15209 | 2.4926075 | 0.5770101 | 4.3198683 |
| 0.0004828 | 1.57E-05 | ENSG00000118503.15 | TNFAIP3 | 585.78719 | 1.1522792 | 0.2668046 | 4.3188125 |
| 0.0004861 | 1.58E-05 | ENSG00000225697.13 | SLC26A6 | 1311.1018 | -1.046609 | 0.2424376 | -4.317025 |
| 0.0004915 | 1.60E-05 | ENSG00000159399.10 | HK2 | 971.09523 | 4.5669862 | 1.0585676 | 4.3143077 |
| 0.000507 | 1.65E-05 | ENSG00000064666.15 | CNN2 | 3454.3152 | -1.0847 | 0.2518361 | -4.307167 |
| 0.0005163 | 1.69E-05 | ENSG00000278558.5 | TMEM191B | 326.13462 | 1.8670197 | 0.4339009 | 4.3028711 |
| 0.0005225 | 1.71E-05 | ENSG00000269821.1 | KCNQ1OT1 | 3938.9 | -1.552831 | 0.361148 | -4.299709 |
| 0.0005351 | 1.75E-05 | ENSG00000184697.7 | CLDN6 | 1010.9809 | -1.640357 | 0.3820016 | -4.294111 |

| | | | | | | | |
|---|---|---|---|---|---|---|---|
| 0.0005383 | 1.77E-05 | ENSG00000089847.12 | ANKRD24 | 1400.5882 | -1.060681 | 0.2471146 | -4.292262 |
| 0.0005432 | 1.79E-05 | ENSG00000101489.20 | CELF4 | 12765.305 | 1.2296285 | 0.2866276 | 4.2899866 |
| 0.0005468 | 1.80E-05 | ENSG00000213949.10 | ITGA1 | 331.51539 | -1.41424 | 0.3297941 | -4.28825 |
| 0.0005479 | 1.81E-05 | ENSG00000152128.13 | TMEM163 | 271.89557 | 1.3810184 | 0.3221022 | 4.2875159 |
| 0.0005492 | 1.81E-05 | ENSG00000243364.8 | EFNA4 | 71.833075 | -2.363903 | 0.5514489 | -4.286713 |
| 0.0005495 | 1.82E-05 | ENSG00000255136.3 | AP001972.3 | 533.73839 | 1.1987201 | 0.2796622 | 4.286315 |
| 0.0005511 | 1.82E-05 | ENSG00000230658.2 | KLHL7-DT | 64.219354 | 2.4965358 | 0.5825705 | 4.28538 |
| 0.0005604 | 1.86E-05 | ENSG00000206140.11 | TMEM191C | 605.2458 | 1.3442152 | 0.313965 | 4.281418 |
| 0.0005611 | 1.86E-05 | ENSG00000137693.14 | YAP1 | 114.46912 | -3.281641 | 0.7665827 | -4.280869 |
| 0.0005637 | 1.87E-05 | ENSG00000166033.13 | HTRA1 | 528.14894 | 1.1532277 | 0.2694725 | 4.2795742 |
| 0.000577 | 1.92E-05 | ENSG00000083814.13 | ZNF671 | 1197.3338 | -1.00904 | 0.2360889 | -4.273982 |
| 0.000577 | 1.92E-05 | ENSG00000115414.20 | FN1 | 949.02412 | -1.534792 | 0.3591151 | -4.273815 |
| 0.0005797 | 1.94E-05 | ENSG00000136371.11 | MTHFS | 347.56724 | 1.3289248 | 0.3110652 | 4.2721742 |
| 0.0005867 | 1.96E-05 | ENSG00000116117.18 | PARD3B | 175.79172 | 2.1087021 | 0.4939562 | 4.2690059 |
| 0.0006102 | 2.05E-05 | ENSG00000187714.7 | SLC18A3 | 7282.3144 | -1.510468 | 0.3546177 | -4.259425 |
| 0.0006368 | 2.16E-05 | ENSG00000147246.10 | HTR2C | 73.588664 | 3.0450464 | 0.7167991 | 4.248117 |
| 0.0006368 | 2.16E-05 | ENSG00000164106.8 | SCRG1 | 375.34718 | -5.181132 | 1.2196593 | -4.248016 |
| 0.0006479 | 2.20E-05 | ENSG00000121905.10 | HPCA | 430.1269 | 1.4474547 | 0.3410922 | 4.2435878 |
| 0.000648 | 2.21E-05 | ENSG00000237515.9 | SHISA9 | 1126.2503 | 1.779906 | 0.4194901 | 4.243023 |
| 0.0006491 | 2.21E-05 | ENSG00000213967.11 | ZNF726 | 238.32636 | 1.4245844 | 0.3358146 | 4.2421752 |
| 0.0006491 | 2.21E-05 | ENSG00000231205.11 | ZNF826P | 1141.5174 | -1.250557 | 0.2947952 | -4.242119 |
| 0.0006503 | 2.22E-05 | ENSG00000074219.14 | TEAD2 | 99.075543 | -2.703686 | 0.6374481 | -4.241422 |
| 0.0006593 | 2.25E-05 | ENSG00000224223.1 | VSTM2A-OT | 561.3563 | 1.6274761 | 0.3840141 | 4.2380635 |
| 0.0006711 | 2.30E-05 | ENSG00000102468.10 | HTR2A | 217.3798 | -2.789413 | 0.6588808 | -4.233563 |
| 0.0006711 | 2.30E-05 | ENSG00000197977.4 | ELOVL2 | 1553.9384 | 1.1592113 | 0.2738149 | 4.2335586 |
| 0.0006758 | 2.32E-05 | ENSG00000092820.18 | EZR | 1857.7929 | -1.768584 | 0.4179866 | -4.231199 |
| 0.0006769 | 2.33E-05 | ENSG00000180801.14 | ARSJ | 21.408607 | -7.940054 | 1.8768309 | -4.230564 |
| 0.0007004 | 2.42E-05 | ENSG00000036565.15 | SLC18A1 | 89.097235 | 2.3373234 | 0.5536322 | 4.2217978 |
| 0.0007004 | 2.42E-05 | ENSG00000166923.12 | GREM1 | 64.470027 | 3.3332063 | 0.7895427 | 4.221692 |
| 0.0007088 | 2.46E-05 | ENSG00000107731.12 | UNC5B | 840.42666 | -1.020251 | 0.2418451 | -4.218615 |
| 0.0007222 | 2.51E-05 | ENSG00000180440.4 | SERTM1 | 110.31088 | -2.441815 | 0.5794384 | -4.214106 |
| 0.0007322 | 2.55E-05 | ENSG00000113319.13 | RASGRF2 | 1923.3163 | 1.2787424 | 0.3037054 | 4.2104697 |
| 0.0007371 | 2.57E-05 | ENSG00000158457.6 | TSPAN33 | 419.50873 | -1.438591 | 0.3418129 | -4.20871 |
| 0.0007385 | 2.58E-05 | ENSG00000175567.10 | UCP2 | 557.7537 | -1.116198 | 0.265255 | -4.20802 |
| 0.0007661 | 2.68E-05 | ENSG00000197822.11 | OCLN | 225.9307 | 1.6102967 | 0.3834771 | 4.1991987 |
| 0.000783 | 2.74E-05 | ENSG00000241956.10 | AC109466.1 | 104.55423 | 2.6438279 | 0.6303856 | 4.1939852 |
| 0.0008025 | 2.81E-05 | ENSG00000173706.14 | HEG1 | 628.88541 | -1.115963 | 0.2664572 | -4.18815 |
| 0.0008096 | 2.84E-05 | ENSG00000249803.6 | AC112178.1 | 670.79357 | -1.078672 | 0.2576932 | -4.185878 |
| 0.0008116 | 2.85E-05 | ENSG00000266456.1 | AP001178.3 | 76.640768 | 2.3239196 | 0.5553191 | 4.1848365 |
| 0.0008232 | 2.90E-05 | ENSG00000145335.17 | SNCA | 20409.052 | -1.291597 | 0.3089182 | -4.181034 |
| 0.0008504 | 3.00E-05 | ENSG00000285769.2 | AP003100.2 | 19.654981 | 7.675578 | 1.8392842 | 4.1731332 |
| 0.0008852 | 3.14E-05 | ENSG00000171848.15 | RRM2 | 103.54709 | -3.619173 | 0.8693268 | -4.163191 |
| 0.0009136 | 3.24E-05 | ENSG00000123454.12 | DBH | 231.8761 | -1.457308 | 0.3506757 | -4.155714 |
| 0.0009156 | 3.26E-05 | ENSG00000285219.2 | HULC | 495.38305 | 1.1256295 | 0.2709203 | 4.154836 |
| 0.0009156 | 3.26E-05 | ENSG00000231453.1 | LINC01305 | 17.890717 | 7.5415501 | 1.8152956 | 4.1544475 |
| 0.0009279 | 3.32E-05 | ENSG00000167785.9 | ZNF558 | 1653.7574 | 8.9895942 | 2.1659853 | 4.1503486 |
| 0.0009292 | 3.33E-05 | ENSG00000077522.13 | ACTN2 | 325.67114 | 1.8839096 | 0.4539784 | 4.1497781 |
| 0.00094 | 3.38E-05 | ENSG00000247157.7 | LINC01252 | 80.885988 | 2.1229175 | 0.5120037 | 4.1462931 |
| 0.00094 | 3.38E-05 | ENSG00000112414.15 | ADGRG6 | 59.631771 | -3.000634 | 0.7237279 | -4.146081 |
| 0.0009444 | 3.41E-05 | ENSG00000160691.19 | SHC1 | 2012.2136 | -1.094288 | 0.2640339 | -4.144499 |
| 0.0009595 | 3.47E-05 | ENSG00000182809.11 | CRIP2 | 3618.2509 | -1.335023 | 0.3224417 | -4.140354 |
| 0.0009656 | 3.49E-05 | ENSG00000244560.7 | AC004890.2 | 448.13933 | 1.4118506 | 0.34114 | 4.138625 |
| 0.0009697 | 3.52E-05 | ENSG00000146242.9 | TPBG | 212.71028 | -2.180798 | 0.5271148 | -4.137235 |
| 0.0009716 | 3.53E-05 | ENSG00000236064.2 | AL109946.1 | 17.086947 | 7.4781384 | 1.8078664 | 4.1364441 |
| 0.0009838 | 3.58E-05 | ENSG00000178038.17 | ALS2CL | 136.07844 | 1.7535752 | 0.4242804 | 4.1330576 |
| 0.0009846 | 3.59E-05 | ENSG00000269473.1 | AC012313.8 | 186.1835 | -1.812047 | 0.4384747 | -4.132615 |
| 0.0009875 | 3.61E-05 | ENSG00000173894.11 | CBX2 | 1489.6761 | -1.386425 | 0.335601 | -4.13117 |
| 0.0010143 | 3.72E-05 | ENSG00000213578.6 | CPLX3 | 17.851536 | 7.537794 | 1.8275601 | 4.1245122 |
| 0.0010145 | 3.72E-05 | ENSG00000009950.16 | MLXIPL | 120.43684 | -2.031978 | 0.4927206 | -4.123996 |

| | | | | | | | |
|---|---|---|---|---|---|---|---|
| 0.0010145 | 3.72E-05 | ENSG00000204603.7 | LINC01257 | 30.165541 | 4.8360623 | 1.1726749 | 4.1239584 |
| 0.0010172 | 3.74E-05 | ENSG00000153317.15 | ASAP1 | 16123.565 | -1.085564 | 0.2633043 | -4.122847 |
| 0.0010251 | 3.78E-05 | ENSG00000258461.5 | AC012651.1 | 44.514839 | 4.3069011 | 1.0451621 | 4.1207973 |
| 0.0010522 | 3.88E-05 | ENSG00000153404.14 | PLEKHG4B | 504.63746 | -1.104035 | 0.2683419 | -4.114286 |
| 0.0010643 | 3.93E-05 | ENSG00000189410.12 | SH2D5 | 565.58356 | -1.077041 | 0.2619661 | -4.111377 |
| 0.0010696 | 3.96E-05 | ENSG00000282413.1 | AL133461.1 | 151.70113 | -1.622982 | 0.3949164 | -4.109684 |
| 0.0010696 | 3.97E-05 | ENSG00000153234.14 | NR4A2 | 801.46343 | -1.85173 | 0.4505985 | -4.109489 |
| 0.0010777 | 4.00E-05 | ENSG00000125851.10 | PCSK2 | 6191.1997 | 1.5668364 | 0.3814829 | 4.1072261 |
| 0.0011043 | 4.12E-05 | ENSG00000128573.26 | FOXP2 | 220.55991 | 2.2664359 | 0.5527087 | 4.1005974 |
| 0.0011065 | 4.14E-05 | ENSG00000079101.16 | CLUL1 | 378.01444 | 1.6657453 | 0.4063266 | 4.0995234 |
| 0.0011065 | 4.15E-05 | ENSG00000180257.13 | ZNF816 | 593.70805 | -1.076933 | 0.2627184 | -4.09919 |
| 0.0011065 | 4.15E-05 | ENSG00000148180.19 | GSN | 620.71757 | -1.740945 | 0.4247162 | -4.099078 |
| 0.0011065 | 4.15E-05 | ENSG00000286088.1 | AC073585.1 | 16.687077 | 7.4436921 | 1.8160347 | 4.0988712 |
| 0.0011289 | 4.24E-05 | ENSG00000184368.16 | MAP7D2 | 5175.2843 | 1.1437787 | 0.2793973 | 4.0937355 |
| 0.0011354 | 4.27E-05 | ENSG00000128610.12 | FEZF1 | 16.93933 | -7.602511 | 1.8578259 | -4.092154 |
| 0.0012056 | 4.56E-05 | ENSG00000117598.13 | PLPPR5 | 2227.1693 | 1.307315 | 0.320655 | 4.0770145 |
| 0.0012056 | 4.56E-05 | ENSG00000272078.1 | AL139423.1 | 202.09014 | 1.7721638 | 0.4346759 | 4.076977 |
| 0.0012226 | 4.64E-05 | ENSG00000237732.9 | CT75 | 270.3842 | 1.4176129 | 0.3480448 | 4.0730758 |
| 0.0012577 | 4.79E-05 | ENSG00000169071.15 | ROR2 | 128.57698 | 2.0461882 | 0.5032905 | 4.0656202 |
| 0.001262 | 4.81E-05 | ENSG00000197601.13 | FAR1 | 11179.746 | -1.004518 | 0.2471396 | -4.064577 |
| 0.0012764 | 4.88E-05 | ENSG00000282870.1 | FRG1DP | 16.40475 | 7.4212165 | 1.8272431 | 4.0614282 |
| 0.0013023 | 5.00E-05 | ENSG00000151136.15 | BTBD11 | 1377.8352 | 1.1523061 | 0.2841168 | 4.0557475 |
| 0.0013283 | 5.11E-05 | ENSG00000264515.6 | AC011474.1 | 18.574928 | 7.5937049 | 1.8748021 | 4.0504036 |
| 0.0013283 | 5.11E-05 | ENSG00000183780.13 | SLC35F3 | 850.1602 | 1.1360793 | 0.2804858 | 4.0503984 |
| 0.0013421 | 5.17E-05 | ENSG00000138606.19 | SHF | 5910.0945 | -1.035633 | 0.2558558 | -4.047723 |
| 0.0013571 | 5.24E-05 | ENSG00000239513.6 | LINC01210 | 50.193069 | 2.827268 | 0.6990187 | 4.0446246 |
| 0.0013589 | 5.25E-05 | ENSG00000184661.14 | CDCA2 | 71.475341 | -2.838938 | 0.7019999 | -4.044072 |
| 0.0013897 | 5.38E-05 | ENSG00000269994.3 | AL513318.2 | 510.03788 | 1.2680877 | 0.3140127 | 4.0383321 |
| 0.0013943 | 5.41E-05 | ENSG00000197852.12 | INKA2 | 1020.2926 | -1.113902 | 0.275919 | -4.037063 |
| 0.0014303 | 5.57E-05 | ENSG00000161638.11 | ITGA5 | 37.380497 | -5.568683 | 1.3816938 | -4.030331 |
| 0.0014321 | 5.58E-05 | ENSG00000285708.1 | AC097634.4 | 65.085455 | -3.79914 | 0.9427605 | -4.029804 |
| 0.0014406 | 5.62E-05 | ENSG00000069869.16 | NEDD4 | 586.41924 | -1.242463 | 0.3084438 | -4.028166 |
| 0.0014515 | 5.68E-05 | ENSG00000173208.4 | ABCD2 | 168.43538 | -1.632635 | 0.4055324 | -4.025905 |
| 0.0014614 | 5.72E-05 | ENSG00000186088.16 | GSAP | 434.29203 | 1.4065753 | 0.3495412 | 4.0240616 |
| 0.001502 | 5.90E-05 | ENSG00000240204.3 | SMKR1 | 265.89608 | 1.2851532 | 0.3199383 | 4.0168785 |
| 0.0015241 | 6.00E-05 | ENSG00000112245.12 | PTP4A1 | 3936.963 | 1.2868377 | 0.3206717 | 4.012944 |
| 0.0015281 | 6.02E-05 | ENSG00000267270.6 | PARD6G-AS1 | 152.82342 | -1.702587 | 0.4243652 | -4.012079 |
| 0.0015683 | 6.19E-05 | ENSG00000196604.13 | POTEF | 144.22865 | -1.621873 | 0.404915 | -4.005465 |
| 0.0015765 | 6.23E-05 | ENSG00000183196.10 | CHST6 | 108.43625 | -2.07881 | 0.5191859 | -4.00398 |
| 0.001579 | 6.25E-05 | ENSG00000285816.1 | AP000944.5 | 763.96483 | -1.035587 | 0.2586953 | -4.003116 |
| 0.0015826 | 6.27E-05 | ENSG00000197971.16 | MBP | 1847.8849 | 1.1763953 | 0.2939268 | 4.0023413 |
| 0.0015963 | 6.34E-05 | ENSG00000196277.16 | GRM7 | 2809.8027 | 1.1181913 | 0.2795602 | 3.999823 |
| 0.0016091 | 6.40E-05 | ENSG00000260641.1 | AC114811.2 | 1115.9575 | -1.141064 | 0.2854308 | -3.99769 |
| 0.0016273 | 6.48E-05 | ENSG00000162407.9 | PLPP3 | 305.58068 | 1.3245026 | 0.3315583 | 3.9947806 |
| 0.0016297 | 6.49E-05 | ENSG00000012660.14 | ELOVL5 | 16006.34 | -1.042562 | 0.2610196 | -3.994192 |
| 0.0016382 | 6.53E-05 | ENSG00000172824.16 | CES4A | 875.59922 | 1.387009 | 0.3473848 | 3.9927162 |
| 0.0016489 | 6.59E-05 | ENSG00000228203.7 | RNF144A-AS | 584.69818 | -1.100012 | 0.2756615 | -3.990444 |
| 0.0016598 | 6.65E-05 | ENSG00000162997.15 | PRORSD1P | 200.36724 | 1.3865705 | 0.3476414 | 3.9885077 |
| 0.0016899 | 6.78E-05 | ENSG00000189067.12 | LITAF | 622.76841 | -1.224855 | 0.3074512 | -3.9839 |
| 0.0017464 | 7.03E-05 | ENSG00000150361.12 | KLHL1 | 1909.9926 | 1.7379472 | 0.4371882 | 3.9752834 |
| 0.0017464 | 7.03E-05 | ENSG00000213793.5 | ZNF888 | 345.91933 | -1.210621 | 0.3045493 | -3.975123 |
| 0.0017554 | 7.08E-05 | ENSG00000128683.14 | GAD1 | 2295.1681 | 4.8728064 | 1.2263491 | 3.9734252 |
| 0.0017874 | 7.23E-05 | ENSG00000163071.11 | SPATA18 | 52.698982 | -3.040013 | 0.76605 | -3.968427 |
| 0.0017874 | 7.24E-05 | ENSG00000130222.11 | GADD45G | 352.08137 | -1.77516 | 0.4473247 | -3.968392 |
| 0.0017973 | 7.30E-05 | ENSG00000285367.1 | AC087564.1 | 109.58609 | 1.7817663 | 0.4492195 | 3.96636 |
| 0.0018022 | 7.32E-05 | ENSG00000283093.1 | CENPVL2 | 162.40642 | 1.4978304 | 0.3777177 | 3.9654754 |
| 0.0018138 | 7.39E-05 | ENSG00000163513.19 | TGFBR2 | 32.387802 | -3.803335 | 0.9595961 | -3.963475 |
| 0.0018172 | 7.41E-05 | ENSG00000146360.8 | GPR6 | 76.534246 | 2.9238705 | 0.7378306 | 3.9627936 |
| 0.0018472 | 7.55E-05 | ENSG00000189120.5 | SP6 | 343.88232 | 1.2346448 | 0.311923 | 3.9581716 |

| | | | | | | | |
|---|---|---|---|---|---|---|---|
| 0.00192 | 7.87E-05 | ENSG00000116691.11 | MIIP | 1498.4901 | -1.172122 | 0.2968748 | -3.948205 |
| 0.0019284 | 7.92E-05 | ENSG00000165948.11 | IFI27L1 | 499.67327 | -1.190344 | 0.3015871 | -3.946931 |
| 0.0020541 | 8.48E-05 | ENSG00000131969.15 | ABHD12B | 33.582576 | 3.429594 | 0.8725922 | 3.9303515 |
| 0.0021067 | 8.72E-05 | ENSG00000064961.19 | HMG20B | 1031.5089 | -1.454054 | 0.3705735 | -3.923794 |
| 0.0021082 | 8.73E-05 | ENSG00000188219.14 | POTEE | 212.71886 | -1.395817 | 0.3557679 | -3.923393 |
| 0.0021149 | 8.77E-05 | ENSG00000102349.18 | KLF8 | 1272.4918 | -1.120839 | 0.2857536 | -3.922398 |
| 0.0022009 | 9.17E-05 | ENSG00000140939.14 | NOL3 | 584.97301 | -1.11285 | 0.2844999 | -3.911601 |
| 0.0022129 | 9.23E-05 | ENSG00000128335.14 | APOL2 | 252.34247 | -1.696888 | 0.4339805 | -3.910056 |
| 0.0022161 | 9.25E-05 | ENSG00000227811.3 | INKA2-AS1 | 749.83568 | -1.169373 | 0.2991132 | -3.909466 |
| 0.0022276 | 9.31E-05 | ENSG00000280132.1 | AC026471.6 | 78.268636 | -2.207168 | 0.5647842 | -3.907985 |
| 0.0022457 | 9.40E-05 | ENSG00000168481.9 | LGI3 | 401.09053 | 1.1700239 | 0.2995791 | 3.9055586 |
| 0.0022492 | 9.42E-05 | ENSG00000163297.17 | ANTXR2 | 727.03282 | -1.224492 | 0.3135744 | -3.904949 |
| 0.0022551 | 9.46E-05 | ENSG00000223591.5 | CENPVL1 | 160.84612 | 1.4788833 | 0.3788043 | 3.9040828 |
| 0.0023417 | 9.86E-05 | ENSG00000008118.10 | CAMK1G | 377.65299 | -1.300152 | 0.333883 | -3.894034 |
| 0.0023417 | 9.87E-05 | ENSG00000064989.13 | CALCRL | 49.086939 | 2.57383 | 0.6609989 | 3.8938488 |
| 0.0023417 | 9.87E-05 | ENSG00000131650.14 | KREMEN2 | 165.91021 | -1.484833 | 0.3813337 | -3.893791 |
| 0.0023487 | 9.91E-05 | ENSG00000136250.12 | AOAH | 107.28252 | 1.7417185 | 0.4474174 | 3.8928269 |
| 0.0023643 | 9.99E-05 | ENSG00000168542.16 | COL3A1 | 253.9394 | -5.596911 | 1.4385149 | -3.890757 |
| 0.0024616 | 0.0001042 | ENSG00000254035.1 | AC104117.3 | 44.876127 | 2.768658 | 0.7134809 | 3.8804934 |
| 0.0024634 | 0.0001044 | ENSG00000196776.16 | CD47 | 23579.831 | 1.0892831 | 0.2807365 | 3.8800908 |
| 0.0024925 | 0.0001058 | ENSG00000175063.17 | UBE2C | 28.163919 | -5.390111 | 1.3902811 | -3.876994 |
| 0.0024997 | 0.0001062 | ENSG00000178878.12 | APOLD1 | 164.73473 | 1.5909267 | 0.4104492 | 3.8760621 |
| 0.0025235 | 0.0001074 | ENSG00000131002.12 | TXLNGY | 1038.5077 | -8.546652 | 2.2065635 | -3.873286 |
| 0.0025336 | 0.0001079 | ENSG00000227012.3 | LINC02527 | 14.446595 | 7.2351687 | 1.8685466 | 3.8720836 |
| 0.0025447 | 0.0001085 | ENSG00000173376.14 | NDNF | 590.89158 | 4.023212 | 1.0393795 | 3.8707825 |
| 0.0025492 | 0.000109 | ENSG00000257185.2 | LINC02293 | 128.04999 | 1.9903711 | 0.5143525 | 3.8696637 |
| 0.0025735 | 0.0001102 | ENSG00000252690.3 | AC105339.2 | 377.98357 | 1.0936585 | 0.2828265 | 3.8668876 |
| 0.0026223 | 0.0001125 | ENSG00000198417.7 | MT1F | 77.474789 | 2.8913998 | 0.7487115 | 3.8618345 |
| 0.0026973 | 0.000116 | ENSG00000244509.4 | APOBEC3C | 16.609786 | -7.57378 | 1.9649295 | -3.854479 |
| 0.0028206 | 0.0001221 | ENSG00000138653.10 | NDST4 | 293.19195 | 1.6119089 | 0.4195585 | 3.8419169 |
| 0.0028375 | 0.0001229 | ENSG00000099953.10 | MMP11 | 766.73201 | -1.16075 | 0.3022614 | -3.84022 |
| 0.0028817 | 0.0001251 | ENSG00000230982.1 | DSTNP1 | 298.27597 | -1.210743 | 0.3156293 | -3.835965 |
| 0.0029026 | 0.0001261 | ENSG00000214216.10 | IQCJ | 85.654505 | 2.0637451 | 0.5382808 | 3.8339564 |
| 0.0029286 | 0.0001273 | ENSG00000175785.13 | PRIMA1 | 352.73389 | 1.1572342 | 0.3020289 | 3.8315353 |
| 0.0029732 | 0.0001294 | ENSG00000172250.16 | SERHL | 131.41079 | 1.6788513 | 0.4386189 | 3.8275855 |
| 0.003024 | 0.0001318 | ENSG00000138722.10 | MMRN1 | 47.80532 | -2.607087 | 0.6819478 | -3.823001 |
| 0.0030629 | 0.0001338 | ENSG00000265174.1 | AP005202.1 | 50.012094 | -4.140169 | 1.083999 | -3.819348 |
| 0.0031341 | 0.0001373 | ENSG00000109929.10 | SC5D | 39401.726 | -1.062264 | 0.2785914 | -3.812982 |
| 0.003135 | 0.0001375 | ENSG00000157570.12 | TSPAN18 | 580.46492 | 1.0529746 | 0.2761768 | 3.8126835 |
| 0.0032507 | 0.0001429 | ENSG00000150594.7 | ADRA2A | 395.52631 | 1.680303 | 0.4418326 | 3.803031 |
| 0.0033581 | 0.0001481 | ENSG00000184497.13 | TMEM255B | 188.51887 | -1.424421 | 0.3754115 | -3.794293 |
| 0.0034092 | 0.0001507 | ENSG00000268603.1 | AC053503.4 | 172.81808 | 1.8830737 | 0.4968719 | 3.7898573 |
| 0.003434 | 0.000152 | ENSG00000080224.17 | EPHA6 | 418.51666 | 1.9548359 | 0.5160836 | 3.7878281 |
| 0.0034374 | 0.0001523 | ENSG00000270617.1 | URGCP-MRF | 266.13134 | 2.0589041 | 0.5436258 | 3.7873556 |
| 0.0034797 | 0.0001544 | ENSG00000124145.6 | SDC4 | 18.615714 | -7.738352 | 2.0450932 | -3.783863 |
| 0.0035418 | 0.0001575 | ENSG00000187984.13 | ANKRD19P | 140.9791 | 1.595104 | 0.4220969 | 3.7789998 |
| 0.0036017 | 0.0001604 | ENSG00000168016.14 | TRANK1 | 647.17842 | 1.3990062 | 0.3706594 | 3.7743713 |
| 0.0036039 | 0.0001607 | ENSG00000288597.1 | AC234782.4 | 126.82495 | 1.9416548 | 0.5144942 | 3.7739099 |
| 0.0036617 | 0.0001638 | ENSG00000122042.10 | UBL3 | 12048.907 | 1.0225142 | 0.2712875 | 3.7691164 |
| 0.0036628 | 0.000164 | ENSG00000215277.8 | RNF212B | 12.948788 | -7.216419 | 1.9147718 | -3.768814 |
| 0.0036765 | 0.0001648 | ENSG00000137834.15 | SMAD6 | 14.960449 | -7.422913 | 1.9701674 | -3.767656 |
| 0.0039146 | 0.0001763 | ENSG00000165509.13 | MAGEC3 | 12.826643 | -7.201951 | 1.9200975 | -3.750825 |
| 0.0039929 | 0.0001804 | ENSG00000125510.18 | OPRL1 | 365.21223 | -1.261499 | 0.3368533 | -3.744951 |
| 0.0040746 | 0.0001846 | ENSG00000101638.13 | ST8SIA5 | 471.06784 | 1.1866738 | 0.3173615 | 3.7391859 |
| 0.0041128 | 0.0001865 | ENSG00000151388.11 | ADAMTS12 | 136.05941 | -2.219569 | 0.5940046 | -3.736619 |
| 0.00421 | 0.0001916 | ENSG00000172799.5 | ZBTB8OSP2 | 25.720746 | 3.7287292 | 0.9997035 | 3.7298351 |
| 0.0043022 | 0.0001967 | ENSG00000123130.17 | ACOT9 | 1198.264 | -1.11833 | 0.300364 | -3.72325 |
| 0.0043169 | 0.000198 | ENSG00000284981.1 | UPK3BL2 | 81.835257 | 2.4080228 | 0.6470578 | 3.7214953 |
| 0.0043174 | 0.0001982 | ENSG00000037280.16 | FLT4 | 169.09669 | 1.4008544 | 0.376448 | 3.7212431 |

| | | | | | | | |
|---|---|---|---|---|---|---|---|
| 0.0043329 | 0.0001991 | ENSG00000189190.10 | ZNF600 | 151.47621 | -1.698047 | 0.45645 | -3.720117 |
| 0.0043534 | 0.0002004 | ENSG00000163046.15 | ANKRD30BL | 13.921123 | 7.1871685 | 1.9328241 | 3.7184804 |
| 0.0043943 | 0.0002025 | ENSG00000287712.1 | AC005865.2 | 13.086231 | 7.0921758 | 1.9086061 | 3.7158928 |
| 0.0044097 | 0.0002034 | ENSG00000260776.5 | AC104758.2 | 14.693224 | -7.399895 | 1.9920119 | -3.714784 |
| 0.0044452 | 0.0002052 | ENSG00000167992.13 | VWCE | 523.07302 | 1.0212519 | 0.275082 | 3.7125357 |
| 0.0044697 | 0.0002067 | ENSG00000271614.1 | ATP2B1-AS1 | 3164.6572 | -1.311617 | 0.3534689 | -3.710699 |
| 0.0045031 | 0.0002088 | ENSG00000074181.9 | NOTCH3 | 257.17008 | -2.138216 | 0.576626 | -3.708151 |
| 0.0045463 | 0.000211 | ENSG00000269891.2 | ARHGAP19-! | 115.05921 | 1.6082965 | 0.4340285 | 3.705509 |
| 0.0046269 | 0.0002151 | ENSG00000128591.15 | FLNC | 494.48324 | -1.664835 | 0.4498814 | -3.700609 |
| 0.0046734 | 0.0002176 | ENSG00000151338.18 | MIPOL1 | 229.78873 | -1.29574 | 0.3504245 | -3.69763 |
| 0.0046881 | 0.0002186 | ENSG00000067048.17 | DDX3Y | 3732.8543 | -9.312678 | 2.5193483 | -3.696463 |
| 0.0046881 | 0.0002187 | ENSG00000154240.17 | CEP112 | 211.11829 | -1.528983 | 0.4136425 | -3.696387 |
| 0.0047846 | 0.0002234 | ENSG00000169855.20 | ROBO1 | 26995.298 | 1.2048806 | 0.3264383 | 3.6909908 |
| 0.0048027 | 0.0002244 | ENSG00000237943.7 | PRKCQ-AS1 | 164.14638 | -1.896775 | 0.5140578 | -3.689809 |
| 0.0048257 | 0.0002257 | ENSG00000137812.20 | KNL1 | 223.78909 | -1.487418 | 0.4032719 | -3.688374 |
| 0.0048467 | 0.0002269 | ENSG00000011638.10 | TMEM159 | 1482.8822 | -1.088554 | 0.2952374 | -3.687048 |
| 0.0048995 | 0.0002297 | ENSG00000225470.8 | JPX | 7163.7501 | 1.2802575 | 0.347533 | 3.6838446 |
| 0.0049377 | 0.0002319 | ENSG00000285437.1 | POLR2J3 | 294.06763 | 1.1801414 | 0.3205666 | 3.6814231 |
| 0.0050215 | 0.0002365 | ENSG00000133640.20 | LRRIQ1 | 390.14668 | -1.13239 | 0.3080098 | -3.676475 |
| 0.0050387 | 0.0002375 | ENSG00000273540.4 | AGBL1 | 88.671617 | 2.0503478 | 0.5578602 | 3.6753791 |
| 0.0051229 | 0.0002423 | ENSG00000138650.9 | PCDH10 | 9355.9107 | -1.058584 | 0.2884213 | -3.670269 |
| 0.005134 | 0.000243 | ENSG00000180178.11 | FAR2P1 | 111.49564 | -1.787307 | 0.4870715 | -3.669495 |
| 0.0051748 | 0.0002454 | ENSG00000147036.12 | LANCL3 | 1326.1979 | -1.032404 | 0.2815346 | -3.66706 |
| 0.0051748 | 0.0002455 | ENSG00000170231.16 | FABP6 | 322.93123 | -1.134386 | 0.3093543 | -3.666947 |
| 0.0051748 | 0.0002456 | ENSG00000186815.12 | TPCN1 | 312.20719 | 1.140274 | 0.3109714 | 3.6668136 |
| 0.0052238 | 0.0002481 | ENSG00000175356.13 | SCUBE2 | 256.26896 | 1.2558794 | 0.3427445 | 3.6641854 |
| 0.0052468 | 0.0002494 | ENSG00000150394.14 | CDH8 | 11034.289 | 1.1749983 | 0.3207888 | 3.6628413 |
| 0.0053082 | 0.000253 | ENSG00000104267.10 | CA2 | 83.099293 | 2.1042351 | 0.5750519 | 3.659209 |
| 0.0053286 | 0.0002542 | ENSG00000167595.15 | PROSER3 | 921.41777 | -1.015601 | 0.2776379 | -3.658006 |
| 0.0053917 | 0.0002576 | ENSG00000088325.16 | TPX2 | 410.68808 | -1.515224 | 0.4146132 | -3.654549 |
| 0.0054515 | 0.0002614 | ENSG00000108439.11 | PNPO | 1119.9079 | 1.0974386 | 0.3005984 | 3.6508468 |
| 0.0054526 | 0.0002616 | ENSG00000287477.1 | AL356495.1 | 23.650075 | 4.4697469 | 1.2243935 | 3.6505805 |
| 0.0054771 | 0.0002635 | ENSG00000273899.5 | NOL12 | 582.99986 | 1.081469 | 0.2963921 | 3.6487777 |
| 0.0055009 | 0.0002649 | ENSG00000167900.12 | TK1 | 15.045701 | -7.431031 | 2.0373247 | -3.647446 |
| 0.0056339 | 0.0002715 | ENSG00000144152.13 | FBLN7 | 410.49059 | 1.0254971 | 0.2816461 | 3.6410842 |
| 0.0056776 | 0.0002743 | ENSG00000185567.7 | AHNAK2 | 5993.6344 | -1.169549 | 0.3214418 | -3.638446 |
| 0.0057045 | 0.0002758 | ENSG00000088756.12 | ARHGAP28 | 6643.3961 | -3.123179 | 0.858721 | -3.637012 |
| 0.0057425 | 0.0002779 | ENSG00000215030.5 | RPL13P12 | 1720.2362 | 1.0881519 | 0.2993471 | 3.6350836 |
| 0.0058063 | 0.0002819 | ENSG00000112984.12 | KIF20A | 44.09968 | -3.281177 | 0.9035641 | -3.631372 |
| 0.0058559 | 0.0002846 | ENSG00000284337.1 | AC013271.1 | 19.466788 | 5.2061588 | 1.4346141 | 3.6289611 |
| 0.0058753 | 0.0002857 | ENSG00000152661.9 | GJA1 | 162.60356 | -6.151067 | 1.6954938 | -3.627891 |
| 0.0059057 | 0.0002875 | ENSG00000100979.15 | PLTP | 554.84421 | -1.265707 | 0.349036 | -3.626295 |
| 0.0059057 | 0.0002877 | ENSG00000236682.2 | MAP3K2-DT | 214.43664 | -1.250992 | 0.3449939 | -3.626129 |
| 0.0059374 | 0.0002897 | ENSG00000153714.6 | LURAP1L | 398.45727 | -1.240922 | 0.3423881 | -3.624313 |
| 0.0059808 | 0.0002923 | ENSG00000223652.2 | AC106786.1 | 27.634694 | 3.2728968 | 0.9036153 | 3.6220024 |
| 0.0060201 | 0.0002945 | ENSG00000239445.6 | ST3GAL6-AS | 135.42155 | 1.5999844 | 0.4419735 | 3.6200916 |
| 0.0060299 | 0.0002955 | ENSG00000261168.1 | AL592424.1 | 404.04744 | -1.015669 | 0.2806303 | -3.619243 |
| 0.0061294 | 0.000301 | ENSG00000184898.7 | RBM43 | 77.313851 | 2.2111679 | 0.6117576 | 3.6144511 |
| 0.0061294 | 0.0003011 | ENSG00000244405.8 | ETV5 | 1677.7041 | -1.19719 | 0.3312311 | -3.614364 |
| 0.0061969 | 0.0003049 | ENSG00000154856.13 | APCDD1 | 277.80655 | 1.3822792 | 0.3827867 | 3.6110952 |
| 0.006342 | 0.0003131 | ENSG00000276644.5 | DACH1 | 5218.9215 | 1.3938178 | 0.3867168 | 3.6042337 |
| 0.0063679 | 0.0003146 | ENSG00000165124.18 | SVEP1 | 442.76414 | 1.1570085 | 0.321127 | 3.6029623 |
| 0.0064351 | 0.0003182 | ENSG00000146409.12 | SLC18B1 | 695.10184 | 1.0237065 | 0.2843614 | 3.6000197 |
| 0.0064736 | 0.0003206 | ENSG00000283294.1 | AP005212.4 | 222.5692 | 1.83309 | 0.5094688 | 3.5980422 |
| 0.0064758 | 0.000321 | ENSG00000180332.6 | KCTD4 | 49.926847 | 2.7555647 | 0.7659151 | 3.5977416 |
| 0.0066359 | 0.00033 | ENSG00000158296.14 | SLC13A3 | 261.49229 | 1.1586167 | 0.3226868 | 3.5905304 |
| 0.0066553 | 0.0003312 | ENSG00000262096.2 | PCDHB19P | 88.933925 | -1.816295 | 0.505994 | -3.589559 |
| 0.0067712 | 0.0003386 | ENSG00000223773.7_PAF | CD99P1 | 58.376912 | -2.14926 | 0.5997068 | -3.583851 |
| 0.0067712 | 0.0003386 | ENSG00000223773.7 | CD99P1 | 58.376912 | -2.14926 | 0.5997068 | -3.583851 |

| | | | | | | | |
|---|---|---|---|---|---|---|---|
| 0.0068018 | 0.0003407 | ENSG00000041982.16 | TNC | 393.13741 | -4.940447 | 1.3791736 | -3.58218 |
| 0.0068786 | 0.0003454 | ENSG00000095596.12 | CYP26A1 | 62.110781 | 2.4195729 | 0.6761124 | 3.5786547 |
| 0.0068957 | 0.0003473 | ENSG00000257379.1 | AC023509.1 | 440.64324 | 1.0083886 | 0.2818929 | 3.5772053 |
| 0.0069183 | 0.0003488 | ENSG00000204033.10 | LRIT2 | 166.66966 | -1.92658 | 0.5387446 | -3.576054 |
| 0.0069645 | 0.0003517 | ENSG00000101292.7 | PROKR2 | 59.311248 | 2.8612376 | 0.8005946 | 3.5738906 |
| 0.0069719 | 0.0003524 | ENSG00000230910.4 | AL391807.1 | 3430.0479 | 1.0900422 | 0.305043 | 3.5734047 |
| 0.007197 | 0.0003655 | ENSG00000266714.9 | MYO15B | 871.32337 | 1.2116926 | 0.3399984 | 3.5638187 |
| 0.0072197 | 0.0003669 | ENSG00000185633.10 | NDUFA4L2 | 370.9954 | 1.1160387 | 0.313249 | 3.5627839 |
| 0.0072303 | 0.000368 | ENSG00000236279.7 | CLEC2L | 437.54999 | 1.3125984 | 0.3684956 | 3.5620467 |
| 0.0072371 | 0.000369 | ENSG00000284070.1 | AP000356.2 | 145.5278 | 1.4763629 | 0.4145555 | 3.5613155 |
| 0.0072641 | 0.0003707 | ENSG00000257842.6 | LINC02588 | 346.38289 | 1.7339052 | 0.4870344 | 3.5601285 |
| 0.0072914 | 0.0003724 | ENSG00000118804.8 | STBD1 | 208.23638 | -2.228375 | 0.6261357 | -3.558934 |
| 0.0074157 | 0.0003802 | ENSG00000185436.12 | IFNLR1 | 250.85526 | -1.234179 | 0.3473188 | -3.553447 |
| 0.0075061 | 0.0003852 | ENSG00000164465.19 | DCBLD1 | 623.4523 | 1.3626061 | 0.3838272 | 3.5500514 |
| 0.0075359 | 0.000387 | ENSG00000267353.1 | AC020928.2 | 11.723852 | 6.9359421 | 1.9544467 | 3.5488009 |
| 0.0075931 | 0.0003905 | ENSG00000213085.10 | CFAP45 | 11.262558 | -7.014298 | 1.9778678 | -3.546394 |
| 0.0076045 | 0.0003914 | ENSG00000176769.9 | TCERG1L | 1141.6652 | -8.298634 | 2.3404187 | -3.54579 |
| 0.0077038 | 0.0003975 | ENSG00000109911.19 | ELP4 | 1871.9444 | 1.1122283 | 0.3140338 | 3.5417469 |
| 0.0077112 | 0.0003982 | ENSG00000173530.6 | TNFRSF10D | 77.849917 | -3.046761 | 0.8603543 | -3.541287 |
| 0.007853 | 0.0004071 | ENSG00000215271.9 | HOMEZ | 1863.5578 | -1.285667 | 0.363651 | -3.535442 |
| 0.0079206 | 0.0004112 | ENSG00000100433.15 | KCNK10 | 2017.2339 | -1.001615 | 0.2835217 | -3.532763 |
| 0.0079661 | 0.0004139 | ENSG00000285778.2 | AL591463.1 | 60.356034 | 2.6746727 | 0.7574741 | 3.5310416 |
| 0.0080306 | 0.0004176 | ENSG00000279114.1 | Z99129.3 | 168.85653 | -1.331489 | 0.3773314 | -3.5287 |
| 0.0081341 | 0.0004236 | ENSG00000240990.10 | HOXA11-AS | 11.802214 | 6.9476711 | 1.971028 | 3.5248972 |
| 0.0084574 | 0.0004424 | ENSG00000122592.8 | HOXA7 | 786.30389 | 1.4582324 | 0.4150484 | 3.5134032 |
| 0.0084598 | 0.000443 | ENSG00000114654.7 | EFCC1 | 10.969907 | -6.976518 | 1.9858938 | -3.513037 |
| 0.0084809 | 0.0004445 | ENSG00000104888.10 | SLC17A7 | 184.1525 | -1.645414 | 0.4684893 | -3.51217 |
| 0.0085517 | 0.0004494 | ENSG00000254768.6 | AC104009.1 | 138.55303 | 2.5881702 | 0.737526 | 3.5092594 |
| 0.0085517 | 0.0004496 | ENSG00000285525.1 | AC099670.3 | 11.975738 | -7.101945 | 2.0238418 | -3.50914 |
| 0.0085702 | 0.0004509 | ENSG00000160284.15 | SPATC1L | 266.36457 | 1.2106486 | 0.3450754 | 3.5083593 |
| 0.0086107 | 0.0004534 | ENSG00000168959.14 | GRM5 | 1153.7835 | 3.3284746 | 0.9491215 | 3.5069004 |
| 0.0086161 | 0.000454 | ENSG00000169946.14 | ZFPM2 | 356.36798 | -1.930672 | 0.5505938 | -3.506526 |
| 0.0087868 | 0.0004633 | ENSG00000272636.4 | DOC2B | 4819.5049 | -2.534803 | 0.7240022 | -3.501099 |
| 0.0088369 | 0.0004664 | ENSG00000227863.3 | AC002383.1 | 11.841395 | 6.9533656 | 1.9870467 | 3.4993468 |
| 0.0088369 | 0.0004667 | ENSG00000182759.4 | MAFA | 5528.9819 | -1.267093 | 0.3621124 | -3.499172 |
| 0.0088476 | 0.000468 | ENSG00000146555.19 | SDK1 | 3714.7302 | -1.007067 | 0.2878616 | -3.498442 |
| 0.0090497 | 0.0004801 | ENSG00000279656.1 | AL132780.4 | 134.59106 | 1.5270591 | 0.4373527 | 3.4915966 |
| 0.0091693 | 0.0004872 | ENSG00000179967.11 | PPP1R14BP3 | 320.71919 | -1.113474 | 0.3192594 | -3.487679 |
| 0.0092704 | 0.000493 | ENSG00000196834.12 | POTEI | 86.719809 | -1.698466 | 0.4874287 | -3.484542 |
| 0.0094515 | 0.0005034 | ENSG00000277268.2 | LHX1-DT | 291.38004 | 1.5037079 | 0.4322303 | 3.4789506 |
| 0.0095421 | 0.0005086 | ENSG00000112667.13 | DNPH1 | 314.55253 | 1.204206 | 0.3464156 | 3.4761888 |
| 0.0096142 | 0.0005128 | ENSG00000151834.15 | GABRA2 | 9280.1224 | 1.2273336 | 0.3532946 | 3.4739662 |
| 0.0097898 | 0.0005226 | ENSG00000014138.9 | POLA2 | 262.75963 | 1.1271634 | 0.3249337 | 3.468903 |
| 0.0098141 | 0.0005243 | ENSG00000178460.18 | MCMDC2 | 447.25664 | 1.1861035 | 0.3420105 | 3.4680328 |

| FoldChange |
|---|
| 97.062065 |
| 95.498756 |
| **47.246555** |
| 0.0052388 |
| 25.987684 |
| 0.0046708 |
| 0.0791807 |
| 0.0295356 |
| 0.0889962 |
| 21.303439 |
| 25.933578 |
| 32.538234 |
| 49.94702 |
| 7.6390488 |
| 49.376109 |
| 0.0857052 |
| 11.977227 |
| 0.0105451 |
| 30.743656 |
| 6.2742304 |
| 21.853511 |
| 19.203259 |
| 0.0133471 |
| 0.178664 |
| 6.2418521 |
| 0.127951 |
| 22.4076 |
| 0.1001865 |
| 0.2049443 |
| 0.1353542 |
| 0.1506485 |
| 0.1499793 |
| 15.16589 |
| 16.113164 |
| 0.0701046 |
| 0.0571195 |
| 0.0252175 |
| 0.1481322 |
| 5.4760184 |
| 0.2047121 |
| 0.1656751 |
| 15.296782 |
| 0.1669784 |
| 20.473317 |
| 53.947455 |
| 9.550949 |
| 0.1865833 |
| 0.0556525 |
| 0.0041633 |
| 8.3768018 |
| 0.1228336 |
| 0.1855311 |
| 4.3480511 |
| 4.8702374 |
| 8.4063025 |
| 5.8735847 |
| 14.396254 |
| 0.1513377 |

| |
|---|
| 8.57E-05 |
| 3.5700852 |
| 0.2354913 |
| 12.484255 |
| 24.418523 |
| 6.1515927 |
| 3.5099667 |
| 0.2378562 |
| 23.10513 |
| 0.0001403 |
| 0.2221926 |
| 0.2496673 |
| 5.7317055 |
| 0.004031 |
| 0.1071925 |
| 17.36421 |
| 0.0001573 |
| 5.3306568 |
| 0.113128 |
| 0.2116971 |
| 4.665499 |
| 0.0568863 |
| 3.4343942 |
| 3.4787237 |
| 0.2151045 |
| 0.2106491 |
| 4.533904 |
| 0.148092 |
| 13.016876 |
| 0.1872488 |
| 0.1607569 |
| 22.381761 |
| 9.3438772 |
| 0.2750103 |
| 0.2824614 |
| 0.0679149 |
| 7.0734265 |
| 0.1871996 |
| 0.3337508 |
| 0.1539306 |
| 0.3140417 |
| 4.0888133 |
| 18.743676 |
| 3.384151 |
| 24.352047 |
| 5.5088597 |
| 2.9553393 |
| 0.0060866 |
| 4.1411796 |
| 0.2420211 |
| 0.249382 |
| 3.6437744 |
| 0.2397907 |
| 0.000305 |
| 5.932515 |
| 29.077563 |
| 0.207832 |
| 4.7284498 |
| 3.2122664 |

4.6782907
7.0166679
0.148825
12.012786
0.2148213
4.4959914
0.2260358
0.3722717
2.843917
0.2769512
0.1853234
108.42343
0.3485808
0.2346913
4.5188214
101.70602
2.920775
3.9115616
0.0926657
0.3557563
3.9723836
6.6035123
0.3776422
9.2148821
0.3374902
2.6680765
8.407802
4.4111093
14.330504
3.3043909
5.3055252
1849.3954
2.7789769
0.2755112
0.3962718
0.1650812
4.0749331
0.2701314
20.270645
0.3597534
0.2717036
0.1504149
0.3589952
0.2947202
0.3915649
0.2756249
0.2336015
5.6845418
2.6857289
2.8186723
0.3309391
0.2534539
2.5337827
7.4712913
0.1675037
0.158759
1188.5241
0.1486188
0.3561052

| |
|---|
| 2.5134688 |
| 0.2883846 |
| 0.4046922 |
| 3.5432064 |
| 2.9069323 |
| 0.381927 |
| 2.635245 |
| 0.2949163 |
| 0.423353 |
| 0.4041488 |
| 0.1686544 |
| 2.6381729 |
| 3.3314475 |
| 3.6420395 |
| 2.4653102 |
| 5.5439743 |
| 0.0687414 |
| 15.536316 |
| 0.2859073 |
| 0.1793722 |
| 0.1779226 |
| 2.4090683 |
| 0.1349413 |
| 0.0009939 |
| 0.3113182 |
| 0.3791711 |
| 0.2309829 |
| 3.5097902 |
| 6.1471415 |
| 0.3019156 |
| 3.1041006 |
| 3.7978453 |
| 4.4105205 |
| 5.1192243 |
| 0.1083481 |
| 0.0772015 |
| 3.07969 |
| 0.2102317 |
| 0.1796495 |
| 2.7147388 |
| 0.3654879 |
| 0.3501982 |
| 0.297284 |
| 18.088343 |
| 0.277099 |
| 0.2399203 |
| 4.4250109 |
| 3.3741395 |
| 0.3966827 |
| 2.6199588 |
| 2.4529875 |
| 3.6721455 |
| 11.666236 |
| 0.3368592 |
| 2.2509521 |
| 4.1468805 |
| 0.4091176 |
| 0.414096 |
| 5.9018793 |

| |
|---|
| 2.4963191 |
| 3.1776835 |
| 0.3578059 |
| 4.0060639 |
| 3.4715913 |
| 0.0021369 |
| 0.001389 |
| 7.4241792 |
| 2.2078864 |
| 2.7705447 |
| 16.717159 |
| 0.3310508 |
| 0.2448915 |
| 0.2242235 |
| 0.3475406 |
| 2.8918202 |
| 4.8178536 |
| 2.7766526 |
| 3.2281129 |
| 4.9142625 |
| 0.3677867 |
| 0.002393 |
| 0.3581299 |
| 0.3507916 |
| 0.3589706 |
| 3.1140124 |
| 0.2975312 |
| 731.96652 |
| 0.2634129 |
| 3.9772094 |
| 14.136258 |
| 3.8924547 |
| 0.4111743 |
| 0.3804475 |
| 0.0015436 |
| 2.4630917 |
| 2.6018461 |
| 2.4260626 |
| 3.3550773 |
| 0.4461876 |
| 0.3877442 |
| 7.3219675 |
| 0.0961327 |
| 7.2457716 |
| 0.3176448 |
| 527.99098 |
| 0.2887291 |
| 0.3719437 |
| 2.3472286 |
| 0.101762 |
| 2.6073124 |
| 5.0242672 |
| 510.57213 |
| 0.1961951 |
| 5.3696366 |
| 2.5155927 |
| 3.4022222 |
| 0.3579669 |
| 7.1075115 |

| |
|---|
| 0.4574402 |
| 0.3885842 |
| 2.8525038 |
| 3.8427477 |
| 0.4205494 |
| 3.9362255 |
| 0.2552774 |
| 0.4155002 |
| 0.2802065 |
| 3.400752 |
| 2.0676381 |
| 0.2890207 |
| 0.4836655 |
| 0.2977087 |
| 0.460358 |
| 0.4234358 |
| 0.001837 |
| 3.3728128 |
| 6.0493283 |
| 3.0245625 |
| 3.2051625 |
| 2.2382108 |
| 59.615923 |
| 3.0294663 |
| 0.2148736 |
| 0.422088 |
| 2.188843 |
| 5.1930474 |
| 2.6595448 |
| 0.3388331 |
| 0.1154413 |
| 3.4287558 |
| 0.0028537 |
| 3.1024787 |
| 0.4436196 |
| 2.63552 |
| 2.1541427 |
| 2.1161916 |
| 0.3678031 |
| 0.1763638 |
| 0.0017062 |
| 2.074767 |
| 2.1944209 |
| 0.413686 |
| 0.3591002 |
| 3.5873857 |
| 0.4833977 |
| 406.4467 |
| 0.4508134 |
| 2.4731158 |
| 9.0620231 |
| 0.4920212 |
| 2.5686735 |
| 0.3641952 |
| 2.5724974 |
| 4.1286565 |
| 5.694826 |
| 0.3688848 |
| 3.2923488 |

| |
|---|
| 0.3069892 |
| 0.2636588 |
| 0.4764795 |
| 0.4388114 |
| 395.76283 |
| 6.5143684 |
| 2.952502 |
| 0.4387273 |
| 0.3117251 |
| 2.7305832 |
| 4.7752505 |
| 6.2453439 |
| 18.37571 |
| 3.4621014 |
| 0.3016043 |
| 0.4348411 |
| 0.0025722 |
| 0.4793234 |
| 0.4270968 |
| 0.4075277 |
| 0.3367149 |
| 0.4778568 |
| 0.3619672 |
| 0.3313949 |
| 2.2242624 |
| 3.6913315 |
| 0.207221 |
| 2.4885481 |
| 0.477024 |
| 0.4125034 |
| 2.0280478 |
| 2.6961033 |
| 0.3144721 |
| 2.0722127 |
| 3.9414664 |
| 5.6906026 |
| 364.89468 |
| 2.5332728 |
| 3.5825751 |
| 6.6069489 |
| 2.9692692 |
| 0.1590282 |
| 0.3204419 |
| 2.4188961 |
| 3.2818946 |
| 2.5147939 |
| 0.4539052 |
| 0.0134507 |
| 0.2202407 |
| 0.4353802 |
| 2.0099011 |
| 2.6551079 |
| 0.3271902 |
| 0.4566242 |
| 0.3602653 |
| 2.5766644 |
| 0.4577456 |
| 0.4517255 |
| 2.2732139 |

| |
|---|
| 0.353414 |
| 2.0605582 |
| 9.4836106 |
| 0.022313 |
| 0.3950294 |
| 3.194823 |
| 2.6019648 |
| 3.7730475 |
| 0.1214159 |
| 2.2460498 |
| 0.4386521 |
| 0.4592251 |
| 2.0507736 |
| 4.0879576 |
| 0.0031245 |
| 0.4837351 |
| 0.3880062 |
| 307.91183 |
| 0.4188767 |
| 0.0023192 |
| 0.4492213 |
| 0.3598016 |
| 2.4315724 |
| 3.8267031 |
| 0.4803962 |
| 2.2576237 |
| 3.4143741 |
| 0.1611441 |
| 0.3826533 |
| 299.67547 |
| 0.2379622 |
| 2.1560158 |
| 3.8333269 |
| 2.4361305 |
| 0.4066655 |
| 2.1518575 |
| 2.1270907 |
| 0.2189881 |
| 0.1539818 |
| 3.3670788 |
| 0.4861401 |
| 2.4497762 |
| 0.2430549 |
| 282.7136 |
| 0.0623831 |
| 3.3961182 |
| 0.4797471 |
| 0.3549233 |
| 155.17099 |
| 0.4641007 |
| 0.4354221 |
| 2.9146485 |
| 0.0887245 |
| 3.7074276 |
| 0.3294496 |
| 4.1637023 |
| 5.3500918 |
| 0.2449923 |
| 3.4773429 |

| |
|---|
| 2.9244934 |
| 10.643096 |
| 4.2921307 |
| 0.3994416 |
| 0.350793 |
| 3.2260336 |
| 2.6776631 |
| 0.2577 |
| 2.1070985 |
| 2.6344872 |
| 7.7058017 |
| 0.4065472 |
| 0.4967785 |
| 2.1754267 |
| 0.0445091 |
| 2.1266182 |
| 5779414.5 |
| 0.4589907 |
| 0.4674428 |
| 0.3983068 |
| 0.3714958 |
| 2.0746724 |
| 0.3729039 |
| 9.8692276 |
| 2.4343973 |
| 0.4459025 |
| 0.4412257 |
| 0.0034245 |
| 0.3674695 |
| 10.592753 |
| 256.39135 |
| 0.2773309 |
| 0.4665043 |
| 247.27868 |
| 250.88585 |
| 0.1362413 |
| 0.3757335 |
| 3.5109557 |
| 0.0038642 |
| 7.9463742 |
| 3.102079 |
| 2.5416135 |
| 0.2542775 |
| 0.4659866 |
| 0.4015603 |
| 3.2022998 |
| 2.1119372 |
| 3.1818109 |
| 245.56554 |
| 4.1296208 |
| 2.2888244 |
| 0.0961233 |
| 0.3928763 |
| 2.9665237 |
| 2.908353 |
| 2.6991265 |
| 2.8916913 |
| 2.6626852 |
| 0.4130599 |

| |
|---|
| 0.2719934 |
| 0.4497002 |
| 3.0518213 |
| 0.1948078 |
| 0.1435428 |
| 0.459883 |
| 18.124976 |
| 0.1130969 |
| 0.3636765 |
| 2.0983637 |
| 2.9833139 |
| 2.2042216 |
| 3.2284027 |
| 0.4448064 |
| 0.1401712 |
| 7.0935175 |
| 0.3667307 |
| 0.2784695 |
| 0.3829036 |
| 3.8662532 |
| 2.1153947 |
| 2.354259 |
| 2.2601936 |
| 0.4121046 |
| 0.0040058 |
| 2.7174349 |
| 0.4475853 |
| 0.0340376 |
| 0.4572306 |
| 0.2736806 |
| 3.3632856 |
| 0.3834643 |
| 5.1649197 |
| 0.2437986 |
| 0.2250383 |
| 2.6339577 |
| 0.2777634 |
| 0.1217969 |
| 0.464891 |
| 0.3558351 |
| 0.001476 |
| 0.4860735 |
| 0.3159787 |
| 0.4446302 |
| 4.2278627 |
| 0.4339851 |
| 0.0046138 |
| 0.1677079 |
| 2.8140885 |
| 0.2688099 |
| 3.0189417 |
| 5.627942 |
| 2.2226476 |
| 0.4841046 |
| 23.70281 |
| 0.4714903 |
| 3.6477823 |
| 0.3408405 |
| 0.320777 |

| |
|---|
| 0.4794058 |
| 2.3450659 |
| 0.3752074 |
| 2.6045217 |
| 0.1942649 |
| 2.2953594 |
| 5.6432873 |
| 2.5389205 |
| 0.1028319 |
| 2.2241094 |
| 0.4968769 |
| 0.3451292 |
| 2.5121538 |
| 4.3130309 |
| 0.3509975 |
| 8.2537308 |
| 0.0275628 |
| 2.7272646 |
| 3.434038 |
| 2.6843715 |
| 0.420286 |
| 0.1535003 |
| 3.0897201 |
| 0.1446448 |
| 2.233353 |
| 0.2934966 |
| 0.004072 |
| 5.0536417 |
| 10.078481 |
| 0.4930304 |
| 0.184052 |
| 2.4262739 |
| 0.3689274 |
| 0.4613079 |
| 3.0531462 |
| 6.2498774 |
| 0.4613831 |
| 0.4734644 |
| 5.0069068 |
| 0.4084985 |
| 204.44628 |
| 0.0813805 |
| 0.364172 |
| 2.1819673 |
| 186.30855 |
| 508.32037 |
| 3.6907387 |
| 4.355739 |
| 0.1249451 |
| 0.4683671 |
| 0.3963858 |
| 2.6607826 |
| 0.2205537 |
| 178.29698 |
| 3.3719313 |
| 0.2847866 |
| 0.3825116 |
| 185.82412 |
| 0.2445197 |

| |
|---|
| 28.562737 |
| 0.4712082 |
| 19.792763 |
| 0.4652134 |
| 0.4739999 |
| 0.3246637 |
| 0.27706 |
| 2.9625435 |
| 4.8113305 |
| 3.1727753 |
| 0.4740356 |
| 0.2991737 |
| 174.09031 |
| 2.20959 |
| 0.0051454 |
| 2.4748053 |
| 3.4156586 |
| 2.6714313 |
| 4.1301329 |
| 0.4984366 |
| 171.39919 |
| 2.2226891 |
| 193.16701 |
| 2.1978293 |
| 0.4878017 |
| 7.0972889 |
| 0.1397637 |
| 2.4084212 |
| 0.4620425 |
| 0.0210697 |
| 0.0718364 |
| 0.4226505 |
| 0.3224987 |
| 2.651071 |
| 2.4370793 |
| 2.4399265 |
| 0.3072347 |
| 0.3249135 |
| 0.2367096 |
| 0.4878173 |
| 2.2601136 |
| 2.1707466 |
| 0.453425 |
| 2.5044653 |
| 0.4854645 |
| 2.6153591 |
| 0.4665127 |
| 2.6145643 |
| 0.4278406 |
| 3.335602 |
| 0.4320825 |
| 29.299547 |
| 0.1215807 |
| 0.292162 |
| 3.4384689 |
| 2.8241768 |
| 0.0716279 |
| 7.588793 |
| 2.353234 |

| |
|---|
| 0.443768 |
| 0.4381984 |
| 10.774836 |
| 0.3649943 |
| 0.3800293 |
| 0.4598262 |
| 0.4623797 |
| 0.3084507 |
| 0.4446146 |
| 0.216559 |
| 2.2501542 |
| 0.4279481 |
| 2.787329 |
| 0.4060835 |
| 5.9538794 |
| 0.3572898 |
| 3.344333 |
| 0.0206615 |
| 6.8147373 |
| 2.1276828 |
| 0.023846 |
| 3.0124278 |
| 0.0026742 |
| 150.66168 |
| 16.259512 |
| 3.9733919 |
| 2.1341454 |
| 7.4199003 |
| 0.0052489 |
| 3.05656 |
| 0.4472799 |
| 0.4320461 |
| 4.1807018 |
| 2.2302945 |
| 3.2017291 |
| 0.1641303 |
| 0.0567133 |
| 0.47888 |
| 2.0748034 |
| 3.2049525 |
| 0.3725688 |
| 3.6886008 |
| 3.8767183 |
| 4.1666968 |
| 0.004683 |
| 3.021163 |
| 2.6371985 |
| 3.8414603 |
| 2.0314561 |
| 0.0067242 |
| 0.0058275 |
| 0.006792 |
| 0.4171104 |
| 2.2762733 |
| 0.2147055 |
| 13.25743 |
| 0.4606266 |
| 5.3074643 |
| 2.6405792 |

| |
|---|
| 0.308203 |
| 145.73146 |
| 136.44501 |
| 0.0059212 |
| 2.0296794 |
| 0.4028692 |
| 0.2271605 |
| 3.0489161 |
| 0.3153803 |
| 0.4073271 |
| 0.0015726 |
| 0.3465216 |
| 2.305182 |
| 0.268543 |
| 0.3566504 |
| 0.4702323 |
| 2.4288232 |
| 2.2659899 |
| 0.4561593 |
| 4.1420581 |
| 0.4801031 |
| 0.2897124 |
| 0.4888947 |
| 0.4555288 |
| 2.2042288 |
| 2.3881267 |
| 2.2579262 |
| 4.2996974 |
| 0.4946222 |
| 0.3498421 |
| 2.1397445 |
| 22.157864 |
| 2.1161898 |
| 0.0057948 |
| 2.0356607 |
| 0.4445604 |
| 0.1147703 |
| 2.1260152 |
| 0.1028649 |
| 36.915603 |
| 0.0140716 |
| 0.4158954 |
| 0.4201592 |
| 0.4231023 |
| 9.6658512 |
| 3.0314004 |
| 0.4945988 |
| 4.6304997 |
| 0.436124 |
| 2.6067987 |
| 2.6277313 |
| 2.2299455 |
| 2.0331357 |
| 3.562994 |
| 6.7531691 |
| 2.2324327 |
| 0.2839492 |
| 0.2254282 |
| 0.2254282 |

| |
|---|
| 0.032567 |
| 5.3501261 |
| 2.011663 |
| 0.2630521 |
| 7.2663838 |
| 2.1288026 |
| 2.3160921 |
| 2.16751 |
| 2.483885 |
| 2.7824637 |
| 3.3262698 |
| 0.2133989 |
| 0.4250843 |
| 2.5714929 |
| 122.44093 |
| 0.0077355 |
| 0.0031759 |
| 2.1617929 |
| 0.1210134 |
| 0.4101812 |
| 0.4994406 |
| 6.3849383 |
| 0.3973579 |
| 123.44042 |
| 2.7477151 |
| 0.0079407 |
| 0.3196547 |
| 6.0133551 |
| 0.0072795 |
| 2.3144166 |
| 10.04548 |
| 0.262307 |
| 0.1725632 |
| 123.92862 |
| 0.415496 |
| 0.4975567 |
| 2.8819776 |
| 0.4621797 |
| 0.3081136 |
| 2.8357059 |
| 2.3041042 |
| 2.3413387 |
| 2.1842884 |
| 2.2753737 |
